# Physiological basis underlying antidepressant-induced activation of TrkB receptors

**DOI:** 10.1101/2021.08.30.458151

**Authors:** Okko Alitalo, Samuel Kohtala, Marko Rosenholm, Piia Kohtala, Roosa Saarreharju, Nobuaki Matsui, Gemma Gonzalez Hernandez, Heidi Kaastrup Müller, Wiebke Theilmann, Mirkka Sarparanta, Anders Klein, Olli Kärkkäinen, Stanislav Rozov, Tomi Rantamäki

**Author notes:** equal contribution (Marko Rosenholm, Piia Kohtala, Roosa Saarreharju, Nobuaki Matsui).

## Abstract

We show that both pharmacological and non-pharmacological treatments of depression activate TrkB receptors—a well-established target of antidepressants—by inducing a physiological response coupled to sedation. Several rapid-acting antidepressants trigger TrkB signaling by evoking a state associated with electroencephalographic slow-wave activity, behavioral immobility, reduced cerebral glucose utilization, and lowered body temperature. Remarkably, antidepressant-induced TrkB signaling was not compromised in animals exhibiting reduced activity-dependent release of BDNF but was diminished by maintaining animals in warm ambient temperature. Most importantly, prevention of the hypothermic response attenuated the behavioral effects produced by rapid-acting antidepressant nitrous oxide. Our results suggest that the phenomenon underlying TrkB transactivation—changes in energy expenditure and thermoregulation—is essential, but not sufficient, for antidepressant responses. Indeed, regardless of differential clinical and pharmacodynamic properties, all drugs that disrupt energy metabolism and induce hypothermia activated TrkB. This study challenges pharmacology-centric hypotheses regarding antidepressant effects and highlight the role of complex changes in bioenergetics and thermoregulation.

**Highlights:**

- Rapid-acting antidepressants evoke homeostatic emergence of slow-wave sleep during which TrkB signaling becomes regulated.
- Non-antidepressant metabolic inhibitors and diverse sedatives activate TrkB signaling.
- Reduction in body temperature determined the ability of antidepressants to transactivate TrkB.
- Drug-induced TrkB signaling was blunted by maintenance of normothermic body temperature.
- Warm ambient temperature after nitrous oxide exposure blocked the antidepressant-like effects.

**Graphical abstract:** 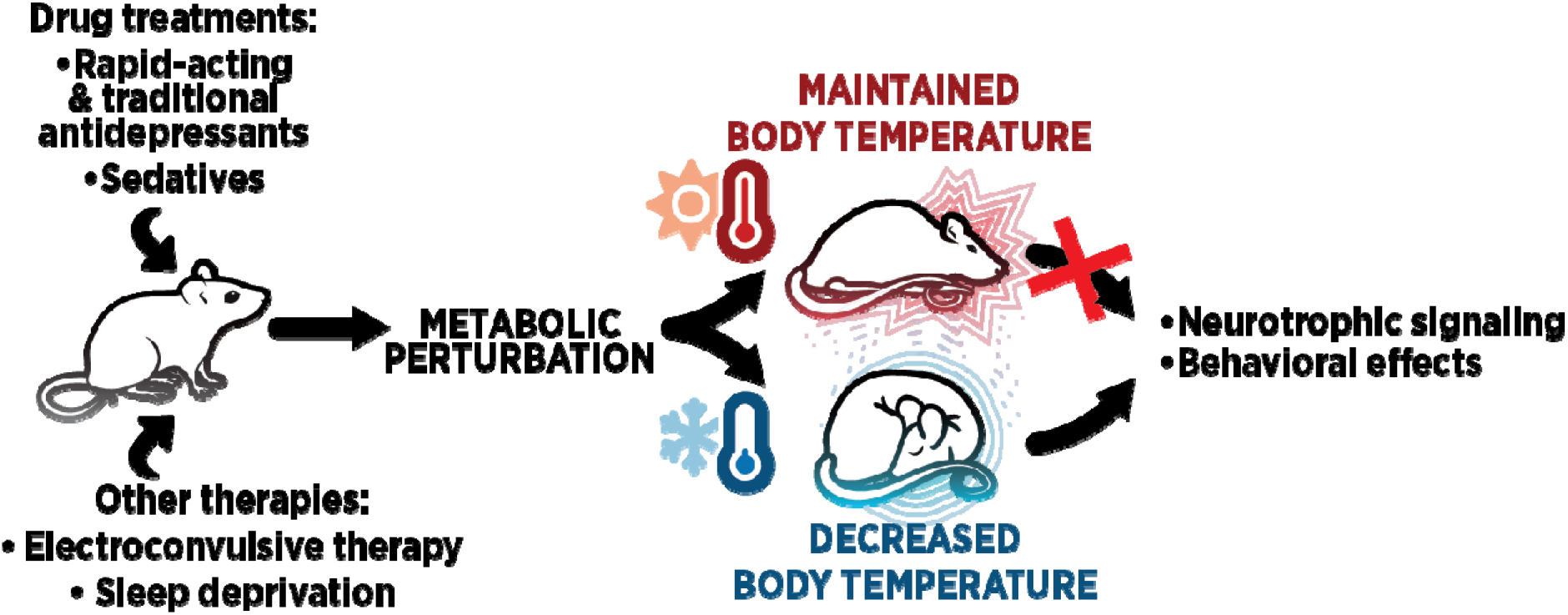

## Introduction

Accumulating evidence suggests that antidepressant drugs act by promoting synaptic plasticity in the adult brain by facilitating the signaling of tropomyosin-related kinase B (TrkB)—the receptor of brain-derived neurotrophic factor (BDNF) (Duman et al., 2016). Pharmacologically diverse antidepressants, including tricyclic antidepressants (e.g., amitriptyline), selective serotonin reuptake inhibitors (e.g., fluoxetine), and rapid-acting antidepressants (e.g., ketamine; *N*-methyl-D-aspartate receptor [NMDAR] antagonist), converge in activation of TrkB and its downstream signaling cascades, which include activation of mammalian target of rapamycin (mTOR) and inhibition of glycogen synthase kinase 3β (GSK3β) in the adult rodent brain (Autry et al., 2011; Li et al., 2010; Rantamäki et al., 2007; Saarelainen et al., 2003).

Several mechanisms have been proposed to underlie antidepressant-induced TrkB activation. Initially, chronic administration of conventional antidepressants was shown to enhance BDNF expression in several brain areas (Nibuya et al., 1995). Subsequent research suggested that ketamine induces rapid BDNF translation and release (Autry et al., 2011; Lepack et al., 2014). Some drugs like amitriptyline and 7,8-dihydroxyflavone have been proposed to act as direct agonists of TrkB (Jang et al., 2009, 2010), however there have been calls to reevaluate whether these drugs actually bind to the receptor *in vivo and* whether the potential interaction with the receptor triggers downstream signaling (Boltaev et al., 2017; Pankiewicz et al., 2021; Rantamäki et al., 2011). More recently, antidepressant drugs have been suggested to facilitate the neuronal responsiveness to BDNF by binding to a transmembrane site of TrkB (Casarotto et al., 2021). Casarotto and colleagues propose that rapid-acting antidepressants bind to TrkB more readily than traditional antidepressants, which require chronic administration and accumulation in the brain tissue to reach concentrations that promote TrkB signaling. However, earlier findings indicate that conventional antidepressants activate brain TrkB receptors very rapidly (Rantamäki et al., 2007; Saarelainen et al., 2003)—within an hour—and without the involvement of BDNF (Rantamäki et al., 2011). Furthermore, the effects of some clinically effective antidepressants are difficult to explain using the principles of classic receptor pharmacology. For example, the antidepressant effects of nitrous oxide (N_2_O; laughing gas) (Nagele et al., 2021) and flurothyl —“a chemical electroconvulsive therapy (ECT) (Krantz et al., 1957)”— become apparent long after their exposure. Notably, our recent findings indicate that these drugs activate TrkB signaling after their acute pharmacological effects have dissipated (Kohtala et al., 2019a). Although both flurothyl and N_2_O are eliminated from the body within minutes, TrkB becomes and remains phosphorylated long afterwards (Kohtala et al., 2019a), suggesting that they render TrkB active using a mechanism that does not involve continuous ligand release and receptor binding.

TrkB activation induced by flurothyl and N_2_O temporally coincides with the emergence of electroencephalographic (EEG) slow-wave activity (SWA) (Kohtala et al., 2019a), a neurophysiological signature of deep sleep (Tononi and Cirelli, 2014) and a predictive biomarker for the therapeutic efficacy of ECT (Folkerts, 1996; Sackeim et al., 1996). Notably, sleep deprivation (SD)—another non-pharmacological antidepressant (Ioannou et al., 2021)—also triggers homeostatic emergence of SWA. Recent studies have also associated the therapeutic effects of ketamine with augmentation of SWA and increased BDNF levels during subsequent sleep after administration, illustrating the association of reduced electrophysiological activity and neurotrophic signaling (Duncan et al., 2013; Kohtala et al., 2020; Rantamäki and Kohtala, 2020). We have observed in rodents that subanesthetic ketamine, analogous to N_2_O, flurothyl, and SD, evokes SWA after its acute effects on high-frequency gamma activity fade out (Kohtala et al., 2019a), similar to earlier findings on NMDAR antagonists (Campbell and Feinberg, 1996). Furthermore, there exists a strong association between EEG slowing and the phosphorylation of TrkB. Specifically, the propensity of several anesthetic agents, including putative rapid-acting antidepressant isoflurane (Langer et al., 1995), to activate TrkB is associated to their potency of suppressing the EEG signal (Antila et al., 2017; Theilmann et al., 2019). Notably, these agents also include ketamine (Kohtala et al., 2019b), even though only subanesthetic doses are considered to induce clinical antidepressant effects. It should be noted that several monoaminergic antidepressants are characterized by sedation—often induced through antihistaminergic activity—as their most prominent side effect. Collectively, these results suggest that TrkB signaling is triggered during a sedative or sleep-like physiological state evoked in response to the antidepressant treatment. Here, we provide strong evidence to support this hypothesis and reveal that disrupted bioenergetics and accompanied hypothermia—rather than a specific receptor-level effect—underlie the TrkB activation induced by various types of antidepressants. Furthermore, we show that similar activation of TrkB is induced by a wide range of sedative agents with no documented antidepressant efficacy. Our findings therefore suggest that the phenomenon underlying TrkB transactivation—changes in energy expenditure and thermoregulation—is essential, but not sufficient as such, for antidepressant responses.

## Results

### Physiological adaptations associated with N_2_O-induced TrkB signaling

N_2_O evokes TrkB signaling after its acute pharmacological effects have dissipated and this co-associates with the emergence of SWA (1–4 Hz band EEG, delta) (Kohtala et al., 2019a), one of the typical physiological features of deep slow-wave sleep (SWS). Consequently, we sought to investigate whether N_2_O regulates also other physiological changes connected with SWS: behavioral immobility (Szymusiak, 2018), attenuated brain energy expenditure (DiNuzzo and Nedergaard, 2017), and reduced body temperature (Landolt et al., 1995). α_2_-adrenergic sedative drug medetomidine was used as a positive control due to its ability to produce sleep-like state and TrkB signaling (Kohtala et al., 2019a; Purdon et al., 2015).

First, we replicated and extended our earlier observation that TrkB signaling is gradually activated only during withdrawal from N_2_O (65%, 20 min) when EEG activity is predominated by SWA (**Figure 1A-B**). This state also coincided with locomotor inactivity in the open field test (**Figure 1C**). The animals were observed to be noticeably sedated, remaining still in a crouched, sleep-like posture throughout the monitoring period. Animals under the influence of N_2_O were slightly more active than control animals. However, medetomidine elicited rapid and sustained sedation along with significant activation of TrkB signaling in the medial prefrontal cortex (mPFC; **Figure S1A**). A similar increase in the phosphorylation of TrkB^Y816^, GSK3β^S9^, and p70SK6^T421/S424^ was observed in both total mPFC homogenate and isolated crude synaptosomal fraction, suggesting that the effect is not restricted solely to the synaptic terminals in the mPFC. Furthermore, the effect was also not localized only to the cortex but occurred also in the hippocampus **(Figure S1B)**.

**Figure 1.**
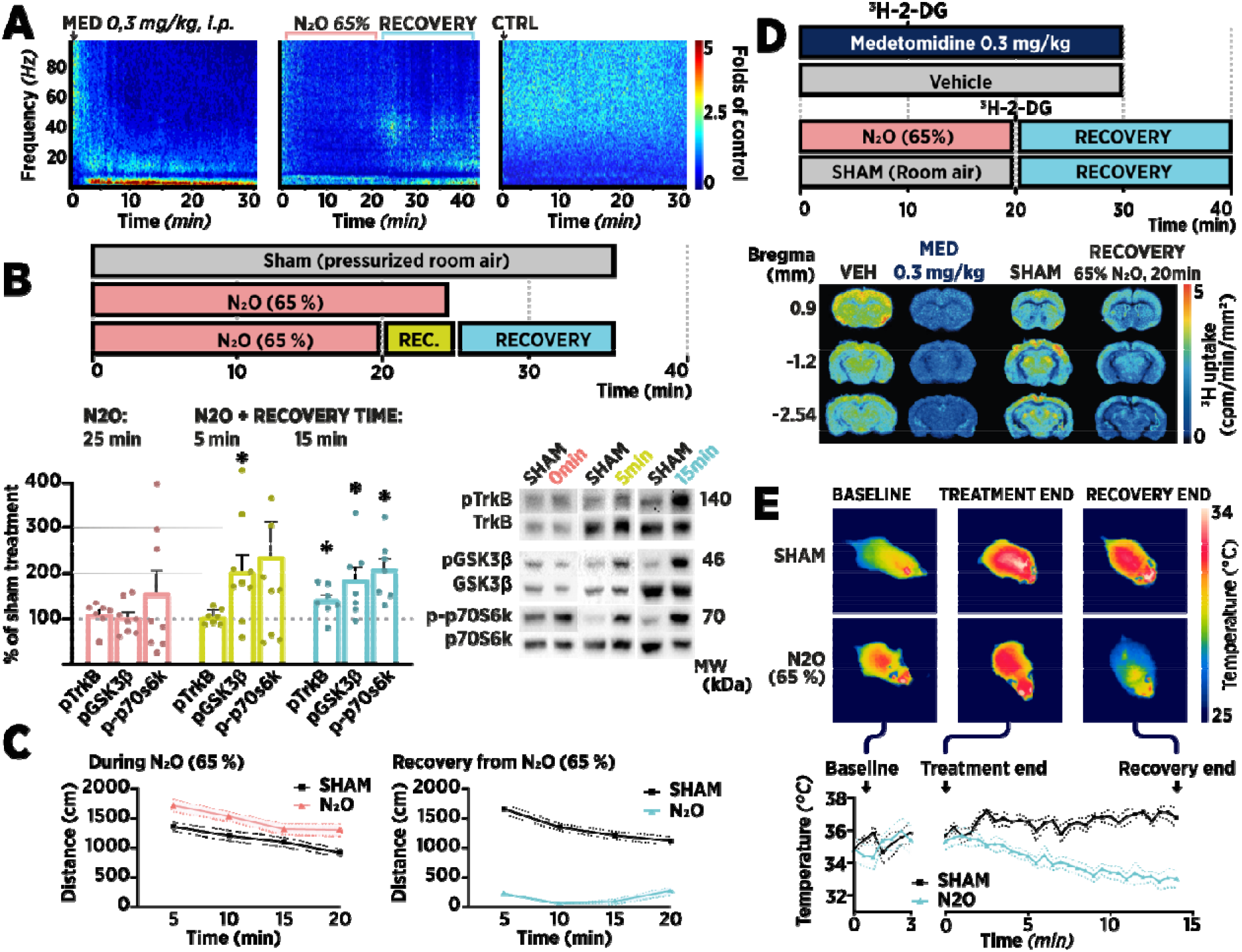
N_2_O evokes a sleep-like state, during which TrkB signaling becomes regulated. **A**. Medetomidine (MED; 0.3 mg/kg, intraperitoneal [i.p.]) produces a robust and rapid increase in SWA, whereas nitrous oxide (N_2_O; 65%, 20 min) administration triggers SWA gradually after exposure. Spectrograms show the difference in spectral power normalized to baseline. Control animals received i.p. injection of vehicle (VEH). **B**. Phosphorylation levels of TrkB^Y816^, GSK3β^S9^, and p70SK6^T421/S424^ increase gradually after N_2_O (65%, 20 min) administration is terminated, reaching significant levels by 15 min. Animals terminated at 25 min of continuous N_2_O exposure showed negligible effects on signaling. **C**. Locomotor activity in the open field during and after N_2_O exposure (65%, 20 min). **D**. ^3^H-deoxy-D-glucose (^3^H-2-DG) autoradiography reveals reduced glucose utilization after N_2_O exposure (65%, 20 min). Glucose utilization during medetomidine (0.3 mg/kg, i.p.) sedation shows more pronounced reduction. **E**. Whole-body thermal imaging reveals reduced body temperature upon withdrawal from N_2_O exposure. Qualitatively, the temperature of distal body parts decreased more markedly during recovery. Increased heat was detected in proximity to brown adipose tissue. Phosphoproteins were normalized against corresponding total protein and compared to the control group set to 100%. Data are presented as mean ± standard error of mean *≤0.05 (for statistical analyses and *n* numbers see **Table S1**). Abbreviations: GSK3β, glycogen synthase kinase 3β; p70S6k, ribosomal protein S6 kinase.

To investigate the potential bioenergetic resemblance between the state evoked by N_2_O and state induced during sleep, β-emission camera-based digital ^3^H-2-deoxyglucose (^3^H-2DG) autoradiography was used (Alitalo et al., 2020; Sokoloff et al., 1977). Comparable methods have been used to assess the metabolic demand during different sleep stages and acute drug effects (Miyamoto et al., 2000; Vyazovskiy et al., 2008) as they facilitate functional brain imaging without the use of anesthesia, which is a significant confounding factor. Uptake of ^3^H-2DG by cells occurs in a similar manner as glucose uptake, but 2DG is unable to undergo glycolytic metabolism and thus traps the radiolabel. The resulting autoradiograph of label distribution in the brain sections thus signifies the regional glucose utilization (i.e., functional activity) during radiolabel uptake time. To assess cerebral metabolism during recovery from N_2_O exposure, we injected animals with ^3^H-2DG immediately after 20 min of 65% N_2_O exposure or sham treatment and terminated them after another 20 min of radiolabel uptake. Animals in a separate cohort were administered medetomidine (0.3 mg/kg) or saline, injected with ^3^H-2DG 10 min afterwards when the sedative effects reached a plateau, and terminated after 20 min. N_2_O-exposed animals displayed a general reduction in neuronal glucose uptake during the recovery, indicating lower energy expenditure (**Figure 1D, Supplementary File A**). The N_2_O-evoked reduction in glucose utilization was subtler than that induced by medetomidine, analogous to the intensity of the electrophysiological and molecular signaling changes produced by the treatments. Animals treated with medetomidine—but not N_2_O—demonstrated significant hyperglycemia, alluding to systemic bioenergetic effects of the anesthetic (**Figure S2A**). Full images of the autoradiographic acquisition plates, results of semiquantitative analysis of regional glucose utilization, and quantitative plasma analysis of ^3^H uptake are presented in **Supplementary File A**.

Next, we assessed the effects of N_2_O on body temperature using an infrared camera, which enables remote temperature monitoring in distinct body parts of freely moving animals without inflicting stress. Analyses revealed a significant decrease in the cutaneous body temperature of 65% N_2_O-exposed animals during recovery (**Figure 2E**). Similar to the gradual activation of TrkB signaling, a progressive reduction in body temperature was observed throughout the monitoring period. Overall, these findings demonstrate that brief exposure to N_2_O evokes the activation of TrkB signaling during the emergence of a state that exhibits various key characteristics of deep sleep. Notably, this state develops only after the acute pharmacodynamic effects of N_2_O have subsided and the compound itself has been metabolized from the system. These findings suggest that the observed TrkB signaling is connected to the physiological response produced by the treatment, as opposed to direct pharmacodynamic effect.

**Figure 2.**
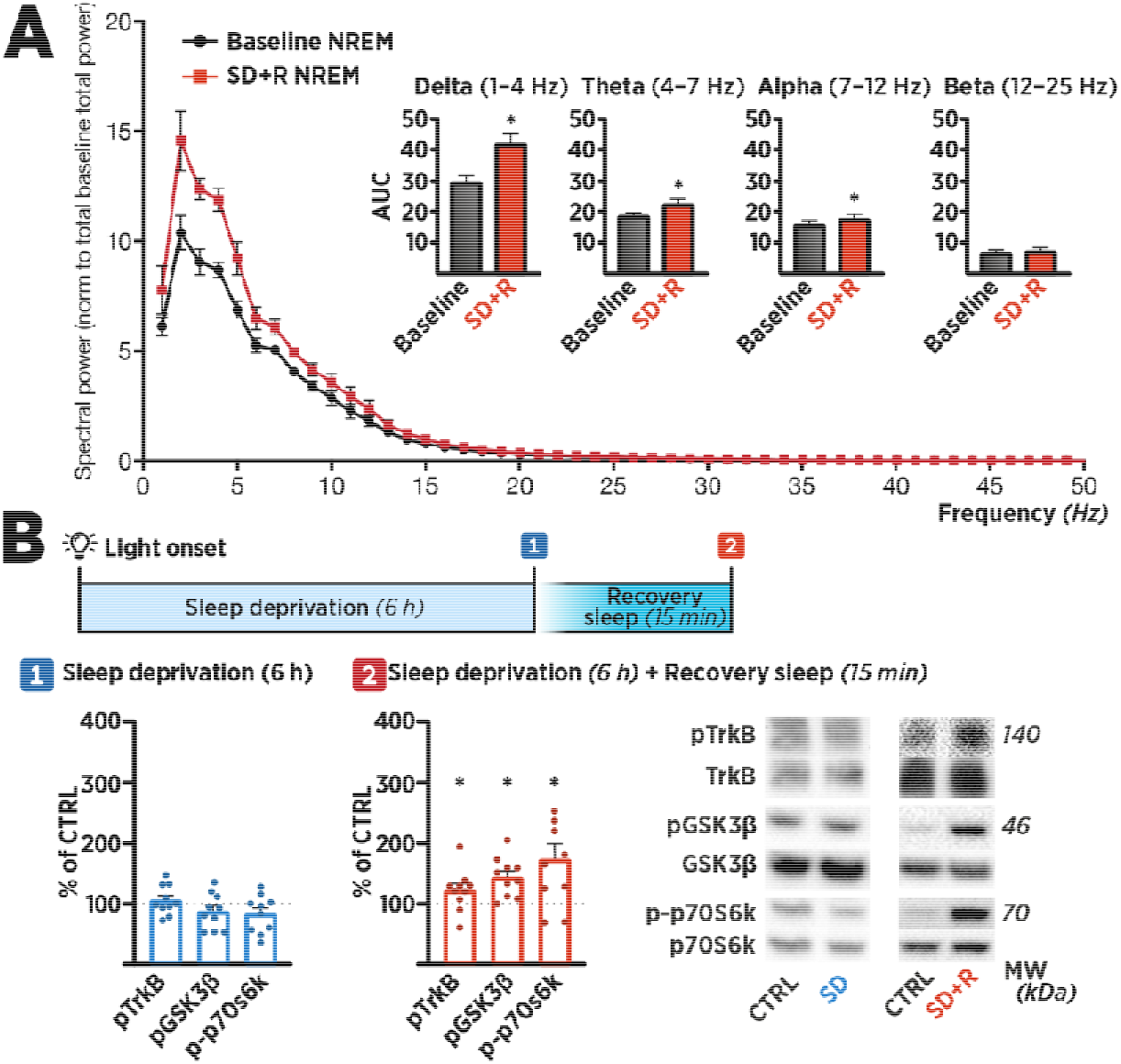
Sleep deprivation triggers TrkB signaling during homeostatically increased slow-wave sleep. **A**. Slow-wave activity within the delta band (1–4 Hz) increased significantly during non-rapid eye movement (NREM) sleep of 1.5 h of recovery sleep (SD+R) following 6 h of sleep deprivation (SD). Baseline NREM represents the spectral power of NREM sleep over 24 h before SD. **B**. Medial prefrontal cortex samples were collected immediately after 6 h of SD or 15 min after the animals have entered recovery sleep based on their immobility and posture. Phosphorylation of TrkB^Y816^, GSK3β^S9^, and p70SK6^T421/S424^ remained unaltered at the end of SD, whereas an increase was observed in the samples obtained from animals in recovery sleep (SD+R). Representative blot images with corresponding molecular weights (MW) are shown on the right. Phosphoproteins were normalized against the total protein signal and compared to the control group set to 100%. Data are presented as mean ± standard error of mean. *≤0.05 (for statistical analyses and *n* numbers see **Table S1**). Abbreviations: GSK3β, glycogen synthase kinase 3β; p70S6k, ribosomal protein S6 kinase; SD-R, sleep deprivation + recovery.

### Sleep deprivation triggers TrkB signaling during homeostatic slow-wave sleep rebound

SD produces rapid antidepressant effects in a subset of patients (Ioannou et al., 2021). Notably, during SD, there is a progressive increase in neuronal excitability and energy demand, which accumulates sleep pressure (Jung et al., 2011). When SD ceases, the subject enters recovery sleep, which is electrophysiologically “deeper” than ordinary sleep. Thus, at first glance, the course of the intervention parallels that of brief flurothyl or N_2_O exposure discussed above—a transient period of excitation followed by homeostatic sedation/sleep. To test whether similar TrkB signaling response occurs during SD-induced recovery sleep, we subjected animals to 6 h of SD starting at the beginning of light period (Zeitgeber time 0 [ZT0]), after which a subset of animals was allowed to enter recovery sleep, which was assessed by behavioral immobility and posture. Recovery sleep after SD was found to increase the spectral power of the slowest EEG bands, most notably in the SWA range (~1–4 Hz; **Figure 2A**). Samples from animals terminated at the end of the SD period demonstrated unaffected phosphorylation of TrkB^Y816^, GSK3β^S9^, and p70SK6^T421/S424^ in comparison with control. However, similar to findings after N_2_O treatment, upregulated neurotrophic signaling was observed in mice that were allowed to enter recovery sleep for 15 min (**Figure 2B**).

### Sedation, reduced energy expenditure, and TrkB signaling

To dissect the role of SWA in drug-induced TrkB signaling, mice were injected with atropine, the muscarinic acetylcholine receptor antagonist; then, brain tissue samples for western blot analysis were collected 30 min afterwards. While atropine induces pronounced cortical SWA (**Figure S3A**), it is accompanied by subcortical arousal and relatively normal behavior in general (Qiu et al., 2015). Despite a strong association of SWA and TrkB signaling with medetomidine, N_2_O, and sleep, phosphorylation levels of TrkB^Y816^, GSK3β^S9^, and p70SK6^T421/S424^ remained unaltered in the mPFC samples from atropine-treated mice (**Figure S3B**), indicating that the increase in SWA does not sufficiently explain the observed alterations in TrkB signaling.

If the drug-induced TrkB signaling was connected to the physiological sedative response instead of pharmacodynamic action, identical signaling should be observed in animals undergoing sedation irrespective of the cause leading to it. Thus, we next tested the ability of a wide array of pharmacological agents to produce TrkB signaling. Mice were subjected to sedative and non-sedative compounds—with or without clinically demonstrated antidepressant properties—and brain samples were collected for western blot analysis at 30 min after treatment (**Figure 3**). Indeed, urethane (NMDA receptor/GABA_A_ receptor modulator), hydroxyzine (H_1_ receptor inverse agonist), sevoflurane, isoflurane, gamma-hydroxybutyrate (GHB; GABA_B_ receptor agonist), and sedative antipsychotics such as clozapine and chlorpromazine significantly activated TrkB signaling. Notably, several antidepressants (amitriptyline, mianserin, fluoxetine, and mirtazapine) also increased TrkB signaling. Antidepressants that were not observed to induce sedation (paroxetine and duloxetine) or stimulants such as amphetamine and atipamezole (α_2_-adrenergic antagonist) had no acute effect on TrkB signaling at the selected doses and time-point. These findings show that numerous agents activate TrkB signaling irrespective of their clinical antidepressant efficacy, but rather, their propensity to induce sedation.

**Figure 3.**
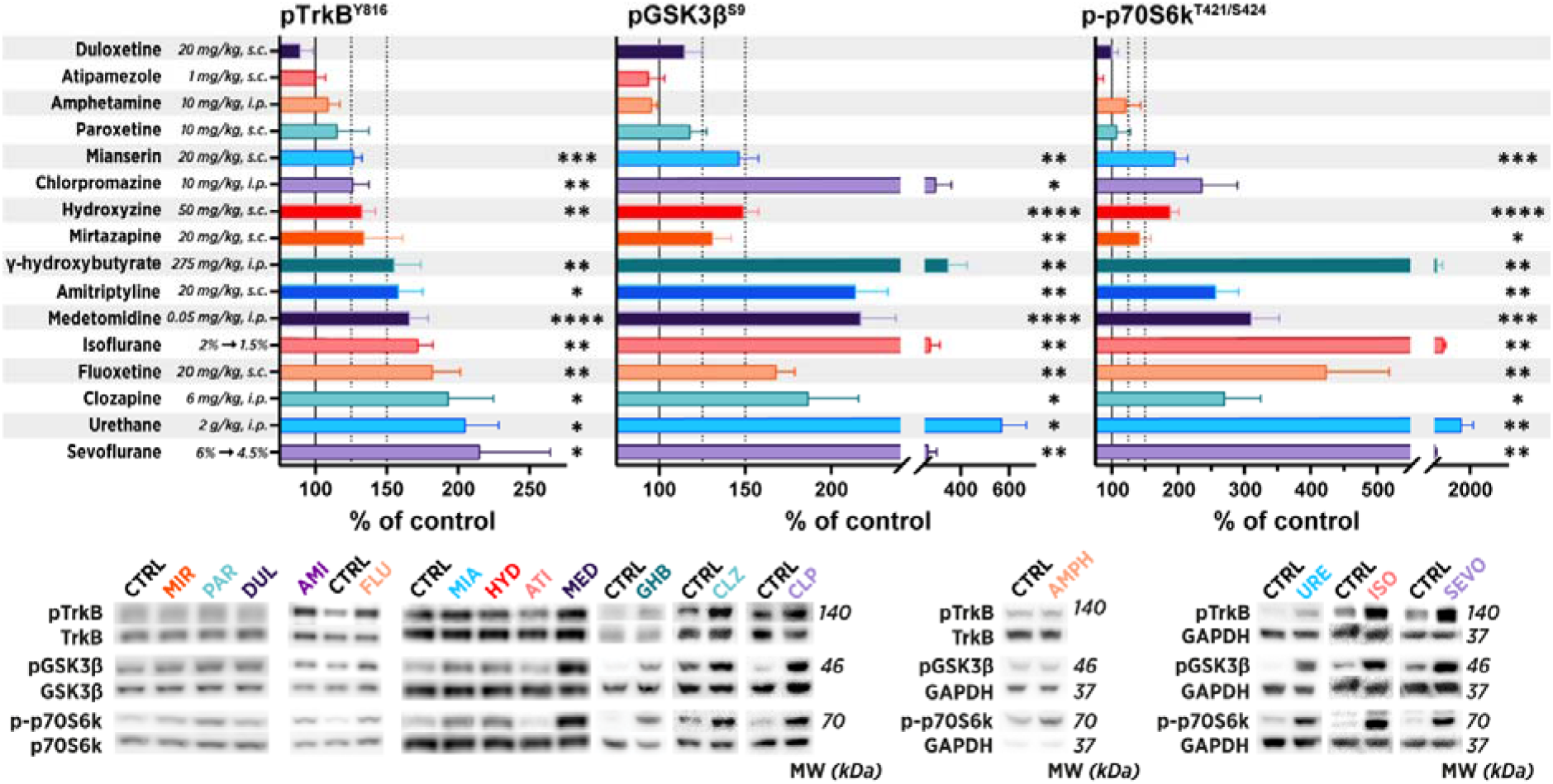
Pharmacological manipulation of TrkB signaling in the adult brain. Diverse sedative-anesthetic agents increased the phosphorylation of TrkB^Y816^, GSK3β^S9^, and p70SK6^T421/S424^ in the medial prefrontal cortex of adult mice, whereas non-sedative antidepressants such as paroxetine and stimulant drugs such as amphetamine and atipamezole showed no effects. Phosphoproteins were normalized against corresponding total protein signal and compared to the saline group set to 100%. Control animals for injected pharmacological agents were injected with a vehicle using equal volume and identical route of administration. Control animals for isoflurane and sevoflurane were subjected to pressurized room air. Data are presented as mean ± standard error of mean. *<0.05, **<0.01, ***<0.005 (for statistical analyses and *n* numbers see **Table S1**). Abbreviations: GSK3β, glycogen synthase kinase 3β; p70S6k, ribosomal protein S6 kinase.

Next, we set to find out the physiological mechanisms that could facilitate TrkB signaling observed during sedation. Flurothyl is analogous to ECT as it produces transient excitation through epileptiform activity, evoking a subsequent postictal state with significant EEG slowing. Similar to N_2_O, which does not regulate TrkB signaling during the ongoing exposure; TrkB signaling was observed to be unaffected in brain samples collected immediately after the flurothyl-induced seizure (**Figure 4A**). However, when the samples were collected an hour after the seizure, significant TrkB signaling was observed, which resembled the signaling observed under continuous anesthesia (**Figure 4A**). This finding prompted us to investigate the similarities between the physiological mechanisms underlying the recovery from a seizure and ongoing anesthesia using a non-targeted metabolomics screen (**Figure 4B; Supplementary File B)**. Overall, in comparison with recovery from a seizure, isoflurane anesthesia produced more prominent and often opposing effects in the metabolome. However, both treatments produced remarkably similar effects on bioenergetic targets involved in adenosine triphosphate (ATP) breakdown and synthesis, lipolysis, and neurotransmission. Moreover, both reduced the levels of succinate, which couples the tricyclic acid (TCA) cycle to electron transport chain (ETC), signifying an effect on ATP production through oxidative phosphorylation (Smith and Robinson, 2011). Both treatments also induced accumulation of glycolytic intermediates; however, the effects were statistically significant only with isoflurane.

**Figure 4.**
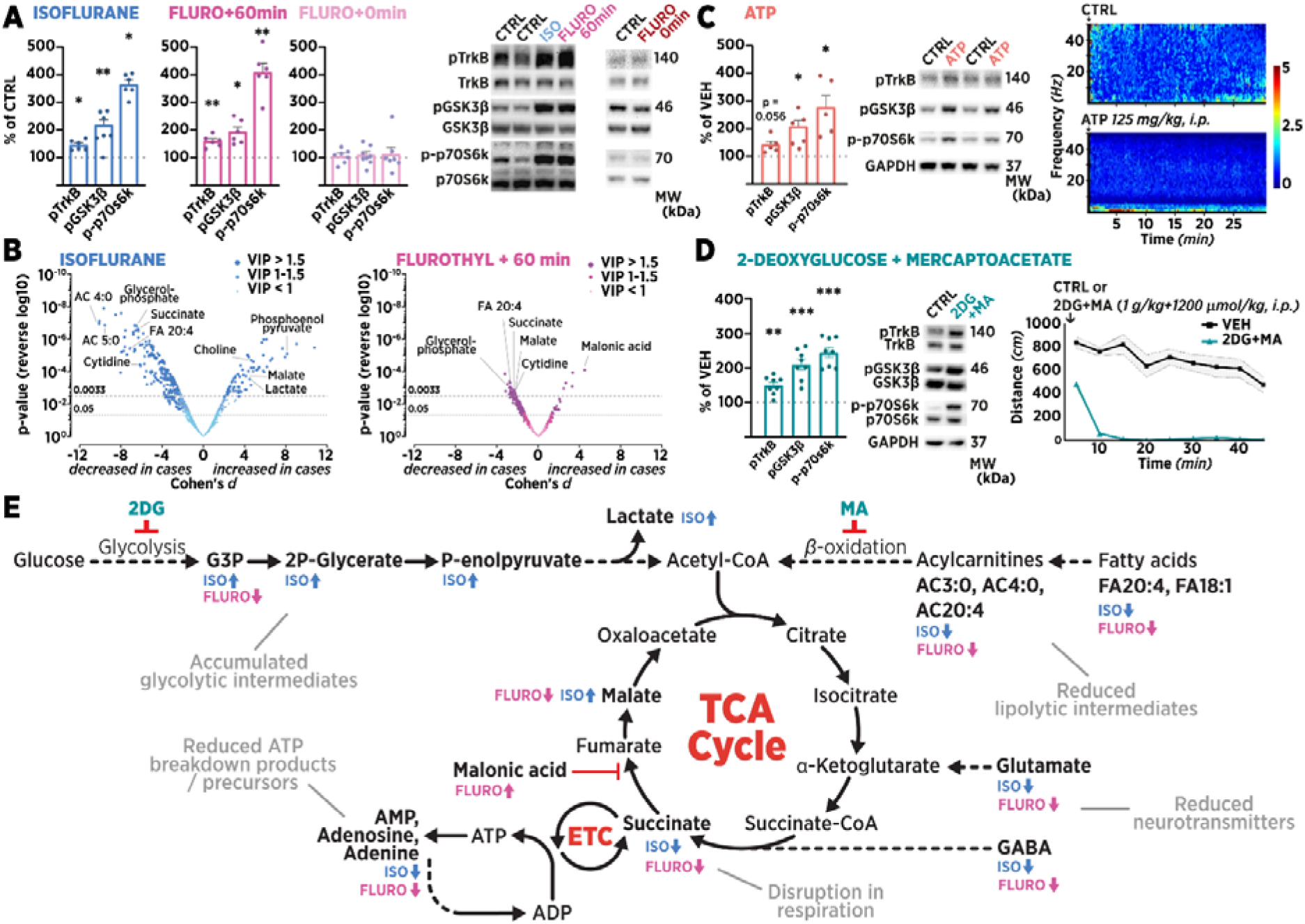
Metabolism and TrkB signaling. **A**. TrkB^Y816^, GSK3β^S9^, and p70SK6^T421/S424^ phosphorylation levels remained regulated during 1 h isoflurane anesthesia and 1 h after a flurothyl-induced seizure. **B**. Continuous isoflurane anesthesia for 1 h and 1 h recovery from brief exposure to flurothyl induce numerous changes in the metabolite profile of mouse prefrontal cortex. Volcano plot demonstrates that the effects of isoflurane are more pronounced than those of flurothyl. For detailed changes, see **Table S2. C**. Adenosine triphosphate (ATP; 125 mg/kg, intraperitoneal [i.p.]) produces a torpor-like state accompanied by electroencephalographic slowing and concomitant TrkB signaling. **D**. Another model of torpor, inhibition of glycolysis and lipid beta-oxidation by injection of 2-deoxy-D-glucose and mercaptoacetate (2DG and MA; 1 g/kg and 1200 μmol/kg, respectively, i.p.), induces sedation and TrkB signaling. **E**. Overview of the bioenergetic changes found in non-targeted metabolomics. Isoflurane anesthesia and flurothyl seizure both reduced succinate levels, which couples the tricarboxylic acid (TCA) cycle to the electron transport chain (ECT) to produce ATP. Both treatments also significantly decreased purinergic breakdown products of ATP, fatty acids (FA), and acylcarnitines (AC). Glycolytic intermediates were accumulated. Inhibition of primary metabolic pathways *in vivo* using 2DG and MA induces the same sedative phenotype and neurotrophic signaling as the sedative and antidepressant drugs used in the study. Phosphoproteins were normalized against the corresponding total protein signal and compared to the control group set to 100%. Data are presented as mean ± standard error of mean. *≤0.05, **≤0.01, ***≤0.005, ****≤0.001 (for statistical analyses and *n* numbers see **Table S1**). Abbreviations: ADP, Adenosine diphosphate; AMP, Adenosine monophosphate; CoA: Coenzyme A; G3P: Glyceraldehyde-3-phosphate; GSK3β, glycogen synthase kinase 3β; 2P-Glycerate: 2-Phosphoglyceric acid; P-enolpyruvate: Phosphoenolpyruvate; p70S6k, ribosomal protein S6 kinase.

Next, we sought to test the intriguing possibility that pharmacological induction of comparable metabolic state using direct inhibitors of energy metabolism regulates TrkB signaling. Two treatments common in the research of mammalian hibernation and torpor were used: ATP (Swoap et al., 2007) and a combination of metabolic inhibitors 2-deoxy-D-glucose and mercaptoacetate (2DG+MA) (Westman and Geiser, 2004). Systemic delivery of ATP slows down the TCA cycle through allosteric inhibition of pyruvate hydrogenase (Martínez-Reyes and Chandel, 2020), producing a state characterized by hypothermia, immobility, and increased slow EEG activity (~1–4 Hz; **Figure 4C**). mPFC samples obtained 30 min after injection showed phosphorylation of TrkB^Y816^, GSK3β^S9^, and p70SK6^T421/S424^ (**Figure 4C**). Administration of 2DG and MA, which inhibit the first steps of glycolysis (hexokinase) and lipid beta-oxidation (fatty-acid CoA dehydrogenase), respectively, also induced profound sedation and TrkB signaling (**Figure 4D**). An overview of the metabolomic changes produced by isoflurane and flurothyl as well as the upstream pathways affected by 2DG+MA are presented in **Figure 4E**. Together, these findings suggest that antidepressant-associated TrkB signaling is activated by two opposing treatments that share the ability to disrupt the bioenergetic balance or by specific inhibition of metabolic pathways upstream of the TCA cycle.

### Modulation of TrkB transactivation and antidepressant effects by thermal control

Next, we investigated whether the sedation-activated TrkB signaling could be prevented by reversing the key changes associated with deep sleep and torpor—hypothermia and EEG slowing —by administering amphetamine, which is known to increase EEG activity (Young, 1988) and thermogenesis (Askew, 1962). Pre-treating the animals with amphetamine blocked medetomidine-induced increase in SWA but only partially prevented the hypothermic effects (**Figure S4A**). Indeed, medetomidine-induced TrkB signaling was only marginally affected by amphetamine in the mPFC and cerebellum (**Figure S4B**). The effects of amphetamine on medetomidine-induced changes in SWA and TrkB signaling may, at least in part, be also related to opposing effects on noradrenergic neurotransmission, which is why we next tested whether the signaling can be regulated directly through temperature.

Homeothermic animals maintain a relatively constant body temperature in a wide range of ambient temperatures by altering the metabolic rate. Consequently, cerebral metabolic rate, for example, changes linearly with the surrounding temperature (Gillooly, 2001; Rosomoff and Holaday, 1954). We set to study whether sedative-induced TrkB signaling can be manipulated using exogenously applied heat during the acute effects of drugs. To this end, a cohort of mice was injected with medetomidine and housed either in a warm incubator maintained at 37 ± 1°C or in room temperature (22 ± 1°C). Core body temperature was measured at 10 min intervals, and the mice were terminated at 30 min. Again, medetomidine induced a progressive reduction in body temperature of the animals at room temperature, which was abolished when the animals were housed in a warm incubator **(Figure S4C)**. Most importantly, preventing the decrease in body temperature decreased the phosphorylation of TrkB^Y816^, GSK3β^S9^, and p70SK6^T421/S424^ by medetomidine (**Figure S4D**).

Next, we set to investigate the role of temperature and metabolic rate in antidepressant-induced TrkB signaling. We chose to use the tricyclic antidepressant amitriptyline as it has been claimed to be—among multiple other drugs—a direct agonist of TrkB receptor (Jang et al., 2009). We studied the acute effects of amitriptyline using a dose (20 mg/kg, i.p.) commonly used in antidepressant research. The treatment produced marked sedation and hypothermia in mice at room temperature **(Figure 5A)**, accompanied by phosphorylation of TrkB^Y816^, GSK3β^S9^, and p70SK6^T421/S424^ in the mPFC samples. Remarkably, in mice housed in a warm incubator, the effect of amitriptyline on both body temperature and neurotrophic signaling was abolished **(Figure 5A)**, whereas only a partial reversal was seen in the hyperglycemic response **(Figure S2B)**. This observation challenges the notion that the effects of the drug are mediated by direct binding to TrkB receptor, but instead suggests that—analogous to the effects of medetomidine, flurothyl, N_2_O, and SD—the mechanism is connected to the ability of the drug to reduce the energy expenditure, and consequently, body temperature. Notably, we repeated the experiment with isoflurane anesthesia with similar results (**Figure S4E**). Thus, we screened the effects of selected TrkB-activating drugs and treatments on body temperature of mice, and found that they all significantly reduced body temperature (**Figure S5**).

**Figure 5.**
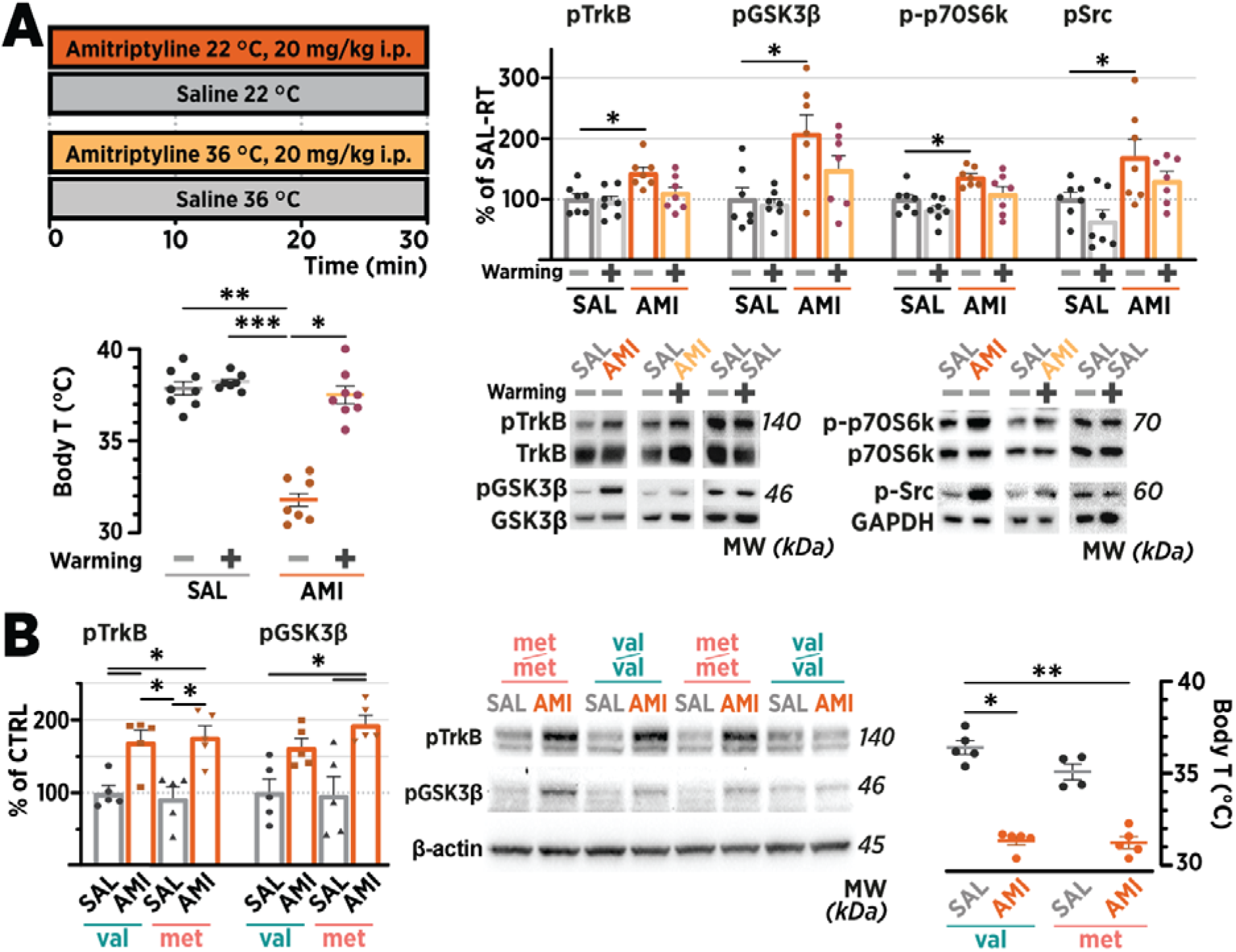
Antidepressants transactivate TrkB through a BDNF-independent, temperature-dependent mechanism. **A**. By 30 min after injection, amitriptyline (AMI; 20 mg/kg, intraperitoneal) caused a significant decrease in the body temperature (Body T) of animals housed at room temperature (22 ± 1°C). Housing the animals in a warm incubator (36 ± 1°C), which had negligible effects on the Body T of saline-treated animals (SAL), blunts the hypothermic effect of AMI. Housing the animals in an incubator (36 ± 1°C) after administration inhibited the AMI-induced phosphorylation of TrkB^Y816^, GSK3β^S9^, p70SK6^T421/S424^, and Src^Y416^ in the medial prefrontal cortex samples collected at 30 min after injection. **B**. AMI readily activates TrkB^Y816^ and GSK3β^S9^ phosphorylation in the medial prefrontal cortex, along with hypothermia, in both Bdnf^val66val^ and Bdnf^met66met^ mice. Phosphoproteins were normalized against corresponding total protein and compared to the control group set to 100%. Data are presented as mean ± standard error of mean. *≤0.05, **≤0.01, ***≤0.005, ****≤0.001 (for statistical analyses and *n* numbers see **Table S1**). Abbreviations: BDNF, brain-derived neurotrophic factor; GSK3β, glycogen synthase kinase 3β.

We have previously shown that both imipramine and isoflurane activate TrkB signaling in the brains of BDNF-deficient mice (Antila et al., 2017; Rantamäki et al., 2011), implying a transactivation mechanism not requiring BDNF. To test this further, we subjected adult mice carrying the met66met mutation in the *Bdnf* gene, which impairs activity-dependent BDNF release (Chen et al., 2004), to acute treatment with amitriptyline (**Figure 5C**). The magnitude of the TrkB signaling response induced by amitriptyline was indistinguishable between the *Bdnf^met66met^* mice and the controls (*Bdnf^val66val^* mice). Considering the established role of Src kinase in regulating neurotrophin-independent Trk receptor transactivation (Lee and Chao, 2001), we assessed Src phosphorylation on the active site in brain homogenates that showed increased TrkB phosphorylation. Indeed, Src^Y416^ phosphorylation was rapidly induced by isoflurane and amitriptyline, and after the acute effects of SD, N_2_O, and flurothyl dissipated (**Figure 5B, Figure S6**). Furthermore, maintaining the mice in a warm incubator fully blocked this phosphorylation change induced by amitriptyline (**Figure 5B**). A non-canonical drug-induced activation mechanism is also supported by the differential phosphorylation of TrkB and extracellular-signal regulated kinase (ERK1/2; **Figure S7**), a well-established downstream target of BDNF-induced TrkB activation (Huang and Reichardt, 2001).

To investigate the functional significance of one of the main findings of this study—TrkB signaling triggered by reduced energy expenditure and hypothermia—we tested whether antidepressant-like behavioral effects can be manipulated by altering the metabolic-thermal responses evoked by antidepressants. N_2_O was used because its pharmacological mechanism of action resembles that of ketamine and its fast kinetics enable more precise differentiation of the acute pharmacological effects and the consecutive evoked physiological responses. We used 60 min exposure of 50% N_2_O (in O_2_), as its rapid and sustained antidepressant effects have been recently revealed (Nagele et al., 2021), but the underlying mechanism remains obscure. We have previously shown that this dosing regimen of N_2_O evokes a similar transient increase in SWA and TrkB signaling (Kohtala et al., 2019a). The behavioral effects of N_2_O were assessed in the chronic corticosterone (CORT) model of depression (**Figure 6A**), previously used to investigate the antidepressant effects of ketamine (Moda-Sava et al., 2019). Animals exposed to CORT showed reduced preference to saccharine, considered to be a sign of anhedonia, which was significantly rescued by N_2_O when tested 32 h after the gas treatment (**Figure 6B**). This behavioral effect was essentially abolished in the animals placed in a warm incubator (36 ± 1°C) for 1 h after the gas exposure. Remarkably, housing the animals in a warm incubator for an hour without prior N_2_O exposure also rescued the CORT-induced decrease in saccharine preference. Warming by itself, however, did not produce as consistent antidepressant-like effect as that with N_2_O. When the saccharine preference was re-tested 1 week later, no treatment effects were noted.

**Figure 6.**
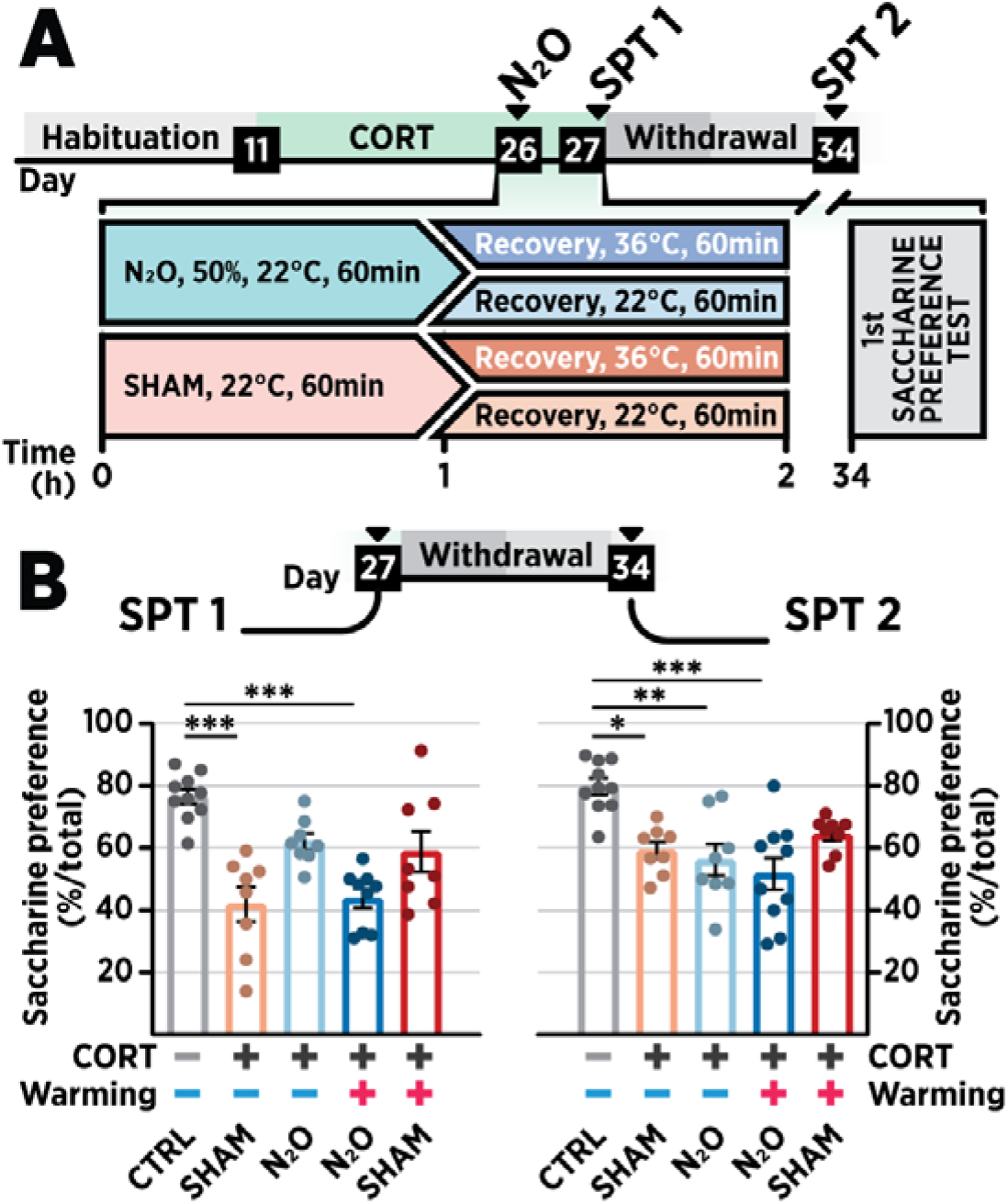
Modulation of TrkB signaling and antidepressant effects through manipulation of ambient temperature. **A**) Schematic overview of the experiment **B)**. A single exposure to nitrous oxide (N_2_O; 50%, 60 min) increased saccharine preference in chronic corticosterone (CORT)-based depression model mice 32 h after treatment administration (SPT1). The effect was fully abolished if the animals were placed in warm ambient temperature (36 ± 1°C) for 60 min during recovery from N_2_O. The antidepressant effects of N_2_O had subsided by the second test 7 days later (SPT2). Data are presented as mean ± standard error of mean. *≤0.05, **≤0.01, ***≤0.005, ****≤0.001 (for statistical analyses and *n* numbers, see **Table S1**).

## Discussion

The effects of antidepressant drugs are commonly explained using classic principles of receptor pharmacology, ignoring the complex, underlying physiological processes. The findings presented in this report suggest that a signaling mechanism considered to be the foundation of antidepressant responses— activation of TrkB signaling—is triggered by an evoked physiological response instead of straightforward pharmacodynamic action. In the present study, the intriguing association between TrkB signaling and bioenergetics was tested by preventing the antidepressant-induced activation of TrkB signaling through manipulation of ambient temperature, and by extension, the metabolic rate. Most importantly, we demonstrate that the antidepressant-induced TrkB activation as well as the behavioral effects are blunted by preventing the drug-induced hypothermic response. To our knowledge, this is the first study causally linking bioenergetics and thermoregulation to the molecular and behavioral effects of antidepressants.

While the ability of antidepressants to induce TrkB signaling has been established since the seminal observations by Saarelainen et al. (2003), the underlying mechanism remains enigmatic. During the past decades, several antidepressants have been proposed to act either by working as direct agonists of TrkB, or more recently, by direct binding to a novel site of TrkB, which facilitates its responsiveness to BDNF (Casarotto et al., 2021; Jang et al., 2009). Ketamine is suggested to have a high propensity to bind to TrkB, whereas conventional antidepressants require chronic administration and accumulation in brain for efficient modulation of the signaling (Casarotto et al., 2021). The authors also suggest that binding to TrkB is specific for antidepressants, whereas other drugs, including antipsychotic chlorpromazine, do not bind and activate the receptor in a similar manner. However, in the present study, despite the lack of homogeneity in structural and pharmacodynamic properties of the compounds tested, all treatments with notable sedative effects— including chlorpromazine and several other non-antidepressants—were found to induce TrkB phosphorylation and signaling *in vivo*. The effects of the treatments on TrkB signaling become evident rapidly, essentially depending on when the sedation and state of reduced energy metabolism emerge. Furthermore, the direct binding hypothesis would not sufficiently explain the identical signaling pattern produced by non-pharmacological interventions that produce similar perturbations in physiology. These results contradict the common notion that emphasizes that increased TrkB signaling is by itself sufficient for antidepressant responses. We argue instead that induced TrkB signaling is only a part of more complicated molecular and functional alterations prerequisite for antidepressant effects.

Previous studies have shown that cerebral energy metabolism is altered in both rodent and primate models of depression and in humans with medication-resistant depression (Detka et al., 2014; Głombik et al., 2020; The Mood Disorders Precision Medicine Consortium et al., 2021). Antidepressants have been shown to acutely impair mitochondrial respiration (Hroudová and Fišar, 2012), whereas studies performed after the acute effects or during chronic administration tend to show facilitation of energy metabolism (Filipović et al., 2017; Perez-Caballero et al., 2021; Scaini et al., 2011). A similar adaptive response has been shown to underlie the antidepressant effects of calorie restriction (Wang et al., 2021). It is conceivable that the observed TrkB signaling is an integral part of hormesis, an adaptive change produced by brief bioenergetic stress (Marosi and Mattson, 2014). Indeed, the metabolomic screening suggested temporal association of TrkB signaling with reduced levels of metabolites associated with ATP production and breakdown, which may represent decreased ATP production capacity (anesthesia) or excessive demand (seizure). This interpretation would be in line with the proposed unified anesthetic mechanism of action, demonstrating a similar sedative–anesthetic state produced *in vivo* in distinct manners that push the bioenergetic capacity to its limit (Voss and Sleigh, 2020). This notion is supported by our findings that the administration of energy metabolism-inhibiting treatments produces significant sedation and TrkB signaling. As mentioned before, further support for the hypothesized physiological origin of the observed phenomenon is provided by the upregulation of neurotrophic signaling during recovery sleep following prolonged waking. Analogous to the transient increase in energy expenditure during exposure to N_2_O or flurothyl, SD is associated with a marked escalation of metabolic demand, followed by homeostatic reduction in energy expenditure and anaplerotic processes during recovery sleep (Jung et al., 2011). Intriguingly, some studies have associated anesthesia and deep sleep with replenishment of ATP reserves in several brain regions, including the mPFC (Dworak et al., 2010, 2011). Notably, a single sedative dose of ATP has been shown to elicit antidepressant-like effects in rodents (Cao et al., 2013), which, as our results suggest, may be associated with TrkB signaling.

Most putative rapid-acting antidepressants share the ability to robustly increase, not decrease, brain energy demand (Kohtala et al., 2020). Regardless of the treatment, the energetic challenge must be compensated, and energy stores replenished after the acute physiological perturbation subsides. Notably, postictal (i.e., after a seizure) EEG slowing has been suggested to predict the antidepressant efficacy and onset-of-action of ECT (Sackeim et al., 1996). Treatments such as N_2_O, subanesthetic-dose ketamine, and ECT-mimicking flurothyl evoke emergence of SWA after their acute pharmacological effects dissipate, although the magnitude and time-scale of the effects differ (Kohtala et al., 2019a). Furthermore, SD gradually builds up excitability in the cerebral cortex (Huber et al., 2013), associated with antidepressant actions in depressed patients, eventually evoking homeostatic emergence of SWA during sleep. Intriguingly, both animal and clinical studies indicate that subanesthetic ketamine leads to very similar augmentation of slow wave sleep (Duncan et al., 2013; Feinberg and Campbell, 1993). We have recently hypothesized that the acute synaptic potentiation induced by subanesthetic ketamine and related treatments renormalize the functional connectivity in depressed patients, leading to rapid amelioration of depressive symptoms (Rantamäki and Kohtala, 2020). We further propose that this acute effect is unstable and requires—according to the principles of the synaptic homeostasis hypothesis of sleep (Tononi and Cirelli, 2014)—consolidation that occurs during subsequent sleep. While this hypothesis needs further testing in patient trials, this report insinuates a pivotal role of TrkB signaling in the plastic processes affected by antidepressant treatments as well as physiological sleep.

The observations presented in this study also provide further insights into the neurobiological mechanisms of therapeutic hypothermia. Hypothermia, among the oldest known neuroprotective treatments (Karnatovskaia et al., 2014), has been shown to rescue the brain from various insults and is physiologically associated with significant structural plasticity (Horowitz and Horwitz, 2019; Popov et al., 2007). The therapeutic effects of hypothermia are widely considered to be mediated through attenuated energy expenditure, oxidative stress, and inflammation; recent studies also highlight the involvement of TrkB signaling. In particular, Peretti and colleagues have shown that RNA-binding motif 3 (RBM3)—a major inducer of cold-induced structural plasticity and neuroprotection (Peretti et al., 2015)—controls the phospholipase-Cγ signaling downstream of TrkB (Peretti et al., 2021). The authors used both pharmacological (5′-AMP) and environmental means (low ambient temperature) to induce and maintain hypothermia, respectively, which was shown to induce BDNF release and TrkB signaling. The employed time frames, however, are considerably longer than those used in the present study and our previous research, showing that distinct hypothermia-inducing treatments upregulate TrkB signaling in just minutes. Considering the temporal dissociation in the observed profile of TrkB phosphorylation and the pharmacodynamic or physiological action of the interventions, as well as the identical response seen in animals with deficient BDNF release, it is difficult to argue that the observed signaling is solely mediated by BDNF release and binding to its target receptor. Particularly, by using treatments such as N_2_O that allow precise temporal control of the pharmacodynamic effects, we have only been able to see TrkB phosphorylation after the supposed activity-dependent release of BDNF ceases. Indeed, this report extends our previous observations suggesting a BDNF-independent mechanism underlying antidepressant-induced TrkB activation (Antila et al., 2017; Rantamäki et al., 2011). While the precise mechanism remain to be fully elucidated, our molecular data support the role for Src family kinase-mediated transactivation (Lee and Chao, 2001). Indeed, Src family kinases, and several molecules connected with TrkB signaling (most notably GSK3β), are regulated by metabolic perturbations through oxidative stress and alterations in neuronal lipid environment (Papadopoli et al., 2021; Park et al., 2016).

### Conclusions

We reveal a strong and unexpected association between antidepressant effects and brain energy metabolism. Various different types of antidepressants, including SD, N_2_O, and conventional antidepressants, trigger a brain state characterized by reduced brain energy expenditure, hypothermia, and TrkB signaling. Most remarkably, maintenance of normothermic body temperature blocks the antidepressant-induced TrkB signaling and behavioral responses. This report challenges pharmacology-centric hypotheses explaining antidepressant-induced TrkB signaling by highlighting the role of bioenergetics and thermoregulation in antidepressant responses and opens critical questions about the mechanistic aspects of TrkB signaling. In particular, our results provide novel insights into how sleep and torpor as well as antidepressant treatments mediate their dramatic effects on neuronal plasticity. Better understanding of these complex physiological processes may enable their application in a more efficient manner in the treatment of neuropsychiatric conditions.

## Supporting information

Table S1. Statistics

## Author contributions

O.A., M.R., S.K. and T.R. planned the experiments; O.A., S.K., M.R., N.M., W.T., R.S., H.M., G.GH, P.K, O.K., A.K., M.S., S.R. and T.R. carried out research; O.A. prepared the figures; O.A., S.K., M.R. O.K. and S.R. run statistical tests, O.A., S.K. and T.R. wrote the paper; O.A., M.R., S.K. and T.R. provided funding; all authors commented the manuscript and accepted final submitted version.

## Materials and Methods

### Animals

Unless otherwise stated, 10–16-week-old C57BL/6JRccHsd mice (Envigo, Venray, Netherlands) were used. Animals were maintained under standard laboratory conditions (22 ± 1 °C, 12 h light−dark cycle, lights on at 6 A.M. or 9 A.M.) with access to food and water available *ad libitum*. BDNF Val66Met mice were generated as previously described (Chen et al., 2006). Briefly, BDNF Val/Met mice were interbred and offspring were genotyped by polymerase chain reaction (PCR) analysis of tail tip-derived genomic DNA. The experiment involving BDNF Val66Met mice was conducted in female BDNF^Val/Val^ and BDNF^Met/Met^ mice. The animal experiments were carried out in compliance with the European Communities Council Directive of 22 September 2010 (Directive 2010/63/EU), according to the guidelines of the Society for Neuroscience, and approved by the County Administrative Board of Southern Finland (Licenses ESAVI/9793/04.10.07/2016 and ESAVI/5844/2019).

### Drug administrations

The following drugs were diluted in isotonic saline and administered intraperitoneally (i.p.) in a volume of 10 ml/kg: adenosine triphosphate (ATP; 125 mg/kg, i.p., Sigma-Aldrich), amitriptyline-HCl (20 mg/kg; s.c.; Tocris), (S)-amphetamine-HCl (10 mg/kg; i.p.; Toronto Research Chemicals; kindly provided by Dr. Esa Korpi, University of Helsinki), atipamezole-HCl (1 mg/kg; s.c.; Antisedan, Orion Pharma), atropine sulfate (100 mg/kg, i.p., Sigma-Aldrich) chlorpromazine-HCl (10 mg/kg; Orion Pharma), (S)-duloxetine-HCl (10 mg/kg; s.c.; Tocris), gamma-hydroxybutyrate sodium (GHB; 275 mg/kg; i.p.; Xyrem; UCB Pharma Ltd.), hydroxyzine-HCl (50 mg/kg, s.c., Sigma-Aldrich), medetomidine-HCl (0.3 mg/kg; i.p.; Domitor; Orion Pharma), mianserine-HCl (20 mg/kg; s.c.; Tocris), mirtazapine-HCl (20 mg/kg; s.c.; Tocris), paroxetine maleate (10 mg/kg; s.c.; Tocris), and urethane (2 g/kg; kindly provided by Dr. Kai Kaila, University of Helsinki). Clozapine (6 mg/kg, 10 ml/kg, i.p.; Sigma-Aldrich) was dissolved in 0.1 M hydrochloric acid, adjusted to pH ~7 with 0.1 M sodium hydroxide, and then diluted to a concentration of 1.2 mg/ml with sterile saline. Equivalent vehicle solution was prepared without the active drug for clozapine control animals. Drug treatments were administered 3–7 hours after the light onset (6 A.M.).

Nitrous oxide (N_2_O), isoflurane, sevoflurane, and flurothyl (Bis(2,2,2-trifluoroethyl) ether; Sigma-Aldrich) were administered as described (Antila et al., 2017; Kohtala et al., 2019a). Briefly, medical grade N_2_O (Linde Healthcare or Woikoski) was mixed with oxygen to achieve a mixture of 50–65% N_2_O. N_2_O was administered into airtight acrylic chambers (for biochemical analyses (width × length × height): 14 × 25 × 9 cm; for biochemical and EEG analyses: 11.5 × 11.5 × 6.5 cm) with a flow rate of 4–8 l/min. Isoflurane anesthesia was induced using 2% isoflurane (Vetflurane, Virbac) and maintained under constant flow of 1.5% isoflurane (Antila et al., 2017). Sevoflurane anesthesia was induced using 6% sevoflurane (Sevoflurane Baxter 100%, Baxter Healthcare) and maintained under 4.5% (Kohtala et al., 2016). Pressurized room air was used as a sham treatment for control animals of inhaled treatments. Anesthetized animals were kept on a prewarmed hot water blanket as a standard procedure to prevent marked hypothermia. Seizure was induced by injecting 10% flurothyl (in 90% ethanol) into a cotton pad placed inside the lid of an airtight Plexiglass chamber (13 × 13 × 13 cm) until the animals exhibited myoclonic seizure. The lid was removed to terminate the seizure, after which the animals were moved to recover in their home cages. For control animals, 90% ethanol without flurothyl was injected to the pad as a sham treatment.

### EEG-recordings

EEG-recordings were performed as described (Kohtala et al., 2019a). Briefly, electrodes were implanted under isoflurane anesthesia. Lidocaine (10 mg/ml) was used as local anesthetic and buprenorphine (0.1 mg/kg, s.c.) and carprofen (5 mg/kg, s.c.) used for postoperative care. Two epidural screw EEG electrodes were placed above the fronto-parietal cortex. A further screw served as mounting support. Two silver wire electrodes were implanted in the nuchal muscles to monitor the EMG (electromyogram). After a recovery period (5-7 days), animals were connected to flexible counterbalanced cables for EEG/EMG recording and habituated to recording cables for three days. The EEG and EMG signals were amplified (gain 5/10 K) and filtered (high pass: 0.3 Hz; low pass 100 Hz; notch filter) with a 16-channel AC amplifier (A-M System, model 3500), sampled at 254 Hz or 70 Hz with 1401 unit (CED), and recorded and processed using Spike2 (version 8.07, Cambridge Electronic Devices). EEG power spectra were calculated within the 1-50 Hz frequency range by fast Fourier transform (FFT = 256, Hanning window, 1.0 Hz resolution). Oscillation power in each bandwidth (delta=1–4 Hz; theta=4–7 Hz; alpha=7–12 Hz; beta=12–25 Hz; gamma low=25–40 Hz; gamma high=60–100 Hz) was computed in 30–300-sec epochs from spectrograms (FFT size: 1024 points) for each animal. Representative sonograms were computed using a Hanning window with a block size of 512.

### Sleep deprivation

Animals were kept awake in their home cages for 6 hours starting from lights on (9 A.M.) by introducing novel objects, or gentle handling whenever the animals appeared sleepy. Animals for biochemical analyses were euthanized using rapid cervical dislocation either immediately after sleep deprivation or 15 minutes after entering sleep, which was determined for each mouse individually as continuous immobility and crouched, sleep-indicating posture.

### Corticosterone model of depression

Depressive phenotype was modeled in animals using chronic corticosterone administration (Moda-Sava et al., 2019). Corticosterone (#46148, Sigma-Aldrich) was dissolved in 99.5% ethanol and diluted with tap water to a final concentration of 100 μg/ml of corticosterone and 1% ethanol (CORT). Mice were exposed to CORT in place of drinking water for 16 days, followed by a weaning phase during which CORT mice received a solution at 50% and then 25% of the full dose for 3 days, respectively. Drinking solutions were replaced every three days. After the weaning period, all mice drank control solution for a total of 3 days until euthanasia.

### Saccharin preference test

Saccharin preference test (SPT) examines the animal’s capacity to experience hedonic pleasure evoked by sweet solutions. Saccharin was chosen instead of sucrose due to saccharin having no caloric value, hence controlling for energy intake. Mice had been habituated to the presence of two drinking bottles. On the test day, drinking was restricted for 8 hours (ZT 5:30–12:30) prior to testing. Saccharin solution was prepared immediately before testing by dissolving saccharin sodium salt hydrate (S1002-500G, Sigma, USA) in tap water to make 0.1% (w/v) solution. The mice were presented saccharin solution in one bottle and water in the other bottle for 4 hours (from ZT 12:30–16:30). Next, the bottles were taken away and experimental solutions were returned to mice. Water and saccharin solution consumption was calculated by weighing the bottles before and after the 4-hour testing period. Saccharin preference was determined by calculating the fraction (%) of saccharin solution consumption divided by total solution consumption.

### High-resolution autoradiographic analysis of functional brain activity

Autoradiographic procedure for the analysis of ^3^H-2-deoxyglucose (2-DG; 1.5μCi/g body weight, #NET549A001MC, PerkinElmer) localization was adapted from the method by Sokoloff et al. (1977) as previously described (Alitalo et al., 2020). Distribution and accumulation of tritiated glucose analog was detected from cryosectioned 20 μm coronal brain sections using a direct particle counting system (Le Beaver; Atlantic Instruments for Research, Nantes, France) with 21-hour acquisition time. Regions of interest (ROI) were chosen for qualitative and quantitative analysis based on our previous experiments and the literature on the effects of NMDAR antagonists in rodents (Miyamoto et al., 2000). ROI were outlined in a proprietary software, Beamage (Atlantic Instruments for Research, Nantes, France), using Paxinos’ and Allen’s mouse brain atlas as a reference (Lein et al., 2007). For semi-quantitative analysis of regional glucose utilization, the activity of a given ROI (cpm/min/mm^2^) was averaged from multiple sections from each animal and normalized to the specific activity of corpus callosum of the same subject (Duncan and Stumpf, 1991). As the regional glucose utilization in white matter is relatively unaffected by fluctuations in metabolic demand, variation resulting from subjects’ glucose homeostasis and amount of injected radiolabel is taken into account (Duncan and Stumpf, 1991). Terminal blood was collected for quantifying circulating glucose and remaining radiolabel concentration using a commercial glucose monitor (FreeStyle Freedom Lite; Abbott, Lake Bluff, IL, USA) and a liquid scintillator (Tri-Carb 2910 TR; PerkinElmer, Waltham, MA, USA), respectively.

### Temperature measurements and thermoregulation

Core body temperature was measured using small-animal rectal probe thermometers (FHC Frederick Haer & Co, Bowdoin, ME, USA; 7001H). The measurements were performed as quickly as possible in order to minimize the increase in body temperature resulting from handling stress. For longitudinal analysis of body temperature, whole-body thermal imaging was used. Thermal images were acquired using a forward-looking infrared (FLIR) P640 thermal camera (FLIR Systems, Inc., Wilsonville, OR, United States; rented from Infradex, Vantaa, Finland), which has a reported temperature range from −40 to +500°C, and a reading accuracy of ±2°C or ±2%. The camera was equipped with a 19-mm focal length lens with an optical angle of 45° and was positioned perpendicular to the imaged plane above the recording chambers. Thermal data was acquired at a rate of one image every 30 seconds, before (baseline; 3 minutes) and after the gas administration (recovery; 14 minutes). Images were analyzed using either FLIR Tools software (version 6.4.18039.1003), where highest radiometric pixel temperature was manually acquired for each rodent cage/image using the rectangle tool, or ThermaCAM Researcher Pro (version 2.10), where the rectangle tool was used to automatically acquire the highest pixel temperatures for each rodent cage/image.

In the experiments involving controlled ambient temperature, animals were kept in vented incubator (Vet-Tech Solutions Ltd, Congleton, United Kingdom) set to maintain the temperature of the chamber at 36±1°C. Incubator temperature was monitored with a separate thermometer. Core temperature measurements were acquired using small-animal portable rectal thermometers (Microtherma 2, Agntho’s AB, Lidingö, Sweden or DC Temperature Controller, FHC, Bowdoin, ME, USA).

### Dissection and processing of the brain samples

Animals were euthanized at indicated times after the treatments by rapid cervical dislocation followed by decapitation. Bilateral mPFC (including prelimbic and infralimbic cortices) was rapidly dissected on a cooled dish and stored at −80 °C until further processing. For the analysis of crude brain homogenates, the samples were homogenized in lysis buffer (137 mM NaCl, 20 mM Tris, 1% NP-40, 10% glycerol, 48 mM NaF, H_2_O, Pierce Protease Inhibitor Mini Tablet (Thermo Scientific; Waltham, MA, USA), Pierce Phosphatase Inhibitor Mini Tablet (Thermo Scientific; Waltham, MA, USA)). After ~15-minute incubation on ice, samples were centrifuged (16,000 × *g*, 15 min, 4°C) and the resulting supernatant was collected for western blot analysis.

Crude synaptosomes were prepared as previously described (Kohtala et al., 2019b). Briefly, brain samples were homogenized in 10% (w/v) ice-cold buffer containing 0.32 M sucrose, 20 mM HEPES pH 7.4, 1 mM EDTA, 1x protease inhibitor cocktail (Roche, Mannheim, Germany), 5 mM NaF, 1 mM Na_3_VO_4_, and 5 mM Na_2_HPO_4_. After centrifugation (800 × *g*, 10 min, 4°C), the supernatant was further centrifuged at 15,300 × *g* for 10 min. The supernatant (cytosolic fraction) was removed and the resultant pellet, designated as crude synaptosomal fraction, was resuspended in a lysis buffer (150 mM NaCl, 50 mM Tris-HCl, pH 7.4, 1% Triton X-100, 0.1% SDS, 1X protease inhibitor cocktail, 2 mM EDTA, 5 mM NaF, 1 mM Na_3_VO_4_, and 5 mM Na_2_HPO_4_).

### Western blot

Sample protein concentrations were measured using Bio-Rad DC protein assay (Bio-Rad Laboratories, Hercules, CA). Proteins (40 μg) were separated with SDS-PAGE under reducing and denaturing conditions and blotted to a PVDF membrane as previously described (Antila et al., 2017). Membranes were incubated overnight in +4°C with following primary phosphoprotein antibodies: anti-p-TrkA^Y785^/p-TrkB^Y816^ (#4168; RRID:AB_10620952; 1:000; Cell Signal Technology, Leiden, Netherlands (CST)), anti-p-GSK3β^S9^ (#9336, RRID: AB_331405, 1:1000, CST), anti-p-p70S6k^T412/S424^ (#9234, RRID:AB_2269803, 1:000, CST), anti-p-ERK1/2^T202/Y204^ (p44/42 MAPK; #9106, RRID: AB_331768, 1:1000, CST), and anti-p-Src^Y416^ (#6943, RRID: AB_10013641, 1:1000, CST). For normalization, the following total protein antibodies were used: anti-TrkB (#4603, RRID:AB_2155125, 1:1000, CST), anti-mTrkB (#AF1494, RRID:AB_2155264, 1:1000, R&D Systems), anti-GSK3β (#9315, RRID:AB_490890, CST), anti-p70s6k (#2708, RRID:AB_390722, CST), anti-ERK1/2 (#9102, RRID:AB_330744, 1:1000, CST), and anti-GAPDH (#2118, RRID: AB_561053, 1:1000, CST). The membranes were washed with TBS/0.1 % Tween (TBST) and incubated with horseradish peroxidase conjugated secondary antibodies (1:5,000 or 1:10,000 in non-fat dry milk, 1 h, room temperature; Bio-Rad Laboratories). Secondary antibodies were visualized using enhanced chemiluminescence (ECL Plus, Thermo Scientific, Vantaa, Finland) for detection by Biorad ChemiDoc MP camera (Bio-Rad Laboratories, Helsinki, Finland).

### Non-targeted metabolomics

The non-targeted metabolomics method has been described in detail before (Klåvus et al., 2020). Briefly, weighed tissue samples were extracted using 100 μL of 80% methanol (v/v H_2_O, LC-MS Ultra CHROMASOLV®, Fluka) per 10 mg of tissue under sonication. After vortexing and centrifugation (13000 rpm, 5 min, 4 °C), the supernatant was collected and filtered to HPLC bottles through Acrodisc CR 4 mm (0.45 μm) filter. Samples were analyzed with a UHPLC-qTOF-MS system (Agilent Technologies) using two different chromatographic techniques, i.e. reversed phase (RP) and hydrophilic interaction (HILIC) chromatography and acquired data in both positive and negative polarity. Welch’s ANOVA was used for statistical analysis. Principal component analysis (PCA) was used for metabolite profiling data to analyze overall variance between all the samples and partial least sum of squares discriminant analysis (PLS-DA) was used to calculate VIP values, identifying molecular features explaining variance between the study groups. 15 principal components were needed to explain 95% of the variance in the metabolomics data, therefore the α level was adjusted to 0.0033 to account for multiple testing. Cohen’s *d* effects sizes were calculated between the treatment and control groups. MassHunter Acquisition (Agilent Technologies, ver.B.04.00) was used for data acquisition, Profinder (Agilent Technologies, ver.B.08.00) was used for feature extraction and peak alignment, the Mass Profiler Professional (MPP, Agilent Technologies, ver.13) for statistics, SIMCA (Umetrics, ver.14.0.0) to perform multivariate analyses, and MS-DIAL (ver.2.90) for metabolite identification.

### Statistical analyses

Depending on whether data were normally distributed or not, either parametric or nonparametric test was used for statistical evaluation. In case of more than two groups, analysis of variance (ANOVA) with post hoc test was used. All statistical analyses were performed with Graphpad Prism software (Versions 8 and 9; La Jolla, CA, USA). All tests were two-sided and a P ≤ 0.05 was considered statistically significant. Details of statistical tests and n numbers for each experiment are shown in **Table S1**.

## Acknowledgements

The Radiochemistry Services of the Helsinki Animal in Vivo Imaging Platform (HAIP) at the Helsinki Institute of Life Science (HiLIFE) and the Biocenter Finland Real-Time Imaging Platform are acknowledged for their contribution to studies involving radiotracers. We would like to thank Dr. Vootele Võikar for helping to conduct the experiments at the Mouse Behavioural Phenotyping Facility (supported by HiLIFE and Biocenter Finland). Oy Woikoski Ab is acknowledged for providing oxygen and nitrous oxide. We are grateful to Sissi Pastell, Virpi Perko, and Maria Partanen for taking care of the animals. Dr. Francis Lee is thanked for providing the BDNF^Val/Val^ and BDNF^Met/Met^ mice. Additionally, Katarzyna Dudek, Piotr Michalowski, Emma Hipkin, Anna Swider, Erika Niittymäki, Janina Järvinen, Surachet Imlimthan, and Iida Hirsso are thanked for helping conduct the experiments. Dr. Mikko Airavaara, Dr. Aaro Jalkanen, Dr. Francis Lee, Dr. Conor Liston and Dr. Nelson Totah are thanked for giving comments on the manuscript.

