## Supplementary Material for "Physiological basis underlying antidepressant-induced activation of TrkB receptors"

**Contents**

- Figure S1-S7

**Supplementary figures**

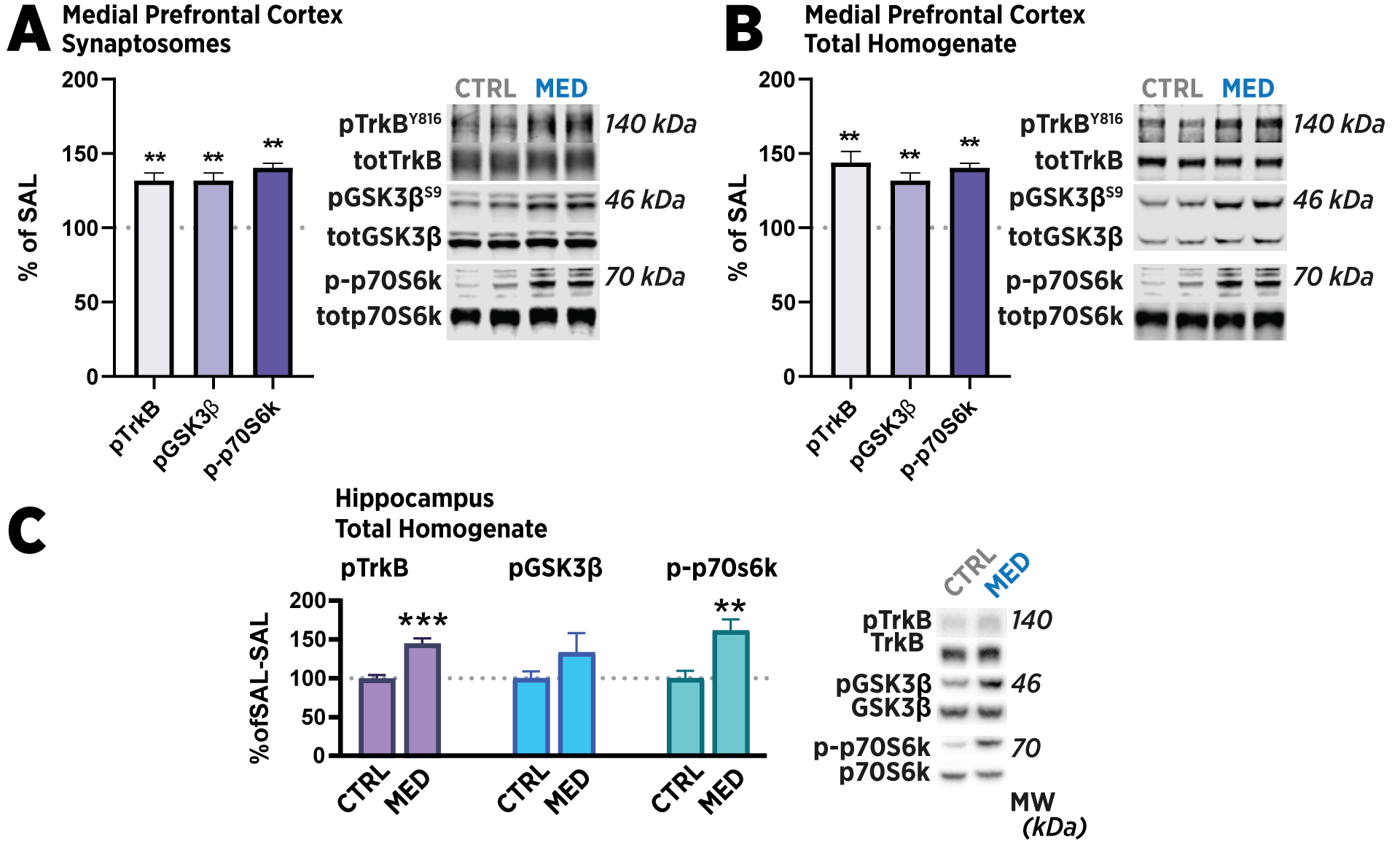

**Figure S1.** **Medetomidine-induced TrkB signaling occurs ubiquitously in different separated fractions and brain regions.** Medial prefrontal cortex samples were collected 30 minutes after medetomidine (MED; 0.3 mg/kg) or saline (CTRL) injection (i.p.). Crude isolated synaptosomal fraction (**A**) and total brain homogenate (**B**) were prepared as from dissected mouse medial prefrontal cortex as described (Kohtala et al., 2019) for the analysis of TrkB^Y816^, and GSK3β^S9^ and p70SK6^T421/S424^ phosphorylation. In a similar manner, total homogenates from mouse hippocampus were prepared and analyzed for TrkB^Y816^, and GSK3β^S9^ and p70SK6^T421/S424^ phosphorylation **(C)**. Phosphoproteins were normalized against corresponding total protein, and the saline group set to 100%. Data are means ± S.E.M. Mann-Whitney. p *≤0.05, **≤0.01, ***≤0.005.

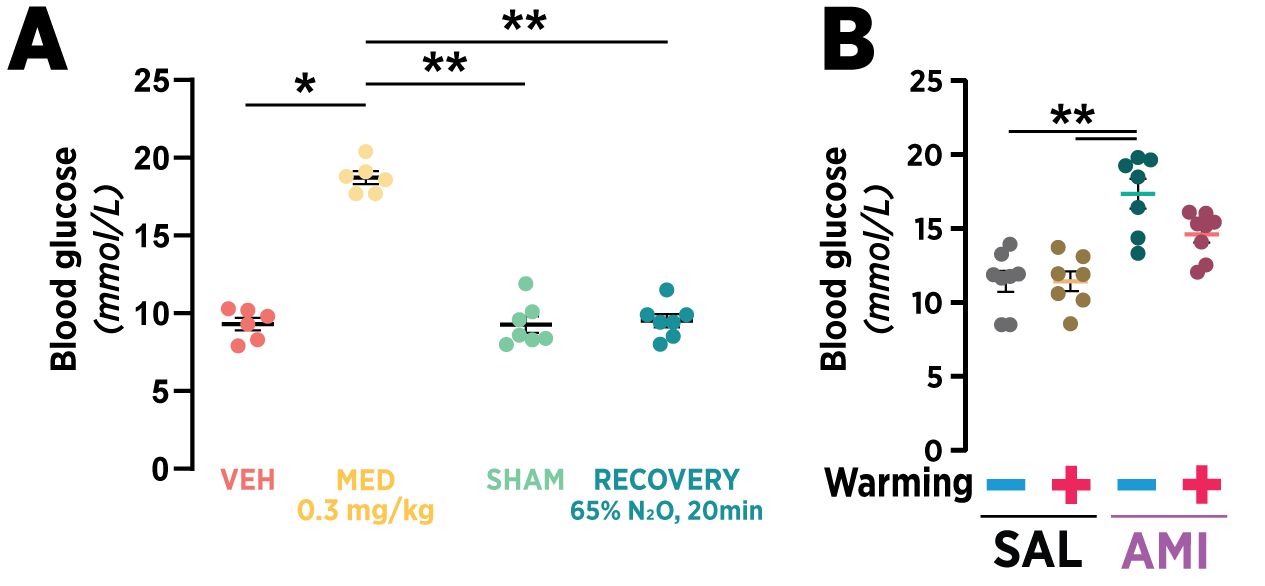

**Figure S2. Changes in blood glucose after administration of treatments inducing TrkB signaling. A.** Medetomidine (MED; 0,3 mg/kg, i.p.) increases terminal blood glucose measured at 30 minutes after injection, while no effect on blood glucose is seen at the end of 20-minute recovery following 20 minutes of nitrous oxide (N2O; 65 % in O_2_). Medetomidine-like effect on blood glucose is seen after amitriptyline (AMI; 20 mg/kg, i.p.) administration at 30 minutes after administration. The effect is attenuated, but not completely abolished, by holding the animals in an incubator after injection until termination. **B.** Even more prominent increase in blood glucose, along with sedated behavior, is seen after chlorpromazine injection (CLP; 10 mg/kg, i.p.). Control animals were treated with equal volume of saline (CTRL) or sham gas administration (SHAM; pressurized room air). Data are means ± S.E.M. Kruskal-Wallis with Dunn’s multiple comparisons test. p *≤0.05, **≤0.01.

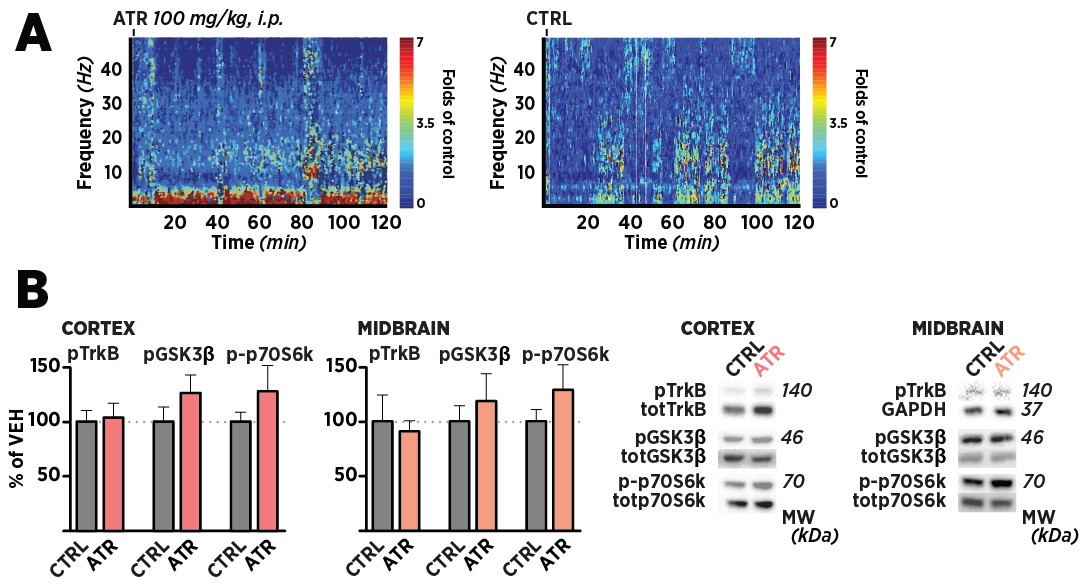

**Figure S3. Despite slowing down cortical activity, atropine does not** **acutely activate TrkB signaling. A.** EEG recorded from a prefrontal cortex-implanted electrode for two hours after atropine (ATR; 100 mg/kg, i.p.) injection shows marked reduction in electrophysiological activity, while the phenotype of the animal was observed to be relatively unaltered. EEG of vehicle-treated animals (CTRL) shown on right. **B.** Medial prefrontal cortex and midbrain samples collected from atropine-treated animals 30 minutes after injection show negligible activation of TrkB^Y816^, and GSK3β^S9^, and p70SK6^T421/S424^. Data are means ± S.E.M. Mann-Whitney test.

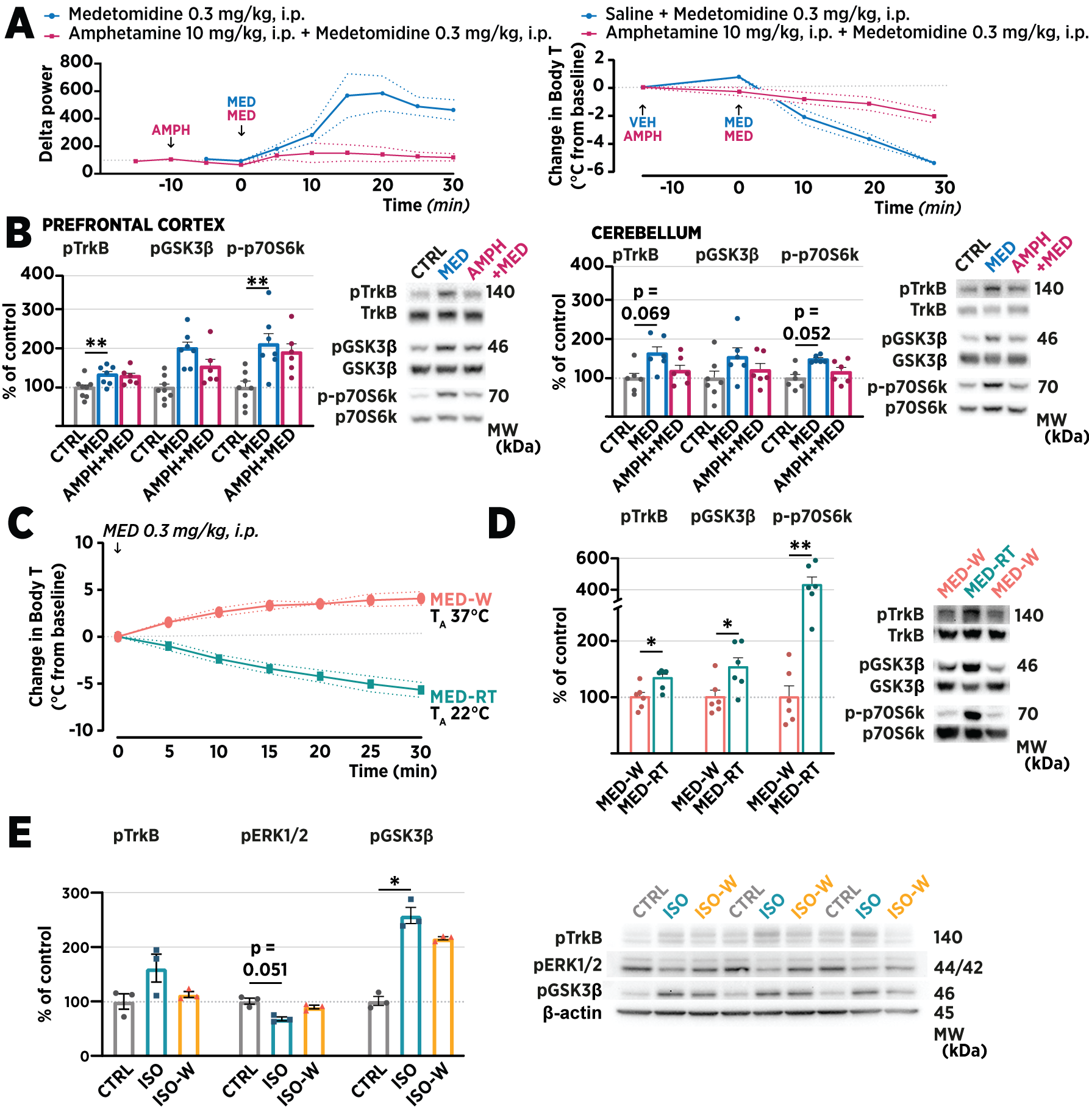

**Figure S4. Disrupting of hypothermia attenuates the effects of medetomidine and isoflurane on TrkB signaling. A**. The impact of thermogenic amphetamine (AMPH; 10 mg/kg, i.p.) pretreatment on medetomidine (MED; 0.3 mg/kg, i.p.)-induced SWA and decrease in core body temperature (Body T). **B**. AMPH fully blocked MED-induced activation of TrkB signaling in the cerebellum, while only partial effect was seen in the medial prefrontal cortex. Treatment groups are compared to a sham-treated group (CTRL). **C**. Keeping MED-treated mice in warm ambient temperature (MED-W; 37 ± 1°C) maintains Body T of the animals slightly above physiological range. **D**. In comparison to room temperature-housed animals (MED-RT), warm ambient temperature attenuates the MED-induced phosphorylation of TrkB^Y816^, GSK3β^S9^, and p70SK6^T421/S424^ in the medial prefrontal cortex samples collected at 30 minutes after injection. **E**. Maintaining the mice in warm ambient temperature attenuates the phosphorylation of TrkB^Y816^ and GSK3β^S9^ induced by isoflurane (ISO) anesthesia. Maintaining the temperature (ISO-W) also prevents the isoflurane-induced decrease in ERK1/2^T202/Y204^ phosphorylation. Phosphoproteins were normalized against corresponding total protein, and the control group set to 100%. Data are means + S.E.M. B,E: Kruskal-Wallis with Dunn’s multiple comparison test; D: Mann-Whitney. Data are means + S.E.M. *≤0.05, **≤0.01, ***≤0.005, ****≤0.001

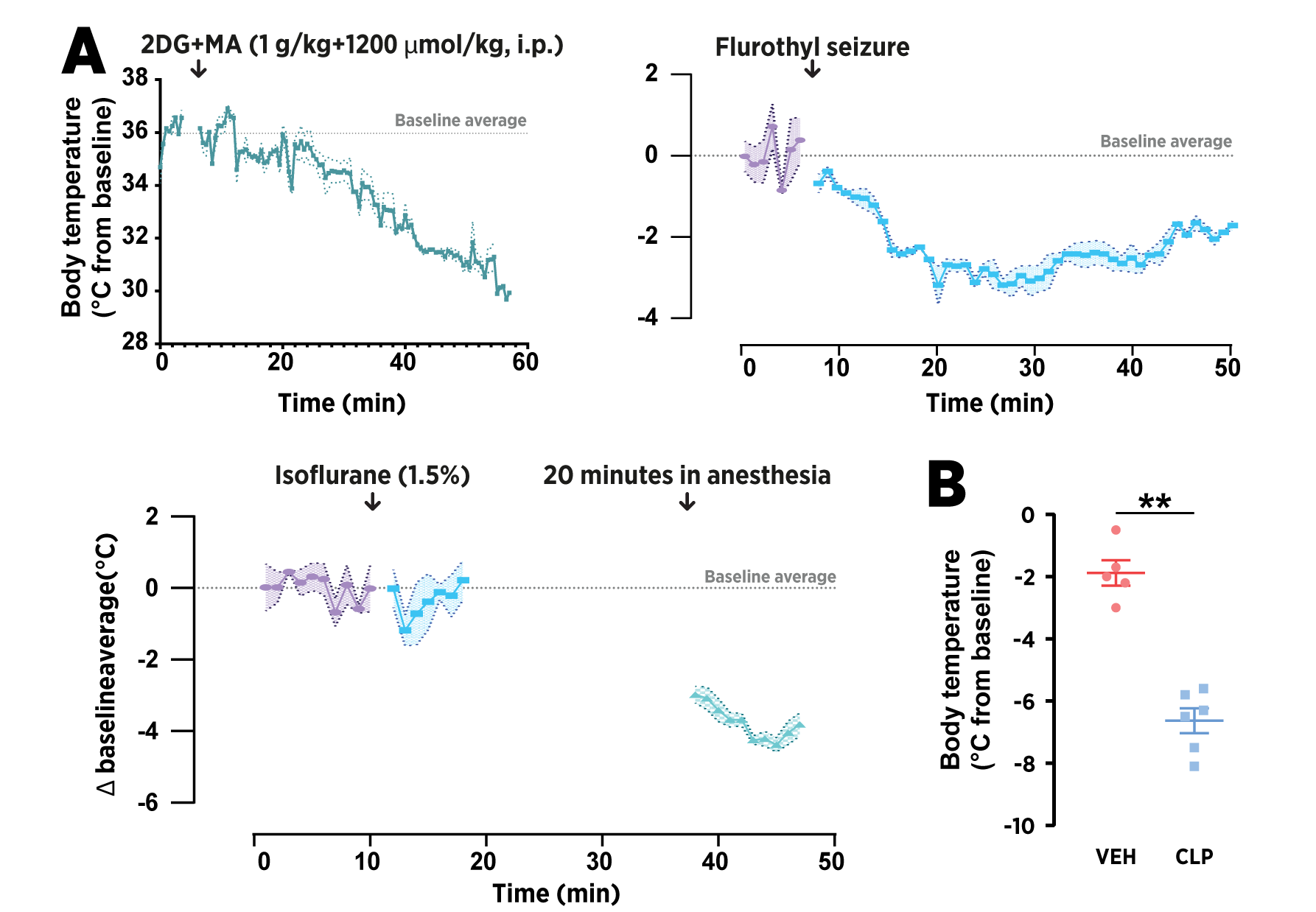

**Figure S5. Prominently TrkB-activating treatments induce hypothermia. A.** FLIR camera screening of effects of different drugs on body temperature in unrestrained animals. Temperature data was recorded after administration of treatments on small cohort of animals (n=2–3). Isoflurane (1.5%) was administered in a separate chamber and the animals were moved to the imaging chamber separately, resulting in the apparent gradual increase of the temperature during the recording periods. **B.** Rectal temperature measurement at 30 minutes after injection demonstrates prominent hypothermic effect of chlorpromazine (CLP; 10 mg/kg, i.p.). Data are means + S.E.M. Mann-Whitney. p **≤0.01

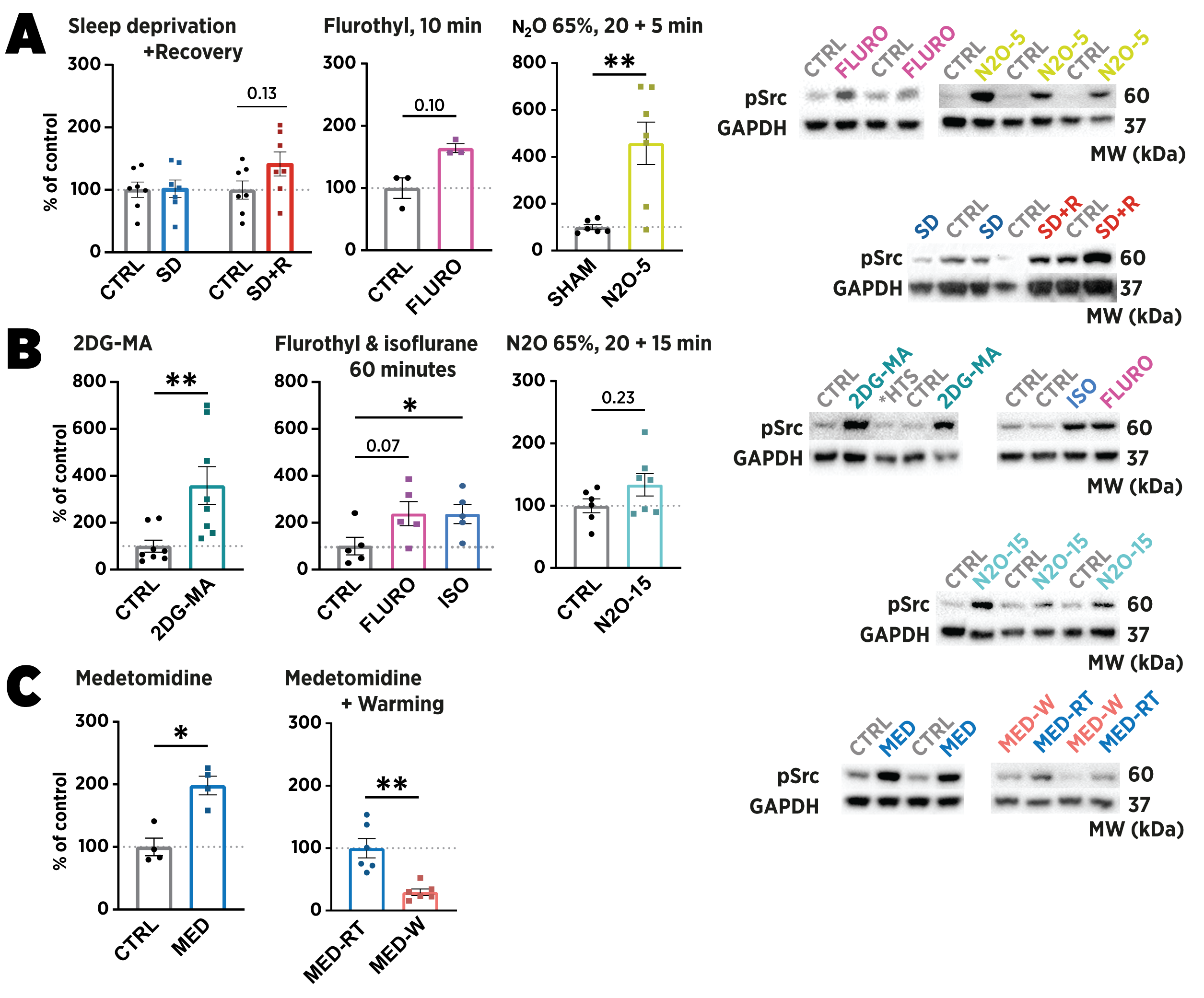

**Figure S6.** **Src family kinase phosphorylation during sedative-state and deep sleep. A.** Src family kinases are phosphorylated at Y416 residue in the medial prefrontal cortex samples collected in the early phase of the homeostatic recovery. Phosphorylation at the end of sleep deprivation (SD) is unaffected, while 15 minutes of recovery sleep (SD+R) shows increase. Similarly, Src is activated 10 and 5 minutes after the cessation of flurothyl (FLURO) seizure and nitrous oxide (N2O; 65 %, 20 minutes) exposure, respectively. **B.** Sustained metabolic inhibition using 2-deoxyglucose and mercaptoacetate (2DG-MA; 1 g/kg and 1200 µmol/kg, respectively, i.p.) shows marked increase in the phosphorylation of Src^Y416^ in samples collected at 45 minutes after injection. 1 hour after FLURO seizure or continuous isoflurane (ISO; 1.5 %) anesthesia, more variability in pSrc ^Y416^ is observed. In full blots incubated with the same pSrc ^Y416^ antibody, clear regulation associated with the treatment is observed in an unknown band of approximately 90 kDa molecular weight. **C.** The activation of Src kinase, along with TrkB signaling, is temperature-dependent. At 30 minutes, medetomidine (MED; 0.3 mg/kg, i.p.) induces prominent phosphorylation of pSrc ^Y416^, which is blunted by keeping the animals in an incubator (37 ± 1°C). The control animals for injected treatments received physiological saline (CTRL), while inhaled treatments were controlled using pressurized room air (SHAM). Phosphoproteins were normalized against corresponding total protein, and the control group set to 100%. Mann-Whitney and Kruskal-Wallis followed by Dunn’s multiple comparison test. Data are means + S.E.M. *≤0.05, **≤0.01.

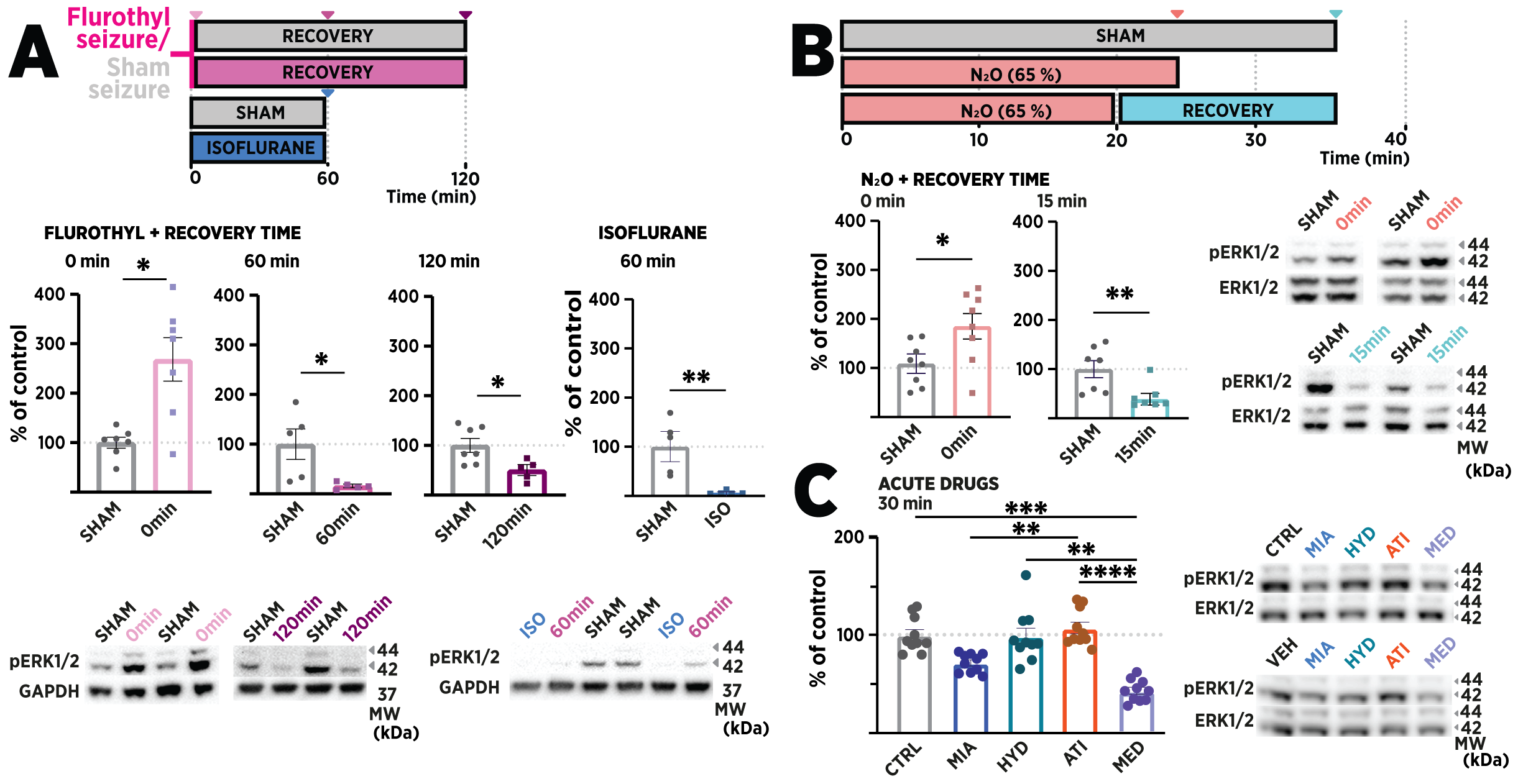

**Figure S7. Non-canonical, sedation-coupled activation of TrkB signaling is associated with reduced ERK1/2 phosphorylation.** Extracellular signal-regulated kinase (ERK1/2; p44/42 MAPK) is phosphorylated in T202/Y204 residues during acute excitatory stimuli such as flurothyl seizure or ongoing 65% nitrous oxide administration (N_2_O; in O_2_) **(A–B)**, during which phosphorylation of TrkB^Y816^, GSK3β^S9^, and p70SK6^T421/S424^ is unaffected. However, as animals become sedated when the excitatory stimulus subsides, the phosphorylation of ERK1/2^T202/Y204^ is rapidly downregulated to a fraction of the baseline level. Analogous to the postictal sedative state, direct sedation with isoflurane (ISO; **A**) or other diverse sedative-anesthetic drugs (**C**) reduces ERK1/2^T202/Y204^ phosphorylation while increasing the phosphorylation of TrkB^Y816^, GSK3β^S9^, and p70SK6^T421/S424^. Drugs, doses, and routes administered in **C** were: mianserin (MIA; 20 mg/kg, s.c.), hydroxyzine (HYD; 50 mg/kg, s.c.), atipamezole (ATI; 1 mg/kg, s.c.), and medetomidine (MED; 0.05 mg/kg, i.p.). Control animals for injected pharmacological agents were administered vehicle (VEH; physiological saline) in equal volume. Control animals for flurothyl and isoflurane were subjected to sham treatments (SHAM) consisting of brief holding in induction chamber and pressurized room air, respectively. Data are means ± S.E.M. A-B Mann-Whitney, C Kruskal-Wallis with Dunn’s multiple comparison test. p * ≤ 0.05, ** ≤ 0.01, *** ≤ 0.005, **** ≤ 0.001.
