## Supplementary File A for "Physiological basis underlying antidepressant-induced activation of TrkB receptors"

**Contents**

- Supplementary File A, Figures 1–3, full autoradiographic acquisition plates
- Supplementary method for scintillator analysis of plasma radioactivity
- Supplementary File A, Table 1 and 2, the results of plasma radioactivity analysis

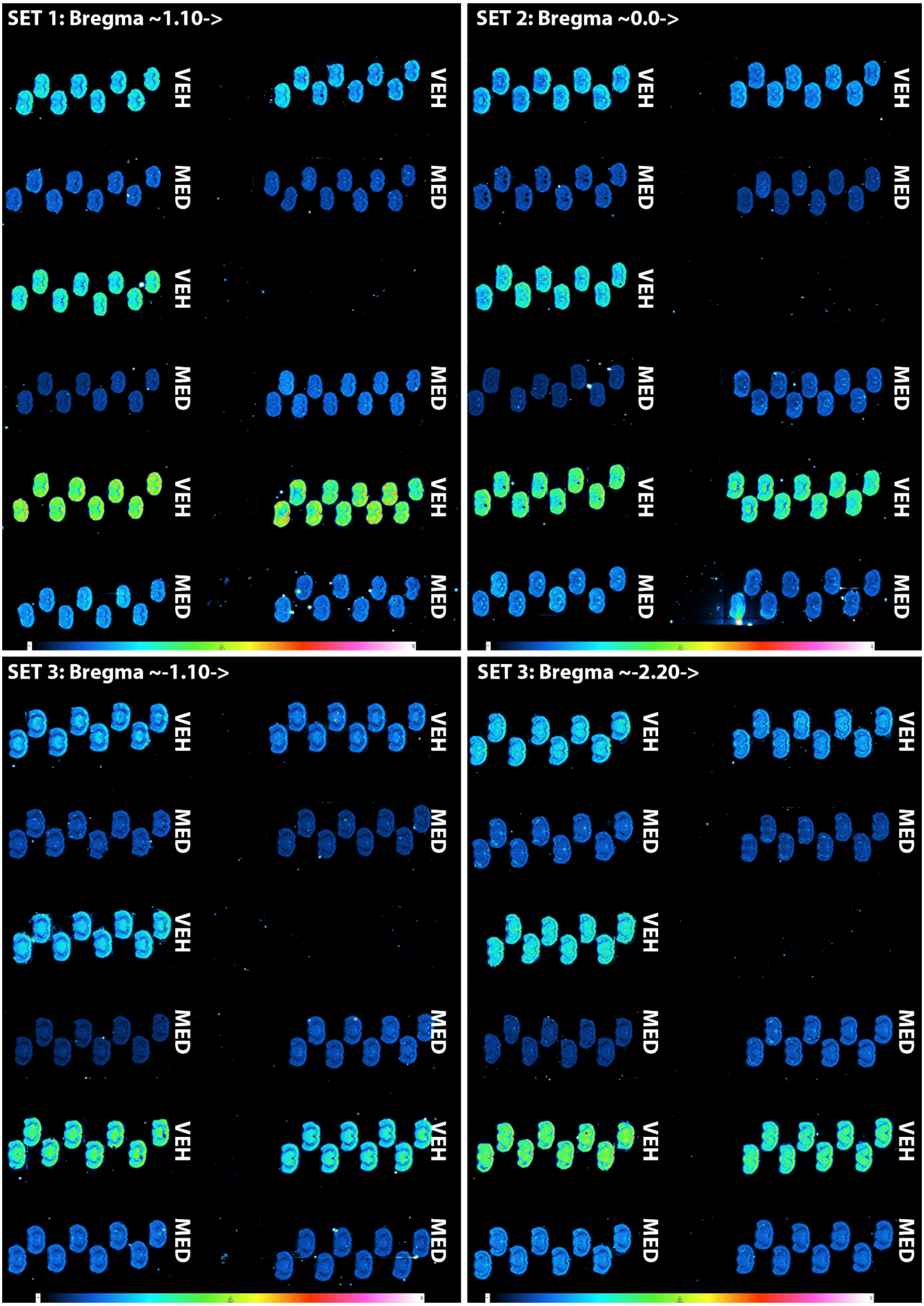


**Supplementary File B, Figure 1.** Full 21-hour acquisition plates from samples collected from animals treated with medetomidine (MED; 0.3mg/kg, i.p.) and saline (VEH). Each animal was injected tritiated 2-deoxy-D-glucose ($^{3}$H-2-DG; ~1.5 µCi/g body weight) 20 minutes before termination, 30 minutes after administration of MED. 4 sets of 20 µm thick coronal brain sections were prepared every 100 µm beginning 1.10 cm from bregma. One animal was discarded from analysis due to missed i.p. injection.

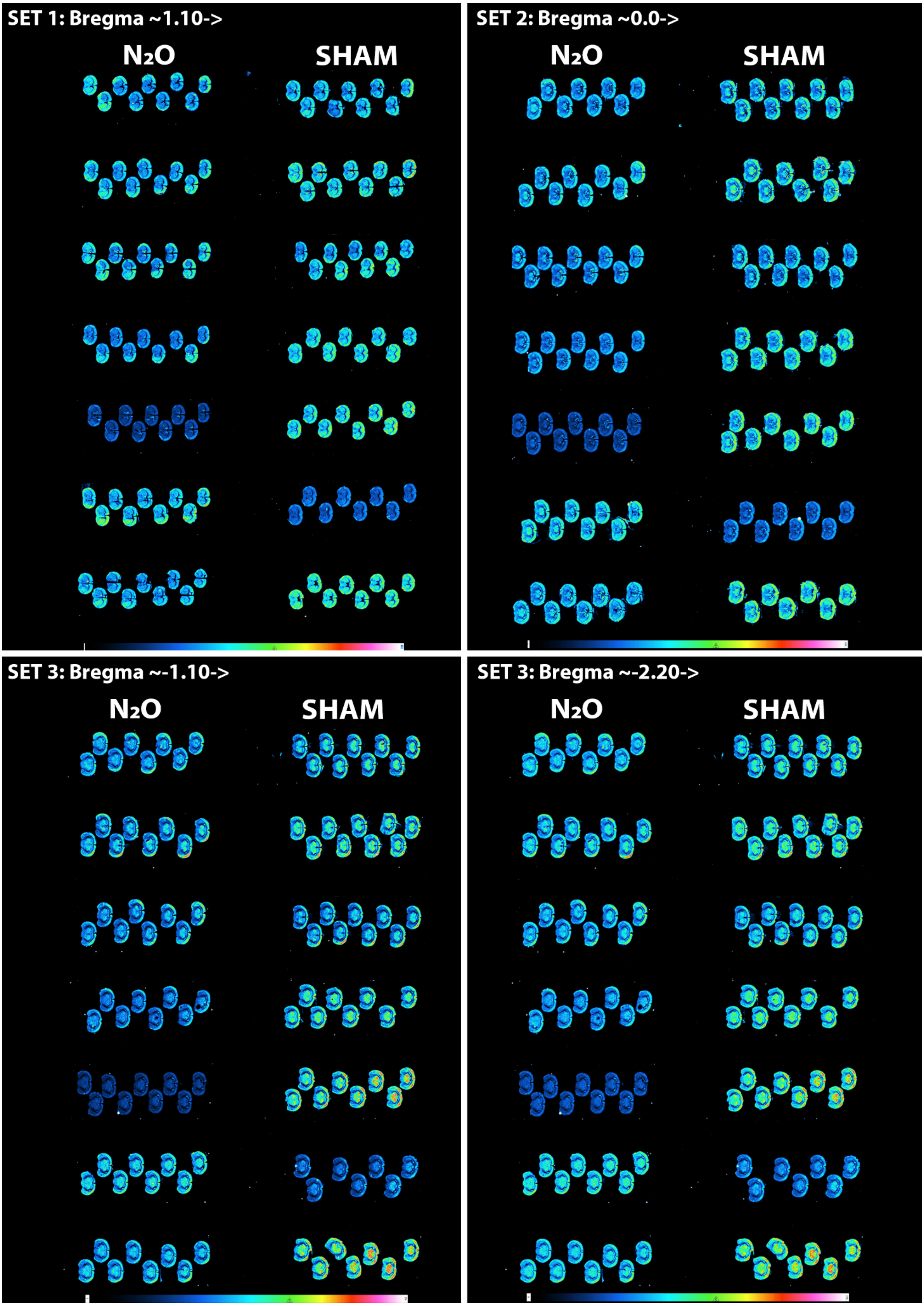


**Supplementary File B, Figure 2.** Full 21-hour acquisition plates from samples collected from animals recovered 20 minutes from either a 20-minute exposure to nitrous oxide ($N_2O$; 65%) or sham treatment (pressurized room air). Each animal was injected tritiated 2-deoxy-D-glucose ($^3$H-2-DG; ~1.5 µCi/g body weight) as they were moved from $N_2O$ chamber to recovery, 20 minutes before termination. 4 sets of 20 µm thick coronal brain sections were prepared every 100 µm beginning 1.10 cm from bregma.

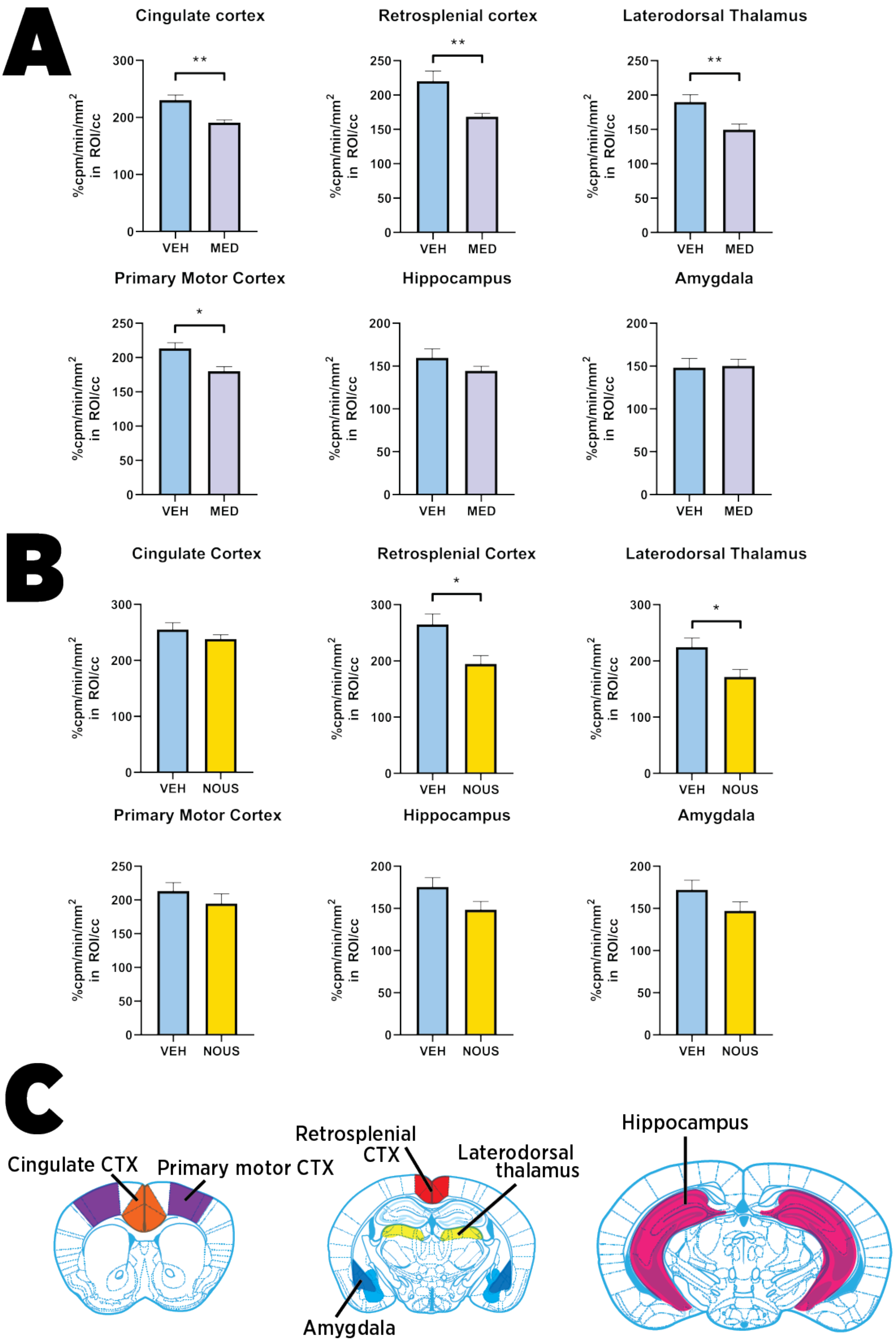


**Supplementary File B, Figure 3. Semiquantitative analysis of the distribution of radiolabeled glucose analog 2-deoxy-D-glucose. A.** Accumulation of 2-DG was significantly reduced in cortical and thalamic structures of medetomidine-treated (MED; 0.3 mg/kg, i.p.) animals in comparison to animals that received vehicle. **B.** During the recovery from 65% $N_2O$ exposure, glucose uptake was slightly reduced in retrosplenial cortex and laterodorsal thalamus, while other regions displayed similar general trend. **C.** Representative brain atlas images from set 1 and 3 and 4 demonstrating the average locations of regions of interest (ROI) used for quantification. ROI were averaged from multiple sections of each animal and normalized to the activity of corpus callosum.

*Supplementary method for liquid scintillator analysis of plasma radioactivity*

Further details of the method are described in the main article materials and methods. Briefly, as the animals were terminated and decapitated for dissection of brain tissue, trunk blood was used for the measurement of blood glucose concentration, after which the all available blood was collected to a K-EDTA tubes (Vacuette, Fisher Scientific) and mixed well to prevent coagulation. Tubes were centrifuged at 2000 × g for 10 minutes, and the supernatant—plasma—was collected to a separate set of tubes and stored in -80°C until further analysis.

For liquid scintillation, 50 µl of plasma from each animal was transferred to a liquid scintillation vial and mixed with 1 ml of MilliQ and 15 ml of scintillator liquid (Optiphase Hisafe 3, PerkinElmer, Boston, MA, USA). Similar procedure was used to prepare a background sample from non-radioactive rodent plasma. Furthermore, two reference samples were prepared using known concentration of tritium from a standard capsule (Internal Standard Kit, 3H-W, Quantulus-Option, 1210-121, PerkinElmer).

The vials were wiped with ethanol and inserted to the liquid scintillator (Tri-Carb 2910 Tr, PerkinElmer). Two 5-minute detections were conducted for each sample. The pipetted volume (5 µl) of plasma was weighed and converted into corresponding volume using the density of plasma (1.025 g/ml). The error in pipetted volume was calculated according to the manufacturer instructions (0.6%; FinnPipette, Sartorius Lab Instruments GmbH, Goettingen, Germany).

Activity of the tritium standard capsules in the sample volume of 16.05 ± 0.0963 ml was corrected from the initial value of 3756.7 ± 30.4 Bq on 2.10.2017 with the following calculation, resulting an activity of 3517.4 ± 28.9 Bq per one standard capsule:

$$A = A_0 * 2^{-\frac{t}{t_{1/2}}}$$

$A$= Activity of the sample on measurement day (Bq)
$A_0$= Activity of the sample in reference day (Bq)
$t$ = time between measurement and reference date (s)
$t_{1/2}$ = half-life of the radionuclide (s)

Error of decay-corrected activity was then calculated using following equation:

$$\Delta A = \sqrt{\left(2^{-\frac{t}{t_{\frac{1}{2}}}} * \Delta A_0\right)^2 + \left(A_0 * ln2 * \frac{t}{t_{\frac{1}{2}}^2} * 2^{-\frac{t}{t_{\frac{1}{2}}}} * \Delta t_{\frac{1}{2}}\right)^2}$$

Next, the sample activities were corrected for the background activity using the equation:

$$cpm = cpm_a - cpm_{background}$$

Error for cpm (count rate) was calculated from the error (2 $\sigma$) given by the measurement instrument (example calculation for standard sample 1 in the first measurement):

$$\Delta cpm = \left(\frac{\frac{0{,}33}{100}}{2}\right) * 72084 = 118.9386 \sim \mathbf{119\ cpm}$$

Activity for plasma samples could be calculated with the known activity and the mean of counts per minute of the standard samples (example calculation done for mouse 1 31.10.2018) using equation:

$$A_{plasma\ sample} = \frac{A_{tritium\ standard\ capsule}}{cpm_{mean\ of\ cpm\ of\ standard\ samples}} * cpm_{plasma\ sample}$$

$$A_{31.10.2018, mouse\ 1} = \frac{3517.400218\ Bq}{752576\ cpm} * 71904\ cpm$$

Error for activity of plasma sample was calculated with error propagation law:

$$\Delta A_{plasma\ sample}$$

$$= \sqrt{\left(\frac{A_{standard}}{cpm_{std}} * \Delta cpm_{plasma}\right)^2 + \left(\frac{1}{cpm_{std}} * cpm_{plasma} * \Delta A_{std}\right)^2 + \left(\frac{1}{cpm_{std}^2} * \Delta cpm_{plasma} * cpm_{plasma} * A_{std}\right)^2}$$

$$\Delta A_{31.10.2018, mouse\ 1} = \sqrt{\left(\frac{3517.4\ Bq}{72576\ cpm} * 118.64\ cpm\right)^2 + \left(\frac{1}{72576\ cpm} * 71904\ cpm * 28.8\ Bq\right)^2 + \left(\frac{1}{(72576\ cpm)^2} * 118.64\ cpm * 71904\ cpm * 3517.4\ Bq\right)^2}$$

$$= 31.91802533\ Bq \sim \mathbf{31.9\ Bq}$$

$$\rightarrow \boldsymbol{A_{plasma\ sample} = 3485 \pm 32\ Bq}$$

Then the activity concentration was calculated by dividing the activity of plasma sample with the total volume of plasma sample (around 50 μl) using the following equation. The error of activity concentration was calculated again using previously described error propagation law. Example calculations done for the same subject below.

$$c_A = \frac{A_{plasma}}{V_{plasma}}$$

$$c_{A,31.10.2018\,mouse\,1} = \frac{3484.8317\,Bq}{0.048175\,ml} = 72336.93196\frac{Bq}{ml} \sim \mathbf{72337\frac{Bq}{ml}}$$

$$c_A = \sqrt{\left(\left(\frac{1}{V_{plasma}} * \Delta A_{plasma}\right)^2 + \left(\frac{1}{V_{plasma}^2} * A_{plasma} * \Delta V_{plasma}\right)^2\right)}$$

$$c_{A,31.10.2018\,mouse\,1}$$

$$= \sqrt{\left(\left(\frac{1}{0.048175\,ml} * 31.9\,Bq\right)^2 + \left(\frac{1}{(0.048175\,ml)^2} * 3484.8317\,Bq * 0.00028905\,ml\right)^2\right)}$$

$$= 792.047\frac{Bq}{ml} = \mathbf{792\frac{Bq}{ml}}$$

$$\rightarrow \mathbf{c_{A,31.10.2018\,mouse\,1} = 72337\ \pm 792\frac{Bq}{ml}}$$

The results of the calculations and the resulting actions are summarized in the Supplementary File A, **Table 1** and **Table 2** below.

**Table 1.** Summary of the results of the Medetomidine experiment using $^3$H-2-DG.

| # | $c_A$ (Bq/ml) | |
|---|---|---|
| 1 | 72268 ± 1119 (1.6 %) | |
| 2 | 50980 ± 793 (1.6 %) | |
| 3 | 71894 ± 1114 (1.6 %) | |
| 4 | 698 ± 20 (2.8 %) | Insufficient injection of the label; subject discarded from qualitative and semiquantitative analysis. |
| 5 | 56379 ± 876 (1.6 %) | |
| 6 | 45500 ± 710 (1.6 %) | |
| 7 | 89362 ± 1381 (1.6 %) | |
| 8 | 80272 ± 1242 (1.6 %) | |
| 9 | 53836 ± 799 (1.5 %) | |
| 10 | 67426 ± 1045 (1.6 %) | |
| 11 | 80708 ± 1248 (1.6 %) | |
| 12 | 54029 ± 840 (1.6 %) | |

**Table 2.** Summary of the results of the $N_2O$ experiment using $^3$H-2-DG.

| # | $c_A$ (Bq/ml) | |
|---|---|---|
| 1 | 57332 ± 891 (1.6 %) | |
| 2 | 56768 ± 881 (1.6 %) | |
| 3 | 65370 ± 1013 (1.6 %) | |
| 4 | 61466 ± 954 (1.6 %) | |
| 5 | 53670 ± 834 (1.6 %) | |
| 6 | 70355 ± 1090 (1.6 %) | |
| 7 | 54655 ± 849 (1.6 %) | |
| 8 | 86224 ± 1333 (1.6 %) | |
| 9 | 111557 ± 1720 (1.6 %) | Too high amount of dosed label: animal discarded from qualitative analysis; taken into account in normalization of semiquantitative analysis -> i.e. subject was not found to be an outlier when the specific regional activity was normalized to the activity of corpus callosum |
| 10 | 61670 ± 957 (1.6 %) | |
| 11 | 80374 ± 1243 (1.6 %) | |
| 12 | 64281 ± 997 (1.6 %) | |
| 13 | 67956 ± 1053 (1.6 %) | |
| 14 | 81112 ± 1255 (1.6 %) | |
