## Supplementary File B for "Physiological basis underlying antidepressant-induced activation of TrkB receptors"

Supplementary File B: Identified metabolites with $p < 0.05$ in the ANOVA comparison

| | | | | | | | | Control | | Flurothyl | | Isoflurane | | ANOVA | Flurothyl vs. Control | | | Isoflurane vs. Control | | |
|---|---|---|---|---|---|---|---|---|---|---|---|---|---|---|---|---|---|---|---|---|
| Group | Metabolite | Exact mass | RT | RP/HILIC | POS/NEG | ID | MSMS | Mean | SD | Mean | SD | Mean | SD | p | t-test p | Cohen's d | PLS-DA VIP | t-test p | Cohen's d | PLS-DA VIP |
| TCA cycle | | | | | | | | | | | | | | | | | | | | |
| | **Malate** | 134.0215 | 5.65 | HILIC | NEG | 2 | 10 eV: 115.0036 (100), 132.8681 (75), 71.0123 (45) | 799196 | 20797 | 697928 | 60986 | 974553 | 64168 | 3.32E-05 | 3.75E-03 | -2.48 | 2.04 | 1.90E-04 | 4.13 | 1.73 |
| | **Succinate** | 118.0266 | 1.14 | HILIC | NEG | 1 | 10 eV: 73.0293 (100), 117.0190 (25) | 1922160 | 146618 | 1623283 | 86658 | 978355 | 151663 | 3.50E-06 | 2.45E-03 | -2.56 | 2.10 | 2.35E-07 | -6.33 | 1.87 |
| | **Malonic acid** | 104.0107 | 1.41 | HILIC | NEG | 2 | 10 eV: 59.0127 (100) | 41417 | 12831 | 151002 | 35982 | 30827 | 8437 | 5.29E-07 | 8.26E-05 | 4.49 | 2.38 | 1.21E-01 | -1.00 | 0.91 |
| Glycolysis | | | | | | | | | | | | | | | | | | | | |
| | **Glyceraldehyde-3-phosphate (G3P)** | 169.9980 | 6.39 | HILIC | NEG | 2 | 10 eV: 96.9693 8100), 78.9585 (35) | 190867 | 34987 | 139962 | 21218 | 262930 | 53482 | 2.40E-04 | 1.45E-02 | -1.81 | 1.82 | 1.50E-02 | 1.63 | 1.26 |
| | **2-Phosphoglyceric acid (2PGA)** | 185.9929 | 7.37 | HILIC | NEG | 2 | 10 eV: 78.9589 (100), 184.9854 (45), 96.9676 (40) | 161566 | 10138 | 175134 | 16682 | 331845 | 42262 | 3.97E-07 | 1.02E-01 | 1.01 | 1.18 | 2.13E-05 | 6.50 | 1.84 |
| | **Phosphoenolpyruvate (PEP)** | 167.9823 | 6.88 | HILIC | NEG | 2 | 10 eV: 78.9592 (100) | 692082 | 50465 | 782286 | 113299 | 2073624 | 298612 | 2.15E-08 | 9.19E-02 | 1.10 | 1.21 | 1.24E-05 | 7.92 | 1.87 |
| | **Lactate** | 90.0316 | 1.50 | HILIC | NEG | 2 | 10 eV: 43.0187 (100), 89.0236 (85) | 106622 | 5599 | 105271 | 11429 | 182339 | 28036 | 2.95E-05 | 7.88E-01 | -0.16 | 0.28 | 2.89E-04 | 4.50 | 1.75 |
| Acetylcarnitines | | | | | | | | | | | | | | | | | | | | |
| | Acetylcarnitine | 203.1159 | 2.96 | HILIC | POS | 1 | 10 eV: 85.0282 (100), 204.1237 (90), 60.0808 (30), 145.0491 (15) | 47224971 | 2592011 | 43923153 | 3419182 | 40201839 | 4261751 | 1.27E-02 | 7.40E-02 | -1.10 | 1.28 | 4.47E-03 | -2.05 | 1.40 |
| | **AC 3:0 (Propionylcarnitine)** | 217.1319 | 2.09 | HILIC | POS | 1 | 10 eV: 85.0285 (100), 218.1367 (30), 159.0657 (15) | 1989933 | 314216 | 1443556 | 192609 | 994186 | 195839 | 1.46E-04 | 5.95E-03 | -2.16 | 1.98 | 1.38E-04 | -3.90 | 1.76 |
| | **AC 4:0 (Butyrylcarnitine)** | 231.1473 | 1.99 | RP | POS | 1 | 10 eV: 85.0281 (100), 232.1542 (75), 173.0772 (5) | 262474 | 28235 | 219385 | 41897 | 32097 | 17356 | 3.48E-06 | 5.12E-02 | -1.23 | 1.42 | 1.13E-07 | -10.11 | 1.93 |
| | **AC 5:0 (Isovalerylcarnitine)** | 245.1635 | 2.89 | RP | POS | 1 | 10 eV: 246.1724 (100), 85.0294 (65), 187.0971 (55) | 235946 | 35975 | 237631 | 33923 | 31174 | 15625 | 1.23E-06 | 9.33E-01 | 0.05 | 0.51 | 6.19E-06 | -7.94 | 1.91 |
| | AC 6:0 (Hexanoylcarnitine) | 259.1787 | 1.09 | HILIC | POS | 1 | 10 eV: 85.0272 (100), 201.1119 (25), 260.1875 (25) | 313220 | 84832 | 246581 | 38809 | 142197 | 104333 | 1.47E-02 | 1.21E-01 | -1.08 | 1.26 | 7.63E-03 | -1.81 | 1.36 |
| | **AC 18:0 (Stearoylcarnitine)** | 427.3670 | 9.59 | RP | POS | 2 | 20 eV: 85.0294 (100), 428.3788 (20) | 653744 | 77372 | 641975 | 81110 | 429472 | 86983 | 1.91E-03 | 7.94E-01 | -0.15 | 0.49 | 4.61E-04 | -2.73 | 1.62 |
| | **AC 20:4 (Arachidonoylcarnitine)** | 447.3366 | 9.07 | RP | POS | 2 | 20 eV: 85.0298 (100), 448.3386 (40) | 1663720 | 563199 | 782503 | 250529 | 405219 | 100342 | 2.36E-05 | 1.02E-02 | -2.17 | 1.91 | 2.50E-03 | -3.79 | 1.68 |
| Lipids and phopsholipids | | | | | | | | | | | | | | | | | | | | |
| | **Glycerol 3-phosphate** | 172.0139 | 6.64 | HILIC | POS | 1 | 40 eV: 80.9730 (100), 98.9798 (5), 62.9628 (5) | 121430 | 11911 | 98128 | 7148 | 59449 | 13757 | 1.11E-04 | 3.10E-03 | -2.45 | 2.17 | 2.91E-06 | -4.83 | 1.83 |
| | **Phosphorylcholine** | 183.0661 | 7.03 | HILIC | POS | 1 | 10 eV: 184.0736 (100), 86.0964 (40), 124.9998 (25), 60.0813 (10) | 40713803 | 1910940 | 36552996 | 2865592 | 33565365 | 1405844 | 5.23E-05 | 1.04E-02 | -1.74 | 1.72 | 3.25E-05 | -4.31 | 1.78 |
| | FA 16:1 | 354.1496 | 10.58 | RP | NEG | 1 | 10 eV: 253.2165 (100) | 102796 | 22511 | 98493 | 23064 | 67100 | 15178 | 1.17E-02 | 7.54E-01 | -0.19 | 0.88 | 1.93E-02 | -1.89 | 1.43 |
| | FA 18:0 | 284.2716 | 11.17 | RP | NEG | 1 | 10 eV: 283.2642 (100) | 4690528 | 354565 | 3797288 | 665073 | 3575009 | 564837 | 3.95E-03 | 1.40E-02 | -1.75 | 1.66 | 1.88E-03 | -2.43 | 1.49 |
| | **FA 18:1** | 282.2556 | 10.92 | RP | NEG | 1 | 10 eV: 281.2483 (100) | 2641734 | 180760 | 2059895 | 335030 | 1463586 | 317244 | 8.42E-05 | 3.40E-03 | -2.26 | 1.91 | 1.22E-05 | -4.73 | 1.78 |
| | FA 20:1 | 310.2778 | 11.23 | RP | NEG | 2 | 20 eV: 309.2700 (100) | 91185 | 7068 | 71306 | 9410 | 67379 | 26153 | 4.09E-03 | 1.92E-03 | -2.41 | 2.05 | 5.46E-02 | -1.43 | 1.02 |
| | FA 20:2 | 308.2709 | 11.02 | RP | NEG | 2 | 10 eV: 307.2589 (100) | 58812 | 11404 | 49862 | 10443 | 36022 | 8319 | 7.36E-03 | 2.02E-01 | -0.82 | 1.04 | 6.77E-03 | -2.31 | 1.53 |
| | **FA 20:4** | 304.2402 | 10.68 | RP | NEG | 2 | 10 eV: 303.2318 (100) | 21198170 | 1398947 | 15919156 | 2475434 | 12188371 | 1133212 | 9.37E-07 | 9.36E-04 | -2.73 | 2.05 | 3.82E-06 | -7.12 | 1.88 |
| | **FA 22:4** | 332.2710 | 10.95 | RP | NEG | 2 | 10 eV: 331.2643 (100) | 316843 | 32710 | 340334 | 76478 | 170287 | 49635 | 4.08E-04 | 4.88E-01 | 0.43 | 0.59 | 1.08E-04 | -3.56 | 1.69 |
| | **FA 22:6** | 328.2415 | 10.66 | RP | POS | 2 | 10 eV: 329.2487 (100) | 225370 | 31242 | 235215 | 38106 | 134080 | 45553 | 2.75E-03 | 6.19E-01 | 0.28 | 0.39 | 1.47E-03 | -2.38 | 1.54 |
| | LysoPE 16:0 | 453.2868 | 10.27 | RP | POS | 2 | 10 eV: 282.2885 (100), 454.2870 (40) | 627745 | 33414 | 633122 | 39799 | 687128 | 27439 | 9.64E-03 | 7.96E-01 | 0.15 | 0.61 | 6.31E-03 | 1.95 | 1.41 |
| Amino acids and analogues | | | | | | | | | | | | | | | | | | | | |
| | **2-Aminoisobutyric acid** | 103.0634 | 5.16 | HILIC | POS | 1 | 10 eV: 87.0443 (100), 69.0331 (50), 45.0334 (43), 43.0180 (20), 41.0387 (10) | 14908908 | 612514 | 14008100 | 771080 | 12974238 | 597623 | 5.80E-04 | 3.89E-02 | -1.30 | 1.45 | 1.50E-04 | -3.20 | 1.67 |
| | Asparagine | 132.0535 | 6.37 | HILIC | NEG | 2 | 10 eV: 114.0192 (100), 70.0286 (25) | 449083 | 57701 | 434937 | 69003 | 342130 | 43562 | 1.04E-02 | 6.95E-01 | -0.22 | 0.34 | 4.55E-03 | -2.11 | 1.46 |
| | **Aspartic acid** | 133.0376 | 6.84 | HILIC | POS | 1 | 10 eV: 74.0236 (100), 88.0396 (75), 134.0471 (10) | 3674665 | 68711 | 3447717 | 173736 | 2513005 | 161600 | 3.15E-07 | 1.29E-02 | -1.87 | 1.75 | 7.86E-08 | -10.09 | 1.91 |
| | N-Acetylaspartic acid (NAA) | 175.0480 | 5.55 | HILIC | NEG | 2 | 20 eV: 59.0297 (100), 88.0408 (70) | 8959762 | 271458 | 9100389 | 421086 | 8546754 | 301692 | 2.94E-02 | 4.85E-01 | 0.41 | 0.57 | 2.49E-02 | -1.44 | 1.19 |
| | **gamma-Aminobutyric acid (GABA)** | 103.0634 | 4.67 | HILIC | POS | 1 | 20 eV: 45.0337 (100), 41.0379 (90), 87.0452 (80), 69.0319 (50) | 13118394 | 732444 | 12218484 | 752779 | 11078853 | 572011 | 5.95E-04 | 5.23E-02 | -1.21 | 1.44 | 3.09E-04 | -3.13 | 1.67 |
| | **Glutamate** | 147.0529 | 6.46 | HILIC | POS | 1 | 10 eV: 84.0446 (100), 102.0539 (45), 130.0514 (45), 148.0588 (20), 56.0492 (5) | 37254775 | 974568 | 35949740 | 1015899 | 34711394 | 873836 | 1.89E-03 | 3.81E-02 | -1.31 | 1.49 | 5.68E-04 | -2.75 | 1.61 |
| | Glutathione (reduced) | 307.0840 | 6.79 | HILIC | POS | 1 | 10 eV: 179.0495 (100), 76.0215 (90), 162.0227 (85), 308.0906 (55), 233.0583 (35) | 17357288 | 431087 | 17123876 | 797613 | 16179891 | 655762 | 1.20E-02 | 5.20E-01 | -0.38 | 0.49 | 2.87E-03 | -2.17 | 1.44 |
| | **Isoleucine** | 131.0948 | 4.27 | HILIC | POS | 1 | 10 eV: 86.0954 (100), 132.1020 (2) | 689907 | 69350 | 694166 | 65122 | 1166516 | 176530 | 2.89E-05 | 9.12E-01 | 0.06 | 0.10 | 1.70E-04 | 3.88 | 1.71 |
| | **Leucine** | 131.0950 | 4.06 | HILIC | POS | 1 | 10 eV: 86.0973 (100), 132.1014 (20) | 827952 | 109924 | 821064 | 117895 | 1778821 | 392878 | 1.95E-05 | 9.15E-01 | -0.06 | 0.21 | 4.57E-04 | 3.78 | 1.68 |
| | Methionine | 149.0512 | 4.61 | HILIC | POS | 2 | 10 eV: 133.0322 (100), 104.0509 (20), 56.0495 (10) | 441675 | 77322 | 411742 | 124760 | 528536 | 61954 | 4.40E-02 | 6.09E-01 | -0.30 | 0.40 | 5.27E-02 | 1.25 | 1.09 |
| | Proline | 115.0636 | 4.95 | HILIC | POS | 1 | 20 eV: 70.0650 (100), 116.0690 (45) | 3597493 | 405568 | 3846994 | 601993 | 5096907 | 993954 | 5.93E-03 | 3.95E-01 | 0.50 | 0.69 | 6.22E-03 | 2.14 | 1.40 |
| | **Spermidine** | 145.1580 | 7.85 | HILIC | POS | 1 | 10 eV: 146.1661 (100), 72.0810 (45), 112.1095 (15), 84.0400 (15) | 270293 | 19741 | 273143 | 28408 | 354324 | 7538 | 2.21E-05 | 8.36E-01 | 0.12 | 0.41 | 4.98E-05 | 6.16 | 1.84 |
| | Threonic acid | 136.0373 | 5.72 | HILIC | NEG | 2 | 20 eV: 75.0090 (100), 72.9925 (25), 71.0139 (20), 134.8657 (10) | 242640 | 56823 | 266107 | 70956 | 408324 | 101394 | 9.84E-03 | 5.22E-01 | 0.37 | 0.57 | 4.38E-03 | 2.09 | 1.45 |
| Purines and purine derivatives | | | | | | | | | | | | | | | | | | | | |
| | **Adenine** | 135.0545 | 1.18 | HILIC | POS | 2 | 10 eV: 136.0610 (100), 119.0311 (10) | 801693 | 79446 | 674031 | 55418 | 538413 | 62434 | 2.11E-04 | 9.47E-03 | -1.89 | 1.82 | 8.15E-05 | -3.71 | 1.73 |
| | **Adenosine** | 267.0974 | 1.42 | RP | POS | 1 | 10 eV: 136.0620 (100), 268.1044 (25) | 14516477 | 3061924 | 10575627 | 2799726 | 6461706 | 2534091 | 1.46E-03 | 3.62E-02 | -1.34 | 1.51 | 4.86E-04 | -2.88 | 1.63 |
| | **Adenosine monophosphate (AMP)** | 347.0635 | 6.70 | HILIC | POS | 1 | 20 eV: 136.0608 (100), 97.0252 (40), 119.0357 (10) | 6290335 | 627424 | 5058866 | 786977 | 2659299 | 760756 | 1.85E-05 | 9.49E-03 | -1.74 | 1.75 | 1.33E-06 | -5.23 | 1.84 |
| | **Cytidine** | 243.0860 | 3.56 | HILIC | POS | 1 | 40 eV: 112.0506 (100), 95.0219 (25), 94.0403 (18) | 528912 | 93180 | 318426 | 90807 | 123637 | 38826 | 1.68E-06 | 1.87E-03 | -2.29 | 1.98 | 3.69E-05 | -6.14 | 1.84 |
| | Cytidine monophosphate (CMP) | 323.0524 | 7.97 | HILIC | POS | 2 | 10 eV: 112.0490 (100), 95.0242 (50) | 68739 | 2733 | 63468 | 4695 | 63077 | 4106 | 2.21E-02 | 3.10E-02 | -1.42 | 1.50 | 1.36E-02 | -1.66 | 1.26 |
| | **Hypoxanthine** | 136.0390 | 1.45 | HILIC | POS | 2 | 10 eV: 137.0447 (100), 94.0363 (20) | 167448 | 20512 | 124111 | 13215 | 91730 | 41917 | 9.60E-04 | 1.96E-03 | -2.57 | 2.10 | 2.23E-03 | -2.43 | 1.51 |
| | Inosine | 268.0813 | 2.62 | HILIC | POS | 1 | 10 eV: 137.0443 (100) | 325921 | 85173 | 238149 | 68033 | 189122 | 86419 | 2.77E-02 | 7.11E-02 | -1.15 | 1.35 | 1.56E-02 | -1.59 | 1.26 |
| Other metabolites | | | | | | | | | | | | | | | | | | | | |
| | **Pantothenic acid (B5)** | 219.1115 | 2.37 | RP | POS | 1 | 10 eV: 90.0551 (100), 220.1175 (80), 116.0324 (45), 202.1072 (30), 184.0967 (20) | 1047927 | 163176 | 826079 | 81117 | 1106051 | 88114 | 4.40E-04 | 1.90E-02 | -1.82 | 1.81 | 4.59E-01 | 0.46 | 0.46 |
| | **Acetylcholine** | 145.1105 | 0.83 | HILIC | POS | 1 | 10 eV: 146.1163 (100), 87.0439 (55), 60.0797 (43) | 6975532 | 645781 | 8127035 | 1643368 | 20385752 | 6667646 | 3.07E-04 | 1.26E-01 | 1.01 | 1.17 | 1.72E-03 | 3.67 | 1.63 |
| | **Choline** | 103.0996 | 1.50 | HILIC | POS | 2 | 10 eV: (104.1064 (100), 60.0803 (45), 45.0331 (25) | 41742404 | 2871488 | 44568931 | 5719455 | 61774032 | 5801525 | 1.10E-05 | 2.80E-01 | 0.66 | 1.11 | 2.05E-05 | 4.62 | 1.81 |
| | 1-Methylhistamine | 125.0953 | 4.64 | HILIC | POS | 2 | 10 eV: 126.1012 (100), 109.0744 (70), 68.0485 (40) | 332635 | 50238 | 318335 | 57943 | 212885 | 62077 | 9.97E-03 | 6.43E-01 | -0.26 | 0.38 | 2.74E-03 | -2.13 | 1.47 |
| | 3-Methylhistidine | 169.0852 | 6.96 | HILIC | POS | 2 | 10 eV: 170.0920 (100), 109.0752 (35), 153.0707 (20) | 129626 | 22159 | 164894 | 32783 | 197274 | 32083 | 4.83E-03 | 4.31E-02 | 1.28 | 1.67 | 1.04E-03 | 2.49 | 1.56 |
| | beta-Homoproline | 129.0792 | 1.01 | HILIC | POS | 2 | 10 eV: 70.0641 (100), 130.0852 (30) | 138050 | 9347 | 139484 | 14864 | 157043 | 9314 | 9.05E-03 | 8.37E-01 | 0.12 | 0.25 | 3.95E-03 | 2.04 | 1.51 |
| | N-Palmitoylethanolamide (PEA) | 299.2833 | 10.44 | RP | POS | 2 | 10 eV: 62.0594 (100), 300.2867 (80) | 134290 | 32087 | 453511 | 503331 | 561776 | 765775 | 3.81E-02 | 1.45E-01 | 1.19 | 1.08 | 1.90E-01 | 1.07 | 0.73 |
| | Triethanolamine | 149.1056 | 2.72 | HILIC | POS | 2 | 10 eV: 150.1107 (100), 44.0480 (25), 132.0971 (10), 88.0733 (5) | 176426 | 36848 | 207666 | 36414 | 154142 | 24721 | 3.71E-02 | 1.55E-01 | 0.85 | 1.09 | 2.42E-01 | -0.72 | 0.71 |

Legend: AC, acetylcarnitine; Bold, p-value below multiple comparison corrected α (0.0033); d, Cohen's d effect size; FA, fatty acid; ID, level of identification (1 = in-house standard library, 2 = public databases); LysoPE, lysophosphatidylethanolamine; SD, standard deviation; VIP, variable importance to projection (from PLS-DA); p, p-value from Welch's ANOVA or t-test; Mean ion abundances are shown
