## Supplementary material for "Physiological basis underlying antidepressant-induced activation of TrkB receptors": Table S1. Statistics

**Figure 1. N2O evokes a sleep-like of state during which TrkB signaling becomes upregulated.**

**A**

| Treatment | Treatment dose | | n |
|---|---|---|---|
| Medetomidine | 0,3 | mg/kg ip | 8 |
| N2O | 65 | % | 4 |
| Saline | | | 4 |

**B** **N2O Recovery time-dependent effects**

**N2O-25+0 min recovery**

**pTrkB**

| | |
|---|---|
| Mann Whitney test | |
| P value | 0,2949 |
| Exact or approximate P value? | Exact |
| P value summary | ns |
| Significantly different (P < 0.05)? | No |
| One- or two-tailed P value? | Two-tailed |
| Sum of ranks in column A,B | 34 , 57 |
| Mann-Whitney U | 13 |
| Difference between medians | |
| Median of column A | 96,55, n=6 |
| Median of column B | 119,3, n=7 |
| Difference: Actual | 22,74 |
| Difference: Hodges-Lehmann | 18,41 |

**pGSKbeta**

| | |
|---|---|
| Mann Whitney test | |
| P value | >0,9999 |
| Exact or approximate P value? | Exact |
| P value summary | ns |
| Significantly different (P < 0.05)? | No |
| One- or two-tailed P value? | Two-tailed |
| Sum of ranks in column A,B | 68 , 68 |
| Mann-Whitney U | 32 |
| Difference between medians | |
| Median of column A | 93,05, n=8 |
| Median of column B | 97,12, n=8 |
| Difference: Actual | 4,07 |
| Difference: Hodges-Lehmann | 0,1264 |

**pp70s6k**

| | |
|---|---|
| Mann Whitney test | |
| P value | 0,6454 |
| Exact or approximate P value? | Exact |
| P value summary | ns |
| Significantly different (P < 0.05)? | No |
| One- or two-tailed P value? | Two-tailed |
| Sum of ranks in column A,B | 63 , 73 |
| Mann-Whitney U | 27 |
| Difference between medians | |
| Median of column A | 82,07, n=8 |
| Median of column B | 89,45, n=8 |
| Difference: Actual | 7,387 |
| Difference: Hodges-Lehmann | 18,68 |

**N2O-20+5 min recovery**

**pTrkB**

| | |
|---|---|
| Mann Whitney test | |
| P value | 0,4908 |
| Exact or approximate P value? | Exact |
| P value summary | ns |
| Significantly different (P < 0.05)? | No |
| One- or two-tailed P value? | Two-tailed |
| Sum of ranks in column A,C | 39 , 66 |
| Mann-Whitney U | 18 |

| | |
|---|---|
| Difference between medians | |
| Median of column A | 96,55, n=6 |
| Median of column C | 104,7, n=8 |
| Difference: Actual | 8,155 |
| Difference: Hodges-Lehmann | 3,795 |

**pGSKbeta**

| | |
|---|---|
| Mann Whitney test | |
| P value | 0,0152 |
| Exact or approximate P value? | Exact |
| P value summary | * |
| Significantly different (P < 0.05)? | Yes |
| One- or two-tailed P value? | Two-tailed |
| Sum of ranks in column A,C | 47 , 106 |
| Mann-Whitney U | 11 |
| Difference between medians | |
| Median of column A | 93,05, n=8 |
| Median of column C | 179,1, n=9 |
| Difference: Actual | 86,04 |
| Difference: Hodges-Lehmann | 88,08 |

**pp70s6k**

| | |
|---|---|
| Mann Whitney test | |
| P value | 0,1672 |
| Exact or approximate P value? | Exact |
| P value summary | ns |
| Significantly different (P < 0.05)? | No |
| One- or two-tailed P value? | Two-tailed |
| Sum of ranks in column A,C | 57 , 96 |
| Mann-Whitney U | 21 |
| Difference between medians | |
| Median of column A | 82,07, n=8 |
| Median of column C | 163,3, n=9 |
| Difference: Actual | 81,24 |
| Difference: Hodges-Lehmann | 80,59 |

**N2O 20 min + 15 min recovery**

**pTrkB**

| | |
|---|---|
| Mann Whitney test | |
| P value | 0,053 |
| Exact or approximate P value? | Exact |
| P value summary | * |
| Significantly different (P < 0.05)? | No |
| One- or two-tailed P value? | Two-tailed |
| Sum of ranks in column A,B | 37 , 68 |
| Mann-Whitney U | 9 |
| Difference between medians | |
| Median of column A | 97,31, n=7 |
| Median of column B | 135,0, n=7 |
| Difference: Actual | 37,64 |
| Difference: Hodges-Lehmann | 42,03 |

**pGSK3b**

| | |
|---|---|
| Mann Whitney test | |
| P value | 0,0379 |
| Exact or approximate P value? | Exact |
| P value summary | * |
| Significantly different (P < 0.05)? | Yes |
| One- or two-tailed P value? | Two-tailed |
| Sum of ranks in column A,B | 36 , 69 |
| Mann-Whitney U | 8 |
| Difference between medians | |
| Median of column A | 108,8, n=7 |
| Median of column B | 178,6, n=7 |
| Difference: Actual | 69,84 |
| Difference: Hodges-Lehmann | 97,41 |

**pp70s6k**

| | |
|---|---|
| Mann Whitney test | |
| P value | 0,0175 |
| Exact or approximate P value? | Exact |
| P value summary | * |
| Significantly different (P < 0.05)? | Yes |
| One- or two-tailed P value? | Two-tailed |
| Sum of ranks in column A,B | 34 , 71 |
| Mann-Whitney U | 6 |
| Difference between medians | |
| Median of column A | 98,49, n=7 |
| Median of column B | 204,8, n=7 |
| Difference: Actual | 106,3 |
| Difference: Hodges-Lehmann | 106,3 |

**C** **N2O Locomotor Activity**

| Treatment | Treatment dose | | n |
|---|---|---|---|
| N2O | 65 | % | 10 |
| SHAM | | | 10 |

**During N2O - AUC**

| | | |
|---|---|---|
| **Unpaired t test** | | |
| P value | <0,0001 | |
| P value su | **** | |
| Significantl | Yes | |
| One- or tw | Two-tailed | |
| t, df | t=6,077, df=18 | |
| **How big is the difference?** | | |
| Mean of cc | | 5299 |
| Mean of cc | | 6414 |
| Difference | 1115 ± 183,5 | |
| 95% confic | 729,5 to 1500 | |
| R squared | | 0,6723 |
| **F test to compare variances** | | |
| F, DFn, Df | 1,062, 9, 9 | |
| P value | | 0,9297 |
| P value su | ns | |
| Significantl | No | |

**After N2O - AUC**

| | | |
|---|---|---|
| **Unpaired t test** | | |
| P value | <0,0001 | |
| P value su | **** | |
| Significantl | Yes | |
| One- or tw | Two-tailed | |
| t, df | t=12,43, df=18 | |
| **How big is the difference?** | | |
| Mean of cc | | 31536 |
| Mean of cc | | 5027 |
| Difference | -26509 ± 2133 | |
| 95% confic | -30991 to -22027 | |
| R squared | | 0,8956 |
| **F test to compare variances** | | |
| F, DFn, Df | 1,612, 9, 9 | |
| P value | | 0,4879 |
| P value su | ns | |
| Significantl | No | |

**D** **Functional autoradiography**

| | | |
|---|---|---|
| MED | 0.03 | 6 |
| VEH | | 5 * |
| N2O | 65 % | 7 |
| SHAM | | 7 |

*Note: One subject discarded from the group due to insufficient radiolabel administration based on detections and plasma analysis

**MED**

**Cingulate Cortex**

| | |
|---|---|
| **Mann Whitney test** | |
| P value | 0,0043 |
| Exact or approximate P value? | Exact |
| P value summary | ** |
| Significantly different (P < 0.05)? | Yes |
| One- or two-tailed P value? | Two-tailed |
| Sum of ranks in column A,B | 45 , 21 |
| Mann-Whitney U | 0 |
| **Difference between medians** | |
| Median of column A | 226,5, n=5 |
| Median of column B | 192,7, n=6 |
| Difference: Actual | -33,81 |
| Difference: Hodges-Lehmann | -39,6 |

**Retrosplenial cortex**

| | |
|---|---|
| **Mann Whitney test** | |
| P value | 0,0087 |
| Exact or approximate P value? | Exact |
| P value summary | ** |
| Significantly different (P < 0.05)? | Yes |
| One- or two-tailed P value? | Two-tailed |
| Sum of ranks in column A,B | 44 , 22 |
| Mann-Whitney U | 1 |
| **Difference between medians** | |
| Median of column A | 220,8, n=5 |
| Median of column B | 169,7, n=6 |
| Difference: Actual | -51,07 |
| Difference: Hodges-Lehmann | -52,63 |

**Laterodorsal thalamus**

| | |
|---|---|
| **Mann Whitney test** | |
| P value | 0,0087 |
| Exact or approximate P value? | Exact |
| P value summary | ** |
| Significantly different (P < 0.05)? | Yes |
| One- or two-tailed P value? | Two-tailed |
| Sum of ranks in column A,B | 44 , 22 |
| Mann-Whitney U | 1 |
| **Difference between medians** | |
| Median of column A | 179,3, n=5 |
| Median of column B | 156,3, n=6 |
| Difference: Actual | -22,98 |
| Difference: Hodges-Lehmann | -42,23 |

**Primary motor cortex**

| | |
|---|---|
| **Mann Whitney test** | |
| P value | 0,0173 |
| Exact or approximate P value? | Exact |
| P value summary | * |
| Significantly different (P < 0.05)? | Yes |
| One- or two-tailed P value? | Two-tailed |
| Sum of ranks in column A,B | 43 , 23 |
| Mann-Whitney U | 2 |
| **Difference between medians** | |
| Median of column A | 215,2, n=5 |
| Median of column B | 183,9, n=6 |
| Difference: Actual | -31,31 |
| Difference: Hodges-Lehmann | -34,4 |

**Hippocampus**

| | |
|---|---|
| **Mann Whitney test** | |
| P value | 0,1775 |
| Exact or approximate P value? | Exact |
| P value summary | ns |
| Significantly different (P < 0.05)? | No |
| One- or two-tailed P value? | Two-tailed |
| Sum of ranks in column A,B | 38 , 28 |
| Mann-Whitney U | 7 |

| | |
|---|---|
| **Difference between medians** | |
| Median of column A | 157,6, n=5 |
| Median of column B | 145,7, n=6 |
| Difference: Actual | -11,84 |
| Difference: Hodges-Lehmann | -16,72 |

**Amygdala**

| | |
|---|---|
| **Mann Whitney test** | |
| P value | 0,9307 |
| Exact or approximate P value? | Exact |
| P value summary | ns |
| Significantly different (P < 0.05)? | No |
| One- or two-tailed P value? | Two-tailed |
| Sum of ranks in column A,B | 29 , 37 |
| Mann-Whitney U | 14 |
| **Difference between medians** | |
| Median of column A | 151,7, n=5 |
| Median of column B | 152,4, n=6 |
| Difference: Actual | 0,7879 |
| Difference: Hodges-Lehmann | 2,457 |

**N2O**

**Cingulate Cortex**

| | |
|---|---|
| **Mann Whitney test** | |
| P value | 0,3829 |
| Exact or approximate P value? | Exact |
| P value summary | ns |
| Significantly different (P < 0.05)? | No |
| One- or two-tailed P value? | Two-tailed |
| Sum of ranks in column A,B | 60 , 45 |
| Mann-Whitney U | 17 |
| **Difference between medians** | |
| Median of column A | 249,5, n=7 |
| Median of column B | 243,5, n=7 |
| Difference: Actual | -6,036 |
| Difference: Hodges-Lehmann | -13,53 |

**Retrosplenial cortex**

| | |
|---|---|
| **Mann Whitney test** | |
| P value | 0,0175 |
| Exact or approximate P value? | Exact |
| P value summary | * |
| Significantly different (P < 0.05)? | Yes |
| One- or two-tailed P value? | Two-tailed |
| Sum of ranks in column A,B | 71 , 34 |
| Mann-Whitney U | 6 |
| **Difference between medians** | |
| Median of column A | 256,1, n=7 |
| Median of column B | 201,8, n=7 |
| Difference: Actual | -54,35 |
| Difference: Hodges-Lehmann | -63,52 |

**Laterodorsal thalamus**

| | |
|---|---|
| **Mann Whitney test** | |
| P value | 0,0379 |
| Exact or approximate P value? | Exact |
| P value summary | * |
| Significantly different (P < 0.05)? | Yes |
| One- or two-tailed P value? | Two-tailed |
| Sum of ranks in column A,B | 69 , 36 |
| Mann-Whitney U | 8 |
| **Difference between medians** | |
| Median of column A | 228,8, n=7 |
| Median of column B | 183,1, n=7 |
| Difference: Actual | -45,61 |
| Difference: Hodges-Lehmann | -52,63 |

**Primary motor cortex**

**Mann Whitney test**

| | |
|---|---|
| P value | 0,0973 |
| Exact or approximate P value? | Exact |
| P value summary | ns |
| Significantly different (P < 0.05)? | No |
| One- or two-tailed P value? | Two-tailed |
| Sum of ranks in column A,B | 66 , 39 |
| Mann-Whitney U | 11 |

**Difference between medians**

| | |
|---|---|
| Median of column A | 214,5, n=7 |
| Median of column B | 206,3, n=7 |
| Difference: Actual | -8,147 |
| Difference: Hodges-Lehmann | -14,53 |

**Hippocampus**

**Mann Whitney test**

| | |
|---|---|
| P value | 0,0728 |
| Exact or approximate P value? | Exact |
| P value summary | ns |
| Significantly different (P < 0.05)? | No |
| One- or two-tailed P value? | Two-tailed |
| Sum of ranks in column A,B | 67 , 38 |
| Mann-Whitney U | 10 |

**Difference between medians**

| | |
|---|---|
| Median of column A | 178,0, n=7 |
| Median of column B | 153,2, n=7 |
| Difference: Actual | -24,82 |
| Difference: Hodges-Lehmann | -27,96 |

**Amygdala**

**Mann Whitney test**

| | |
|---|---|
| P value | 0,1649 |
| Exact or approximate P value? | Exact |
| P value summary | ns |
| Significantly different (P < 0.05)? | No |
| One- or two-tailed P value? | Two-tailed |
| Sum of ranks in column A,B | 64 , 41 |
| Mann-Whitney U | 13 |

**Difference between medians**

| | |
|---|---|
| Median of column A | 171,9, n=7 |
| Median of column B | 148,2, n=7 |
| Difference: Actual | -23,71 |
| Difference: Hodges-Lehmann | -21,8 |

**Figure 2. TrkB signaling is increased during physiological deep sleep.**

A

| Treatment Duration | n | |
|---|---|---|
| 6h SD + 1.5h recovery sleep, NREM sleep spectral power | 4 | |
| 24-hour baseline recording, NREM sleep spectral power | 4 | (within-subject control) |

| | | AUC | 95 % CI | Std. error | t-test p vs Baseline |
|---|---|---|---|---|---|
| Delta | Base | 26,87 | 23,97 to 29,77 | 1,478 | |
| | SDR | 36,81 | 32,18 to 41,44 | 2,362 | 0,0118 |
| Theta | Base | 19 | 17,43 to 20,57 | 0,8013 | |
| | SDR | 24,7 | 21,90 to 27,50 | 1,427 | 0,0145 |
| Alpha | Base | 16,02 | 14,40 to 17,64 | 0,8256 | |
| | SDR | 19,81 | 17,61 to 22,01 | 1,122 | 0,0346 |
| Beta | Base | 7,056 | 6,299 to 7,814 | 0,3866 | |
| | SDR | 8,763 | 7,575 to 9,950 | 0,6061 | 0,0552 |

**B** **SD**

**pTrkB**

| **Mann Whitney test** | |
|---|---|
| P value | 0,8125 |
| Exact or approximate P value? | Exact |
| P value summary | ns |
| Significantly different (P < 0.05)? | No |
| One- or two-tailed P value? | Two-tailed |
| Sum of ranks in column A,B | 60 , 93 |
| Mann-Whitney U | 32 |
| **Difference between medians** | |
| Median of column A | 107,4, n=7 |
| Median of column B | 98,85, n=10 |
| Difference: Actual | -8,54 |
| Difference: Hodges-Lehmann | 1,766 |

**pGSK3b**

| **Mann Whitney test** | |
|---|---|
| P value | 0,2698 |
| Exact or approximate P value? | Exact |
| P value summary | ns |
| Significantly different (P < 0.05)? | No |
| One- or two-tailed P value? | Two-tailed |
| Sum of ranks in column A,B | 75 , 78 |
| Mann-Whitney U | 23 |
| **Difference between medians** | |
| Median of column A | 99,88, n=7 |
| Median of column B | 86,60, n=10 |
| Difference: Actual | -13,28 |
| Difference: Hodges-Lehmann | -12,43 |

**pp70s6k**

| **Mann Whitney test** | |
|---|---|
| P value | 0,3148 |
| Exact or approximate P value? | Exact |

| | |
|---|---|
| P value summary | ns |
| Significantly different (P < 0.05)? | No |
| One- or two-tailed P value? | Two-tailed |
| Sum of ranks in column A,B | 74 , 79 |
| Mann-Whitney U | 24 |

**Difference between medians**

| | |
|---|---|
| Median of column A | 124,1, n=7 |
| Median of column B | 90,75, n=10 |
| Difference: Actual | -33,34 |
| Difference: Hodges-Lehmann | -23,21 |

**SD+Recovery**

**pTrkB**

**Mann Whitney test**

| | |
|---|---|
| P value | 0,0524 |
| Exact or approximate P value? | Exact |
| P value summary | * |
| Significantly different (P < 0.05)? | No |
| One- or two-tailed P value? | Two-tailed |
| Sum of ranks in column A,B | 79 , 131 |
| Mann-Whitney U | 24 |

Difference between medians

| | |
|---|---|
| Median of column A | 95,88, n=10 |
| Median of column B | 125,6, n=10 |
| Difference: Actual | 29,74 |
| Difference: Hodges-Lehmann | 25,3 |

**pGSK3b**

**Mann Whitney test**

| | |
|---|---|
| P value | 0,0147 |
| Exact or approximate P value? | Exact |
| P value summary | * |
| Significantly different (P < 0.05)? | Yes |
| One- or two-tailed P value? | Two-tailed |
| Sum of ranks in column A,B | 73 , 137 |
| Mann-Whitney U | 18 |

**Difference between medians**

| | |
|---|---|
| Median of column A | 99,29, n=10 |
| Median of column B | 144,7, n=10 |
| Difference: Actual | 45,38 |
| Difference: Hodges-Lehmann | 38,59 |

**pp70s6k**

**Mann Whitney test**

| | |
|---|---|
| P value | 0,0433 |
| Exact or approximate P value? | Exact |
| P value summary | * |
| Significantly different (P < 0.05)? | Yes |
| One- or two-tailed P value? | Two-tailed |
| Sum of ranks in column A,B | 78 , 132 |
| Mann-Whitney U | 23 |

**Difference between medians**

| | |
|---|---|
| Median of column A | 87,09, n=10 |
| Median of column B | 173,0, n=10 |

| | |
|---|---|
| Difference: Actual | 85,96 |
| Difference: Hodges-Lehmann | 76,18 |

| | | | | | | | | | | | | | | | | | | | | | | | | | |
|---|---|---|---|---|---|---|---|---|---|---|---|---|---|---|---|---|---|---|---|---|---|---|---|---|---|
| **pTrkB** | **Treatment** | **Mianserin** | **Hydroxyzine** | **Atipamezole** | **Medetomidine** | **Control** | **Amitriptyline** | **Fluoxetine** | **Control** | **Mirtazapine** | **Paroxetine** | **Duloxetine** | **Control** | **Amphetamine** | **Control** | **GHB** | **Control** | **Clozapine** | **Control** | **Chlorpromazin** | **Control** | **Isoflurane** | **Sevoflurane** | **Urethane** | **Control** |
| | **Dose (mg/kg)** | 20 | 50 | 1 | 0,05 | | 20 | 20 | | 20 | 10 | 20 | | 10 | | 275 | | 6 | | 10 | | 4%/2% | 6%/4.5% | 2000 | |
| | **Route** | sc | sc | sc | sc | sc | sc | sc | sc | sc | sc | sc | sc | ip | | ip | ip | ip | ip | ip | ip | | | | |
| | **n** | 10 | 10 | 10 | 10 | 10 | 6 | 6 | 6 | 6 | 6 | 6 | 6 | 4 | 5 | 5 | 6 | 6 | 6 | 6 | 6 | 6 | 6 | 6 | 6 |
| | **Mann Whitney** | | | | | | | | | | | | | | | | | | | | | | | | |
| | **P value** | 0,0003 | 0,0016 | 0,5408 | <0,0001 | | 0,0152 | 0,0043 | | 0,2403 | 0,7381 | 0,8182 | | 0,5556 | | 0,0043 | | 0,026 | | 0,0087 | | 0,0022 | 0,0152 | 0,0286 | |
| | **Exact or approximate P value?** | Exact | Exact | Exact | Exact | | Exact | Exact | | Exact | Exact | Exact | | Exact | | Exact | | Exact | | Exact | | Exact | Exact | Exact | |
| | **P value summary** | *** | ** | ns | **** | | * | ** | | ns | ns | ns | | ns | | ** | | * | | ** | | ** | * | * | |
| | **Significantly different (P < 0.05)?** | Yes | Yes | No | Yes | | Yes | Yes | | No | No | No | | No | | Yes | | Yes | | Yes | | Yes | Yes | Yes | |
| | **One- or two-tailed P value?** | Two-tailed | Two-tailed | Two-tailed | Two-tailed | | Two-tailed | Two-tailed | | Two-tailed | Two-tailed | Two-tailed | | Two-tailed | | Two-tailed | | Two-tailed | | Two-tailed | | Two-tailed | Two-tailed | Two-tailed | |
| | **Sum of ranks in column A,B** | 61 , 149 | 65,50 , 144,5 | 113,5 , 96,50 | 57,50 , 152,5 | | 24 , 54 | 22 , 56 | | 31 , 47 | 36,50 , 41,50 | 41 , 37 | | 22 , 23 | | 21 , 45 | | 25 , 53 | | 23 , 55 | | 21 , 57 | 24 , 54 | 10 , 26 | |
| | **Mann-Whitney U** | 6 | 10,5 | 41,5 | 2,5 | | 3 | 1 | | 10 | 15,5 | 16 | | 7 | | 0 | | 4 | | 2 | | 0 | 3 | 0 | |
| | **Difference between medians** | | | | | | | | | | | | | | | | | | | | | | | | |
| | **Median of column A** | 100,5, n=10 | 100,5, n=10 | 100,5, n=10 | 100,5, n=10 | | 96,06, n=6 | 96,06, n=6 | | 92,50, n=6 | 92,50, n=6 | 92,50, n=6 | | 106,2, n=5 | | 103,6, n=6 | | 93,19, n=6 | | 103,7, n=6 | | 106,0, n=6 | 106,0, n=6 | 103,5, n=4 | |
| | **Median of column B** | 123,0, n=10 | 131,5, n=10 | 92,00, n=10 | 180,0, n=10 | | 160,6, n=6 | 167,9, n=6 | | 114,5, n=6 | 106,5, n=6 | 95,00, n=6 | | 113,4, n=4 | | 143,4, n=5 | | 179,8, n=6 | | 118,6, n=6 | | 170,8, n=6 | 168,4, n=6 | 216,2, n=4 | |
| | **Difference: Actual** | 22,5 | 31 | -8,5 | 79,5 | | 64,54 | 71,85 | | 22 | 14 | 2,5 | | 7,152 | | 39,84 | | 86,57 | | 14,95 | | 64,87 | 62,42 | 112,6 | |
| | **Difference: Hodges-Lehmann** | 25,5 | 32 | -3 | 79 | | 57,97 | 74,65 | | 22 | 14 | -8 | | 10,9 | | 45,26 | | 87,16 | | 17,71 | | 68,26 | 63,53 | 112,6 | |
| **pGSK3b** | **Treatment** | **Mianserin** | **Hydroxyzine** | **Atipamezole** | **Medetomidine** | **Control** | **Amitriptyline** | **Fluoxetine** | **Control** | **Mirtazapine** | **Paroxetine** | **Duloxetine** | **Control** | **Amphetamine** | **Control** | **GHB** | **Control** | **Clozapine** | **Control** | **Chlorpromazin** | **Control** | **Isoflurane** | **Sevoflurane** | **Urethane** | **Control** |
| | **Mann Whitney test** | | | | | | | | | | | | | | | | | | | | | | | | |
| | **P value** | 0,0012 | <0,0001 | 0,7817 | <0,0001 | | 0,0022 | 0,0022 | | 0,0087 | 0,1645 | 0,3745 | | 0,9048 | | 0,0087 | | 0,0411 | | 0,026 | | 0,0022 | 0,0022 | 0,019 | |
| | **Exact or approximate P value?** | Exact | Exact | Exact | Exact | | Exact | Exact | | Exact | Exact | Exact | | Exact | | Exact | | Exact | | Exact | | Exact | Exact | Exact | |
| | **P value summary** | ** | **** | ns | **** | | ** | ** | | ** | ns | ns | | ns | | ** | | * | | * | | ** | ** | * | |
| | **Significantly different (P < 0.05)?** | Yes | Yes | No | Yes | | Yes | Yes | | Yes | No | No | | No | | Yes | | Yes | | Yes | | Yes | Yes | Yes | |
| | **One- or two-tailed P value?** | Two-tailed | Two-tailed | Two-tailed | Two-tailed | | Two-tailed | Two-tailed | | Two-tailed | Two-tailed | Two-tailed | | Two-tailed | | Two-tailed | | Two-tailed | | Two-tailed | | Two-tailed | Two-tailed | Two-tailed | |
| | **Sum of ranks in column A,B** | 64,50 , 145,5 | 57 , 153 | 109 , 101 | 58 , 152 | | 21 , 57 | 21 , 57 | | 23 , 55 | 30 , 48 | [illegible]2,50 , 45,50 | | 24 , 21 | | 22 , 44 | | 26 , 52 | | 25 , 53 | | 21 , 57 | 21 , 57 | 11 , 44 | |
| | **Mann-Whitney U** | 9,5 | 2 | 46 | 3 | | 0 | 0 | | 2 | 9 | 11,5 | | 9 | | 1 | | 5 | | 4 | | 0 | 0 | 1 | |
| | **Difference between medians** | | | | | | | | | | | | | | | | | | | | | | | | |
| | **Median of column A** | 98,00, n=10 | 98,00, n=10 | 98,00, n=10 | 98,00, n=10 | | 99,94, n=6 | 99,94, n=6 | | 99,50, n=6 | 99,50, n=6 | 99,50, n=6 | | 91,14, n=5 | | 93,87, n=6 | | 93,33, n=6 | | 80,81, n=6 | | 103,7, n=6 | 103,7, n=6 | 55,94, n=4 | |
| | **Median of column B** | 137,0, n=10 | 146,0, n=10 | 97,50, n=10 | 246,0, n=10 | | 148,0, n=6 | 180,6, n=6 | | 120,5, n=6 | 122,0, n=6 | 106,5, n=6 | | 99,20, n=4 | | 347,6, n=5 | | 191,5, n=6 | | 308,8, n=6 | | 257,9, n=6 | 250,1, n=6 | 588,8, n=6 | |
| | **Difference: Actual** | 39 | 48 | -0,5 | 148 | | 48,1 | 80,64 | | 21 | 22,5 | 7 | | 8,062 | | 253,7 | | 98,15 | | 228 | | 154,2 | 146,5 | 532,9 | |
| | **Difference: Hodges-Lehmann** | 38 | 47 | -6,5 | 142 | | 48,03 | 80,56 | | 25 | 21 | 7,5 | | 5,1 | | 253,7 | | 81,42 | | 228 | | 156,2 | 153,2 | 444,8 | |
| **pp70s6k** | **Treatment** | **Mianserin** | **Hydroxyzine** | **Atipamezole** | **Medetomidine** | **Control** | **Amitriptyline** | **Fluoxetine** | **Control** | **Mirtazapine** | **Paroxetine** | **Duloxetine** | **Control** | **Amphetamine** | **Control** | **GHB** | **Control** | **Clozapine** | **Control** | **Chlorpromazin** | **Control** | **Isoflurane** | **Sevoflurane** | **Urethane** | **Control** |
| | **Mann Whitney test** | | | | | | | | | | | | | | | | | | | | | | | | |
| | **P value** | 0,0003 | <0,0001 | 0,123 | 0,0001 | | 0,0022 | 0,0022 | | 0,1212 | 0,8182 | 0,3939 | | 0,5556 | | 0,0043 | | 0,0152 | | 0,0411 | | 0,0022 | 0,0022 | 0,0095 | |
| | **Exact or approximate P value?** | Exact | Exact | Exact | Exact | | Exact | Exact | | Exact | Exact | Exact | | Exact | | Exact | | Exact | | Exact | | Exact | Exact | Exact | |
| | **P value summary** | *** | **** | ns | *** | | ** | ** | | ns | ns | ns | | ns | | ** | | * | | * | | ** | ** | ** | |
| | **Significantly different (P < 0.05)?** | Yes | Yes | No | Yes | | Yes | Yes | | No | No | No | | No | | Yes | | Yes | | Yes | | Yes | Yes | Yes | |
| | **One- or two-tailed P value?** | Two-tailed | Two-tailed | Two-tailed | Two-tailed | | Two-tailed | Two-tailed | | Two-tailed | Two-tailed | Two-tailed | | Two-tailed | | Two-tailed | | Two-tailed | | Two-tailed | | Two-tailed | Two-tailed | Two-tailed | |
| | **Sum of ranks in column A,B** | 61 , 149 | 56 , 154 | 126 , 84 | 59 , 151 | | 21 , 57 | 21 , 57 | | 29 , 49 | 37 , 41 | 33 , 45 | | 22 , 23 | | 21 , 45 | | 24 , 54 | | 26 , 52 | | 21 , 57 | 21 , 57 | 10 , 45 | |
| | **Mann-Whitney U** | 6 | 1 | 29 | 4 | | 0 | 0 | | 8 | 16 | 12 | | 7 | | 0 | | 3 | | 5 | | 0 | 0 | 0 | |
| | **Difference between medians** | | | | | | | | | | | | | | | | | | | | | | | | |
| | **Median of column A** | 96,00, n=10 | 96,00, n=10 | 96,00, n=10 | 96,00, n=10 | | 84,43, n=6 | 84,43, n=6 | | 88,00, n=6 | 88,00, n=6 | 88,00, n=6 | | 100,4, n=5 | | 101,1, n=6 | | 105,0, n=6 | | 69,77, n=6 | | 81,27, n=6 | 81,27, n=6 | 100,0, n=4 | |
| | **Median of column B** | 212,0, n=10 | 180,0, n=10 | 76,50, n=10 | 277,5, n=10 | | 240,2, n=6 | 399,4, n=6 | | 146,5, n=6 | 111,5, n=6 | 109,0, n=6 | | 107,6, n=4 | | 641,1, n=5 | | 231,1, n=6 | | 209,8, n=6 | | 948,6, n=6 | 605,6, n=6 | 1490, n=6 | |
| | **Difference: Actual** | 116 | 84 | -19,5 | 181,5 | | 155,8 | 315 | | 58,5 | 23,5 | 21 | | 7,194 | | 540 | | 126 | | 140 | | 867,4 | 524,4 | 1390 | |
| | **Difference: Hodges-Lehmann** | 101,5 | 89,5 | -22,5 | 175,5 | | 152,9 | 294,3 | | 48,5 | 7 | 8 | | 10,41 | | 540 | | 144,4 | | 124,4 | | 842,8 | 492,7 | 1390 | |

**Figure 4: Metabolism**

**A** **1h**

| **Treatment** | **Treatment dose** | **n** | | **p** | | **test** | | **p** | |
|---|---|---|---|---|---|---|---|---|---|
| Isoflurane | 1,5 | 6 | 0,0449 | 0,0024 | 0,0386 | Kruskal-wallis | * | ** | * |
| Flurothyl | seizure | 6 | 0,002 | 0,0386 | 0,0024 | Kruskal-wallis | ** | * | ** |
| SHAM | Room air | 6 | | | | | | | |

**pTrkB**

| | |
|---|---|
| Kruskal-Wallis test | |
| P value | 0,0001 |
| Exact or approximate P \ | Exact |
| P value summary | *** |
| Do the medians vary sigı | Yes |
| Number of groups | 3 |
| Kruskal-Wallis statistic | 12,12 |

| Dunn's multiple compari | Mean rank | Significant | Summary | Adjusted P | Z |
|---|---|---|---|---|---|
| VEH vs. ISO | -7,5 | Yes | * | 0,0449 | 2,433 |
| VEH vs. FLURO | -10,5 | Yes | ** | 0,002 | 3,407 |
| ISO vs. FLURO | -3 | No | ns | 0,9912 | 0,9733 |

**pGSK3beta**

| | |
|---|---|
| Kruskal-Wallis test | |
| P value | 0,0001 |
| Exact or approximate P \ | Exact |
| P value summary | *** |
| Do the medians vary sigı | Yes |
| Number of groups | 3 |

| Dunn's multiple compari | Mean rank | Significant | Summary | Adjusted P | Z |
|---|---|---|---|---|---|
| VEH vs. ISO | -10,33 | Yes | ** | 0,0024 | 3,353 |
| VEH vs. FLURO | -7,667 | Yes | * | 0,0386 | 2,487 |
| ISO vs. FLURO | 2,667 | No | ns | >0,9999 | 0,8652 |

**pp70s6k**

| | |
|---|---|
| Kruskal-Wallis test | |
| P value | 0,0001 |
| Exact or approximate P \ | Exact |
| P value summary | *** |
| Do the medians vary sigı | Yes |
| Number of groups | 3 |

| Dunn's multiple compari | Mean rank | Significant | Summary | Adjusted P | Z |
|---|---|---|---|---|---|
| VEH vs. ISO | -7,667 | Yes | * | 0,0386 | 2,487 |
| VEH vs. FLURO | -10,33 | Yes | ** | 0,0024 | 3,353 |
| ISO vs. FLURO | -2,667 | No | ns | >0,9999 | 0,8652 |

**Immediate**

| | **Dose** | **n** |
|---|---|---|
| **Flurothyl** | seizure | 7 |
| **SHAM** | room air | 7 |

**pTrkB**

| | |
|---|---|
| Mann Whitney test | |
| P value | 0,535 |
| Exact or approximate P \ | Exact |
| P value summary | ns |
| Significantly different (P < | No |
| One- or two-tailed P valu | Two-tailed |
| Sum of ranks in column / | 47 , 58 |
| Mann-Whitney U | 19 |
| Difference between medians | |
| Median of column A | 97,82, n=7 |
| Median of column B | 111,1, n=7 |
| Difference: Actual | 13,29 |
| Difference: Hodges-Lehr | 5,809 |

**pGSK3beta**

| | |
|---|---|
| Mann Whitney test | |
| P value | 0,9015 |
| Exact or approximate P \ | Exact |
| P value summary | ns |

| | |
|---|---|
| Significantly different (P < | No |
| One- or two-tailed P valu | Two-tailed |
| Sum of ranks in column A | 51 , 54 |
| Mann-Whitney U | 23 |
| Difference between medians | |
| Median of column A | 113,2, n=7 |
| Median of column B | 106,7, n=7 |
| Difference: Actual | -6,478 |
| Difference: Hodges-Lehr | 1,015 |

**pp70s6k**

| | |
|---|---|
| Mann Whitney test | |
| P value | 0,8048 |
| Exact or approximate P v | Exact |
| P value summary | ns |
| Significantly different (P < | No |
| One- or two-tailed P valu | Two-tailed |
| Sum of ranks in column A | 50 , 55 |
| Mann-Whitney U | 22 |
| Difference between medians | |
| Median of column A | 90,76, n=7 |
| Median of column B | 96,23, n=7 |
| Difference: Actual | 5,467 |
| Difference: Hodges-Lehr | 0,8656 |

**B**

**1h**
**Treatment**
Isoflurane
Flurothyl
SHAM

**C**

| | | | |
|---|---|---|---|
| EEG | ATP | 125 mg/kg, ip | within-subject control |
| | Control | Saline | |
| **WB** | ATP | 125 mg/kg, ip | |
| | Control | Saline | |

**pTrkB**

| | |
|---|---|
| Mann Whitney test | |
| P value | 0,0556 |
| Exact or approximate P v | Exact |
| P value summary | ns |
| Significantly different (P < | No |
| One- or two-tailed P valu | Two-tailed |
| Sum of ranks in column A | 18 , 37 |
| Mann-Whitney U | 3 |
| Difference between medians | |
| Median of column A | 97,25, n=5 |
| Median of column B | 131,0, n=5 |
| Difference: Actual | 33,74 |
| Difference: Hodges-Lehr | 39,35 |

**pGSK3b**

| | |
|---|---|
| Mann Whitney test | |
| P value | 0,0317 |
| Exact or approximate P v | Exact |
| P value summary | * |
| Significantly different (P < | Yes |
| One- or two-tailed P valu | Two-tailed |
| Sum of ranks in column A | 17 , 38 |
| Mann-Whitney U | 2 |
| Difference between medians | |
| Median of column A | 80,08, n=5 |
| Median of column B | 219,4, n=5 |
| Difference: Actual | 139,3 |
| Difference: Hodges-Lehr | 102,1 |

**pp70**

| | |
|---|---|
| Mann Whitney test | |
| P value | 0,0159 |

| | |
|---|---|
| Exact or approximate P v | Exact |
| P value summary | * |
| Significantly different (P < | Yes |
| One- or two-tailed P valu | Two-tailed |
| Sum of ranks in column A | 16 , 39 |
| Mann-Whitney U | 1 |
| Difference between medians | |
| Median of column A | 93,73, n=5 |
| Median of column B | 250,5, n=5 |
| Difference: Actual | 156,7 |
| Difference: Hodges-Lehr | 156,7 |

**D**

| | | |
|---|---|---|
| **VEH** | 20 ml/kg | 8 |
| **2-DG+MA** | 1g+1200µmol, 20 ml/kg | 8 |

**WB**

**pTrkB**

| | |
|---|---|
| Mann Whitney test | |
| P value | 0,0019 |
| Exact or approximate P v | Exact |
| P value summary | ** |
| Significantly different (P < | Yes |
| One- or two-tailed P valu | Two-tailed |
| Sum of ranks in column A | 40 , 96 |
| Mann-Whitney U | 4 |
| Difference between medians | |
| Median of column A | 103,9, n=8 |
| Median of column B | 151,5, n=8 |
| Difference: Actual | 47,57 |
| Difference: Hodges-Lehr | 52,13 |

**pGSK3beta**

| | |
|---|---|
| Mann Whitney test | |
| P value | 0,0003 |
| Exact or approximate P v | Exact |
| P value summary | *** |
| Significantly different (P < | Yes |
| One- or two-tailed P valu | Two-tailed |
| Sum of ranks in column A | 37 , 99 |
| Mann-Whitney U | 1 |
| Difference between medians | |
| Median of column A | 102,7, n=8 |
| Median of column B | 210,7, n=8 |
| Difference: Actual | 108 |
| Difference: Hodges-Lehr | 110,9 |

**pp70s6k**

| | |
|---|---|
| Mann Whitney test | |
| P value | 0,0002 |
| Exact or approximate P v | Exact |
| P value summary | *** |
| Significantly different (P < | Yes |
| One- or two-tailed P valu | Two-tailed |
| Sum of ranks in column A | 36 , 100 |
| Mann-Whitney U | 0 |
| Difference between medians | |
| Median of column A | 104,9, n=8 |
| Median of column B | 246,4, n=8 |
| Difference: Actual | 141,5 |
| Difference: Hodges-Lehr | 145,7 |

**Locomotor activity**

| | | |
|---|---|---|
| **VEH** | 20 ml/kg | 8 |
| **2-DG+MA** | 1g+1200µmol, 20 ml/kg | 8 |

**Test details**

| | |
|---|---|
| Test name | Unpaired t test of time bin vs VEH |
| Variance a | Individual variance for each row |
| Multiple co | False Discovery Rate (FDR) |

Method Two-stage step-up (Benjamini, Krieger, and Yekutieli)
Desired FI 1,00%

| time | P value | lean of VE | an of 2DG | Difference | of differer | t ratio | df | q value | p |
|---|---|---|---|---|---|---|---|---|---|
| 5 | 0,000022 | 845,5 | 486,5 | 358,9 | 57,51 | 6,242 | 14 | 0,000002 | **** |
| 10 | <0,000001 | 766 | 57,27 | 708,7 | 39,09 | 18,13 | 14 | <0,000001 | **** |
| 15 | <0,000001 | 829,9 | 11,8 | 818,1 | 84,76 | 9,651 | 14 | <0,000001 | **** |
| 20 | 0,000012 | 641,6 | 0 | 641,6 | 96,92 | 6,62 | 14 | 0,000001 | **** |
| 25 | <0,000001 | 718,4 | 7,335 | 711,1 | 67,88 | 10,48 | 14 | <0,000001 | **** |
| 30 | <0,000001 | 667 | 15,82 | 651,2 | 72,9 | 8,932 | 14 | <0,000001 | **** |
| 35 | 0,000006 | 632,8 | 23,68 | 609,1 | 87,25 | 6,982 | 14 | <0,000001 | **** |
| 40 | 0,000003 | 620 | 11,12 | 608,9 | 81,69 | 7,454 | 14 | <0,000001 | **** |
| 45 | 0,000006 | 479,8 | 2,421 | 477,4 | 67,94 | 7,027 | 14 | <0,000001 | **** |
| 50 | 0,000223 | 485,1 | 1,894 | 483,2 | 98,05 | 4,928 | 14 | 0,000022 | *** |
| 55 | 0,000278 | 380,7 | 1,426 | 379,3 | 78,88 | 4,808 | 14 | 0,000026 | *** |
| 60 | 0,010978 | 369,4 | 0,5953 | 368,8 | 125,9 | 2,93 | 14 | 0,000924 | *** |

**Figure 5: Temperature-dependence of drug-induced TrkB signaling**

**A** Amitriptyline + Warming, body temperature

| | dose | n |
|---|---|---|
| SAL+RT | | 8 |
| AMI+RT | 20 mg/kg | 7 |
| SAL+WARM | | 7 |
| AMI+WARI | 20 mg/kg | 8 |

**pTrkB**

**SAL-RT vs AMI-RT**

| | |
|---|---|
| Column B | AMI-RT |
| vs. | vs, |
| Column A | SAL-RT |
| Mann Whitney test | |
| P value | 0,0111 |
| Exact or approximate P value? | Exact |
| P value summary | * |
| Significantly different (P < 0.05)? | Yes |
| One- or two-tailed P value? | Two-tailed |
| Sum of ranks in column A,B | 33 , 72 |
| Mann-Whitney U | 5 |
| Difference between medians | |
| Median of column A | 105,0, n=7 |
| Median of column B | 138,9, n=7 |
| Difference: Actual | 33,85 |
| Difference: Hodges-Lehmann | 45,75 |

**SAL-RT vs AMI-W**

| | |
|---|---|
| Column D | AMI-WARM |
| vs. | vs, |
| Column C | SAL-RT |
| Mann Whitney test | |
| P value | >0,9999 |
| Exact or approximate P value? | Exact |
| P value summary | ns |
| Significantly different (P < 0.05)? | No |
| One- or two-tailed P value? | Two-tailed |
| Sum of ranks in column C,D | 52 , 53 |
| Mann-Whitney U | 24 |
| Difference between medians | |
| Median of column C | 93,38, n=7 |
| Median of column D | 94,21, n=7 |
| Difference: Actual | 0,8228 |
| Difference: Hodges-Lehmann | 0,4523 |

**SAL-RT vs SAL-WARM**

| | |
|---|---|
| Column F | SAL-WARM |
| vs. | vs, |
| Column E | SAL-RT |
| Mann Whitney test | |
| P value | >0,9999 |
| Exact or approximate P value? | Exact |
| P value summary | ns |
| Significantly different (P < 0.05)? | No |
| One- or two-tailed P value? | Two-tailed |
| Sum of ranks in column E,F | 52 , 53 |
| Mann-Whitney U | 24 |
| Difference between medians | |
| Median of column E | 93,38, n=7 |
| Median of column F | 94,21, n=7 |
| Difference: Actual | 0,8228 |
| Difference: Hodges-Lehmann | 0,4523 |

**pGSK3beta**

**SAL-RT vs AMI-RT**

| | |
|---|---|
| Column B | AMI-RT |
| vs. | vs, |
| Column A | SAL-RT |
| Mann Whitney test | |
| P value | 0,0111 |
| Exact or approximate P value? | Exact |
| P value summary | * |
| Significantly different (P < 0.05)? | Yes |
| One- or two-tailed P value? | Two-tailed |
| Sum of ranks in column A,B | 33 , 72 |
| Mann-Whitney U | 5 |
| Difference between medians | |
| Median of column A | 73,92, n=7 |
| Median of column B | 208,1, n=7 |
| Difference: Actual | 134,1 |
| Difference: Hodges-Lehmann | 119,4 |

**SAL-RT vs AMI-W**

| | |
|---|---|
| Column D | AMI-WARM |
| vs. | vs, |
| Column C | SAL-RT |
| Mann Whitney test | |
| P value | 0,2086 |
| Exact or approximate P value? | Exact |
| P value summary | ns |
| Significantly different (P < 0.05)? | No |
| One- or two-tailed P value? | Two-tailed |
| Sum of ranks in column C,D | 42 , 63 |
| Mann-Whitney U | 14 |
| Difference between medians | |
| Median of column C | 115,1, n=7 |
| Median of column D | 167,0, n=7 |
| Difference: Actual | 51,92 |
| Difference: Hodges-Lehmann | 52,45 |

**SAL-RT vs SAL-WARM**

| | |
|---|---|
| Column F | SAL-WARM |
| vs. | vs, |
| Column E | SAL-RT |
| Mann Whitney test | |
| P value | 0,535 |
| Exact or approximate P value? | Exact |
| P value summary | ns |
| Significantly different (P < 0.05)? | No |
| One- or two-tailed P value? | Two-tailed |
| Sum of ranks in column E,F | 58 , 47 |
| Mann-Whitney U | 19 |
| Difference between medians | |
| Median of column E | 109,7, n=7 |
| Median of column F | 92,04, n=7 |
| Difference: Actual | -17,61 |
| Difference: Hodges-Lehmann | -12,81 |

**pp70s6k**

**SAL-RT vs AMI-RT vs AMI-WARM**

| | |
|---|---|
| Kruskal-Wallis test | |
| P value | 0,0554 |
| Exact or approximate P value? | Exact |
| P value summary | ns |
| Do the medians vary signif. (P < 0.05) | No |
| Number of groups | 3 |
| Kruskal-Wallis statistic | 5,618 |
| Data summary | |

| | |
|---|---|
| Number of treatments (columns) | 3 |
| Number of values (total) | 21 |

| Dunn's multiple comparisons test | Mean rank | Significant | Summary | Adjusted P | A-? | |
|---|---|---|---|---|---|---|
| SalRT vs. AMI-RT | -7,857 | Yes | * | 0,0357 | B | AMI-RT |
| SalRT vs. AMI-W | -4,143 | No | ns | 0,4232 | C | AMI-W |

| Test details | Mean rank | Mean rank | Mean rank diff, | n1 | n2 | Z |
|---|---|---|---|---|---|---|
| SalRT vs. AMI-RT | 7 | 14,86 | -7,857 | 7 | 7 | 2,369 |
| SalRT vs. AMI-W | 7 | 11,14 | -4,143 | 7 | 7 | 1,249 |

**SAL-RT vs SAL-W**

| | |
|---|---|
| Column B | SAL-WARM |
| vs. | vs, |
| Column A | SAL-RT |
| Mann Whitney test | |
| P value | 0,1282 |
| Exact or approximate P value? | Exact |
| P value summary | ns |
| Significantly different (P < 0.05)? | No |
| One- or two-tailed P value? | Two-tailed |
| Sum of ranks in column A,B | 65 , 40 |
| Mann-Whitney U | 12 |
| Difference between medians | |
| Median of column A | 102,7, n=7 |
| Median of column B | 83,39, n=7 |
| Difference: Actual | -19,26 |
| Difference: Hodges-Lehmann | -15,92 |

**pGSK3beta**

**SAL-RT vs AMI-RT**

| | |
|---|---|
| Column B | AMI-RT |
| vs. | vs, |
| Column A | SAL-RT |
| Mann Whitney test | |
| P value | 0,0973 |
| Exact or approximate P value? | Exact |
| P value summary | ns |
| Significantly different (P < 0.05)? | No |
| One- or two-tailed P value? | Two-tailed |
| Sum of ranks in column A,B | 39 , 66 |
| Mann-Whitney U | 11 |
| Difference between medians | |
| Median of column A | 102,7, n=7 |
| Median of column B | 150,1, n=7 |
| Difference: Actual | 47,4 |
| Difference: Hodges-Lehmann | 60,25 |

**SAL-RT vs AMI-W**

| | |
|---|---|
| Column D | AMI-W |
| vs. | vs, |
| Column C | SAL-RT |
| Mann Whitney test | |
| P value | 0,2086 |
| Exact or approximate P value? | Exact |
| P value summary | ns |
| Significantly different (P < 0.05)? | No |
| One- or two-tailed P value? | Two-tailed |
| Sum of ranks in column C,D | 42 , 63 |
| Mann-Whitney U | 14 |
| Difference between medians | |
| Median of column C | 79,45, n=7 |
| Median of column D | 135,9, n=7 |
| Difference: Actual | 56,48 |
| Difference: Hodges-Lehmann | 44,7 |

**SAL-RT vs SAL-WARM**

| | |
|---|---|
| Column F | SAL-WARM |
| vs. | vs, |
| Column E | SAL-RT |
| Mann Whitney test | |
| P value | 0,3829 |
| Exact or approximate P value? | Exact |
| P value summary | ns |
| Significantly different (P < 0.05)? | No |
| One- or two-tailed P value? | Two-tailed |
| Sum of ranks in column E,F | 60 , 45 |
| Mann-Whitney U | 17 |
| Difference between medians | |
| Median of column E | 72,85, n=7 |
| Median of column F | 39,50, n=7 |
| Difference: Actual | -33,35 |
| Difference: Hodges-Lehmann | -35,2 |

**Body Temperature**

| | |
|---|---|
| Kruskal-Wallis test | |
| P value | 0,0006 |
| Exact or approximate P value? | Approximate |
| P value summary | *** |
| Do the medians vary signif. (P < 0.05) | Yes |
| Number of groups | 4 |
| Kruskal-Wallis statistic | 17,44 |

| Dunn's multiple comparisons test | Mean rank | Significant | Summary | Adjusted P Value | |
|---|---|---|---|---|---|
| SAL-W vs. AMI-W | 6,17 | No | ns | >0,9999 | A-B |
| SAL-W vs. SAL-RT | 3,482 | No | ns | >0,9999 | A-C |
| SAL-W vs. AMI-RT | 18,36 | Yes | *** | 0,0006 | A-D |
| AMI-W vs. SAL-RT | -2,688 | No | ns | >0,9999 | B-C |
| AMI-W vs. AMI-RT | 12,19 | Yes | * | 0,0446 | B-D |
| SAL-RT vs. AMI-RT | 14,88 | Yes | ** | 0,0065 | C-D |

| Test details | Mean rank | Mean rank | Mean rank diff, | n1 | n2 | Z |
|---|---|---|---|---|---|---|
| SAL-W vs. AMI-W | 22,36 | 16,19 | 6,17 | 7 | 8 | 1,355 |
| SAL-W vs. SAL-RT | 22,36 | 18,88 | 3,482 | 7 | 8 | 0,7647 |
| SAL-W vs. AMI-RT | 22,36 | 4 | 18,36 | 7 | 7 | 3,903 |
| AMI-W vs. SAL-RT | 16,19 | 18,88 | -2,688 | 8 | 8 | 0,6109 |
| AMI-W vs. AMI-RT | 16,19 | 4 | 12,19 | 8 | 7 | 2,676 |
| SAL-RT vs. AMI-RT | 18,88 | 4 | 14,88 | 8 | 7 | 3,267 |

**B**

**pTrkB**

| | |
|---|---|
| Kruskal-Wallis test | |
| P value | 0,0086 |
| Exact or approximate P value? | Approximate |
| P value summary | ** |
| Do the medians vary signif. (P < 0.05) | Yes |
| Number of groups | 4 |
| Kruskal-Wallis statistic | 11,67 |
| Data summary | |
| Number of treatments (columns) | 4 |
| Number of values (total) | 20 |

| Dunn's test | Mean rank | Significant | Summary | Individual P Value | |
|---|---|---|---|---|---|
| VVS vs. VVA | -8,4 | Yes | * | 0,0248 | A-B |
| VVS vs. MMS | 0 | No | ns | >0,9999 | A-C |
| VVS vs. MMA | -9,6 | Yes | * | 0,0103 | A-D |
| VVA vs. MMS | 8,4 | Yes | * | 0,0248 | B-C |
| VVA vs. MMA | -1,2 | No | ns | 0,7484 | B-D |
| MMS vs. MMA | -9,6 | Yes | * | 0,0103 | C-D |

| Test details | Mean rank | Mean rank | Mean rank diff, | n1 | n2 | Z |
|---|---|---|---|---|---|---|
| VVS vs. VVA | 6 | 14,4 | -8,4 | 5 | 5 | 2,245 |
| VVS vs. MMS | 6 | 6 | 0 | 5 | 5 | 0 |

| | | | | | | |
|---|---|---|---|---|---|---|
| VVS vs. MMA | 6 | 15,6 | -9,6 | 5 | 5 | 2,566 |
| VVA vs. MMS | 14,4 | 6 | 8,4 | 5 | 5 | 2,245 |
| VVA vs. MMA | 14,4 | 15,6 | -1,2 | 5 | 5 | 0,3207 |
| MMS vs. MMA | 6 | 15,6 | -9,6 | 5 | 5 | 2,566 |

**pGSK3beta**

| | |
|---|---|
| Kruskal-Wallis test | |
| P value | 0,0071 |
| Exact or approximate P value? | Approximate |
| P value summary | ** |
| Do the medians vary signif. (P < 0.05)' | Yes |
| Number of groups | 4 |
| Kruskal-Wallis statistic | 12,1 |
| Data summary | |
| Number of treatments (columns) | 4 |
| Number of values (total) | 20 |

| Dunn's test | Mean rank | Significant' | Summary | Adjusted P Value | |
|---|---|---|---|---|---|
| VVS vs. VVA | -6,8 | No | ns | 0,415 | A-B |
| VVS vs. MMS | 0,2 | No | ns | >0,9999 | A-C |
| VVS vs. MMA | -10,6 | Yes | * | 0,0277 | A-D |
| VVA vs. MMS | 7 | No | ns | 0,3682 | B-C |
| VVA vs. MMA | -3,8 | No | ns | >0,9999 | B-D |
| MMS vs. MMA | -10,8 | Yes | * | 0,0234 | C-D |

| Test details | Mean rank | Mean rank | Mean rank diff, | n1 | n2 | Z |
|---|---|---|---|---|---|---|
| VVS vs. VVA | 6,2 | 13 | -6,8 | 5 | 5 | 1,817 |
| VVS vs. MMS | 6,2 | 6 | 0,2 | 5 | 5 | 0,05345 |
| VVS vs. MMA | 6,2 | 16,8 | -10,6 | 5 | 5 | 2,833 |
| VVA vs. MMS | 13 | 6 | 7 | 5 | 5 | 1,871 |
| VVA vs. MMA | 13 | 16,8 | -3,8 | 5 | 5 | 1,016 |
| MMS vs. MMA | 6 | 16,8 | -10,8 | 5 | 5 | 2,886 |

**Body Temperature**

| | |
|---|---|
| Kruskal-Wallis test | |
| P value | 0,0023 |
| Exact or approximate P value? | Approximate |
| P value summary | ** |
| Do the medians vary signif. (P < 0.05)' | Yes |
| Number of groups | 4 |
| Kruskal-Wallis statistic | 14,52 |
| Data summary | |
| Number of treatments (columns) | 4 |
| Number of values (total) | 19 |

| Dunn's test | Mean rank | Significant' | Summary | Adjusted P Value | |
|---|---|---|---|---|---|
| VVS vs. VVA | 10,7 | Yes | * | 0,0155 | A-B |
| VVS vs. MMS | 3,6 | No | ns | >0,9999 | A-C |
| VVS vs. MMA | 11,5 | Yes | ** | 0,0072 | A-D |
| VVA vs. MMS | -7,1 | No | ns | 0,3566 | B-C |
| VVA vs. MMA | 0,8 | No | ns | >0,9999 | B-D |
| MMS vs. MMA | 7,9 | No | ns | 0,2158 | C-D |

| Test details | Mean rank | Mean rank | Mean rank diff, | n1 | n2 | Z |
|---|---|---|---|---|---|---|
| VVS vs. VVA | 16,6 | 5,9 | 10,7 | 5 | 5 | 3,013 |
| VVS vs. MMS | 16,6 | 13 | 3,6 | 5 | 4 | 0,9558 |
| VVS vs. MMA | 16,6 | 5,1 | 11,5 | 5 | 5 | 3,238 |
| VVA vs. MMS | 5,9 | 13 | -7,1 | 5 | 4 | 1,885 |
| VVA vs. MMA | 5,9 | 5,1 | 0,8 | 5 | 5 | 0,2253 |
| MMS vs. MMA | 13 | 5,1 | 7,9 | 4 | 5 | 2,097 |

**Figure 6. Modulation of TrkB signaling and antidepressant effects**

**B Corticosterone (CORT) model, nitrous oxide (N2O) + warming**

| **Treatment** | **Dose** | **Warming** | **n** |
|---|---|---|---|
| No CORT | | | 10 |
| CORT + SHAM | | | 8 |
| CORT + SHAM | | 36 | 8 |
| CORT + N2O | 65 % | | 8 |
| CORT + N2O | 65 % | 36 | 9 |

**SPT1**

| | |
|---|---|
| Kruskal-Wallis test | |
| P value | <0,0001 |
| Exact or approximate P value? | Approximate |
| P value summary | **** |
| Do the medians vary signif. (P < 0.05)? | Yes |
| Number of groups | 5 |
| Kruskal-Wallis statistic | 25,79 |

| | |
|---|---|
| Data summary | |
| Number of treatments (columns) | 5 |
| Number of values (total) | 43 |

| Dunn's multiple comparisons test | Mean rank ( | Significant? | Summary | Adjusted P Value | |
|---|---|---|---|---|---|
| NO CORT vs. CORT sham | 23,93 | Yes | *** | 0,0006 | A-B |
| NO CORT vs. CORT sham warm | 14,61 | No | ns | 0,1413 | A-C |
| NO CORT vs. CORT N2O | 9,738 | No | ns | >0,9999 | A-D |
| NO CORT vs. CORT N2O warm | 25,41 | Yes | *** | 0,0001 | A-E |
| CORT sham vs. CORT sham warm | -9,313 | No | ns | >0,9999 | B-C |
| CORT sham vs. CORT N2O | -14,19 | No | ns | 0,238 | B-D |
| CORT sham vs. CORT N2O warm | 1,486 | No | ns | >0,9999 | B-E |
| CORT sham warm vs. CORT N2O | -4,875 | No | ns | >0,9999 | C-D |
| CORT sham warm vs. CORT N2O warm | 10,8 | No | ns | 0,7668 | C-E |
| CORT N2O vs. CORT N2O warm | 15,67 | No | ns | 0,1019 | D-E |

| Test details | Mean rank : | Mean rank 2 | Mean rank ( | n1 | n2 | Z |
|---|---|---|---|---|---|---|
| NO CORT vs. CORT sham | 36,3 | 12,38 | 23,93 | 10 | 8 | 4,018 |
| NO CORT vs. CORT sham warm | 36,3 | 21,69 | 14,61 | 10 | 8 | 2,454 |
| NO CORT vs. CORT N2O | 36,3 | 26,56 | 9,738 | 10 | 8 | 1,635 |
| NO CORT vs. CORT N2O warm | 36,3 | 10,89 | 25,41 | 10 | 9 | 4,406 |
| CORT sham vs. CORT sham warm | 12,38 | 21,69 | -9,313 | 8 | 8 | 1,484 |
| CORT sham vs. CORT N2O | 12,38 | 26,56 | -14,19 | 8 | 8 | 2,26 |
| CORT sham vs. CORT N2O warm | 12,38 | 10,89 | 1,486 | 8 | 9 | 0,2436 |
| CORT sham warm vs. CORT N2O | 21,69 | 26,56 | -4,875 | 8 | 8 | 0,7767 |
| CORT sham warm vs. CORT N2O warm | 21,69 | 10,89 | 10,8 | 8 | 9 | 1,77 |
| CORT N2O vs. CORT N2O warm | 26,56 | 10,89 | 15,67 | 8 | 9 | 2,569 |

**SPT2**

| | |
|---|---|
| Kruskal-Wallis test | |
| P value | 0,0004 |
| Exact or approximate P value? | Approximate |
| P value summary | *** |
| Do the medians vary signif. (P < 0.05)? | Yes |
| Number of groups | 5 |
| Kruskal-Wallis statistic | 20,7 |

| | |
|---|---|
| Data summary | |
| Number of treatments (columns) | 5 |
| Number of values (total) | 44 |

| Dunn's multiple comparisons test | Mean rank ( | Significant? | Summary | Adjusted P Value |
|---|---|---|---|---|

| | | | | | |
|---|---|---|---|---|---|
| NO CORT vs. CORT sham | 19,16 | Yes | * | 0,0166 | A-B |
| NO CORT vs. CORT sham warm | 12,48 | No | ns | 0,4061 | A-C |
| NO CORT vs. CORT N2O | 20,73 | Yes | ** | 0,0067 | A-D |
| NO CORT vs. CORT N2O warm | 23,45 | Yes | *** | 0,0004 | A-E |
| CORT sham vs. CORT sham warm | -6,688 | No | ns | >0,9999 | B-C |
| CORT sham vs. CORT N2O | 1,563 | No | ns | >0,9999 | B-D |
| CORT sham vs. CORT N2O warm | 4,288 | No | ns | >0,9999 | B-E |
| CORT sham warm vs. CORT N2O | 8,25 | No | ns | >0,9999 | C-D |
| CORT sham warm vs. CORT N2O warm | 10,98 | No | ns | 0,7166 | C-E |
| CORT N2O vs. CORT N2O warm | 2,725 | No | ns | >0,9999 | D-E |

| Test details | Mean rank : | Mean rank 2 | Mean rank ( | n1 | n2 | Z |
|---|---|---|---|---|---|---|
| NO CORT vs. CORT sham | 37,35 | 18,19 | 19,16 | 10 | 8 | 3,145 |
| NO CORT vs. CORT sham warm | 37,35 | 24,88 | 12,48 | 10 | 8 | 2,047 |
| NO CORT vs. CORT N2O | 37,35 | 16,63 | 20,73 | 10 | 8 | 3,402 |
| NO CORT vs. CORT N2O warm | 37,35 | 13,9 | 23,45 | 10 | 10 | 4,082 |
| CORT sham vs. CORT sham warm | 18,19 | 24,88 | -6,688 | 8 | 8 | 1,041 |
| CORT sham vs. CORT N2O | 18,19 | 16,63 | 1,563 | 8 | 8 | 0,2433 |
| CORT sham vs. CORT N2O warm | 18,19 | 13,9 | 4,288 | 8 | 10 | 0,7037 |
| CORT sham warm vs. CORT N2O | 24,88 | 16,63 | 8,25 | 8 | 8 | 1,285 |
| CORT sham warm vs. CORT N2O warm | 24,88 | 13,9 | 10,98 | 8 | 10 | 1,801 |
| CORT N2O vs. CORT N2O warm | 16,63 | 13,9 | 2,725 | 8 | 10 | 0,4472 |
